# Two independent origins of XY sex chromosomes in *Asparagus*

**DOI:** 10.1101/2025.09.05.674532

**Authors:** Philip C. Bentz, Sarah B. Carey, Francesco Mercati, Haley Hale, Valentina Ricciardi, Francesco Sunseri, Alex Harkess, Jim Leebens-Mack

## Abstract

The relatively young and repeated evolutionary origins of dioecy (separate sexes) in flowering plants enable investigation of molecular dynamics occurring at the earliest stages of sex chromosome evolution. With two independently young origins of dioecy in the genus, *Asparagus* is a model taxon for studying genetic sex-determination and sex chromosome evolution. Dioecy first evolved in *Asparagus* ∼3-4 million years ago (Ma) in the ancestor of a now widespread Eurasian clade that includes garden asparagus (*Asparagus officinalis*), while the second origin occurred in a smaller, geographically restricted, Mediterranean Basin clade including *Asparagus horridus*. The XY sex chromosomes and sex-determination genes in garden asparagus have been well characterized, but the genetics underlying dioecy in the Mediterranean Basin clade are unknown. We generated new haplotype-resolved reference genomes for garden asparagus and *A. horridus*, to elucidate the sex chromosomes of *A. horridus* and explore how dioecy evolved between these two closely related lineages. Analysis of the *A. horridus* genome revealed an independently evolved XY system derived from different ancestral autosomes (chromosome 3) with different sex-determining genes than documented for garden asparagus (on chromosome 1). We estimate that proto-XY chromosomes evolved around 1-2 Ma in the Mediterranean Basin clade, following an ∼2.1-megabase inversion between the ancestral pair. Recombination suppression and LTR retrotransposon accumulation drove the establishment and expansion of the Y-linked sex-determination region (Y-SDR) that now reaches ∼9.6-megabases in *A. horridus*. The new garden asparagus genome revealed a Y-SDR that spans ∼1.9-megabases with ten hemizygous genes. Our results evoke hemizygosity as the most probable mechanism responsible for the origin of proto-XY recombination suppression in the Eurasian clade, and that neofunctionalization of one duplicated gene (*SOFF*) drove the origin of dioecy. These findings support previous inference based on phylogeographic analysis revealing two recent origins of dioecy in *Asparagus.* Moreover, this work implicates alternative molecular mechanisms for two separate shifts to dioecy in a model taxon important for investigating young sex chromosome evolution.

**SIGNIFICANCE STATEMENT:** Flowering plants with separate sexes are ideal systems for investigating genome dynamics underlying the earliest stages of sex chromosome evolution across the tree of life. We use *Asparagus* as a model to better understand early sex chromosome formation more generally, by investigating how different XY sex chromosomes evolved between two young, closely related clades. Genomic comparisons of garden asparagus and *Asparagus horridus* (wild related species) revealed distinct evolutionary origins of XY-chromosomes with different sex-determination mechanisms. Whereas the garden asparagus Y-chromosome originally evolved around 3-4 million years ago (Ma), following a small segmental duplication, the Y-chromosome in *Asparagus horridus* evolved more recently (∼1-2 Ma) following a large structural inversion between a different chromosome pair. Interestingly, both evolutionary transitions from hermaphroditism to separate sexes occurred as ancestors of garden asparagus and *Asparagus horridus* independently dispersed northward out of southern Africa.

## INTRODUCTION

Separate sexes and sex chromosomes have evolved many times across the tree of life. Sex chromosomes exhibit unique evolutionary innovations relative to autosomes, including regions of suppressed recombination, size differences between X and Y (or Z/W) chromosomes, and the evolution of sex-specific gene content and expression patterns. Across the angiosperms, most extant species produce perfect flowers that produce male and female gametophytes (pollen and ovules, respectively), but separate sexes (i.e., dioecy, or unisexual flowers on different plants) have evolved in less than 10% of angiosperm species (1). In contrast to ancient sex-determination systems, including the XY system in placental mammals, dioecy and sex chromosomes have evolved independently and more recently across many angiosperm clades (2). The repeated and recent evolution of dioecy across the angiosperms offers an opportunity to investigate the earliest stages of sex chromosome evolution (3, 4). However, it is not yet clear whether the origin and evolution of dioecy occurs through a common set of genomic and molecular mechanisms, or rather, there are a myriad evolutionary paths for the transition from autosomes in bisexual species to sex chromosomes in dioecious species (5–7). By studying sex chromosomes of different evolutionary ages, we may begin to understand the molecular mechanisms and ecology driving the evolution of separate sexes more broadly. Investigations of independently evolving sex chromosomes among closely related dioecious species may be especially informative for understanding the origin and evolution of sex chromosomes.

The genus *Asparagus* Tourn. ex L. (Asparagaceae) is an important model system for studying genetic sex-determination and sex chromosome dynamics in plants. Investigations of the genetic basis of sex-determination in garden asparagus (*Asparagus officinalis* L.) characterized Y-specific genes responsible for the suppression of pistil (female) development and completion of pollen (male) development (8–11). However, little is known regarding sex-determination and sex chromosomes in the other 50+ dioecious species of *Asparagus,* partly due to historical uncertainty surrounding sexual modes and species relationships across the genus. To address these limitations, we recently reviewed all sexual systems reported in the genus (12) and released an updated phylogeny based on 1,726 nuclear genes (13) and robust species sampling (>150 spp.) (14). Phylogeographic analysis supports two independent origins of dioecy associated with separate long-distance dispersal events, both occurring around 2-4 million years ago (Ma), in a widespread Eurasian clade and a smaller Mediterranean Basin clade (12, 14, 15) (**Fig. 1**). Two recently-derived dioecious systems within *Asparagus* makes it an ideal system for testing for common themes in molecular dynamics contributing to dioecy and sex chromosome evolution.

**Figure 1.**
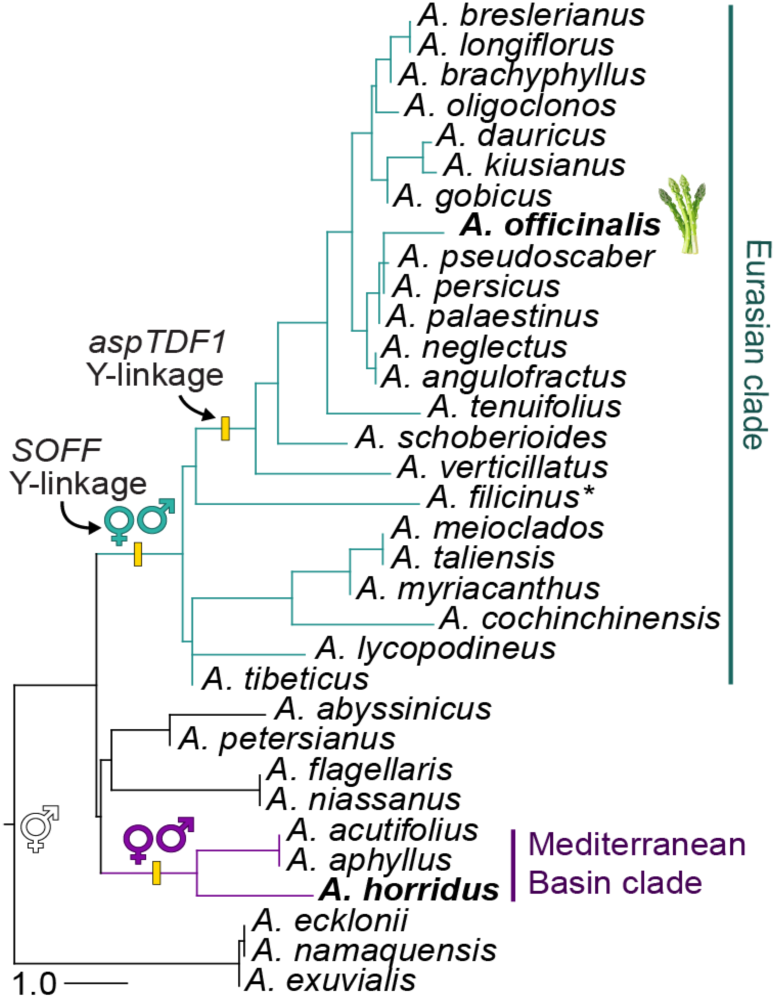
Transitions from an ancestral bisexual state to lineages with separate sexes—dioecy evolved twice in the genus *Asparagus*. The dioecious Eurasian clade includes garden asparagus (*A. officinalis*) and over 50 additional species that are widespread across Eurasia. The dioecious Mediterranean Basin clade, with *A. horridus*, is much less speciose (∼3-4 spp.) and geographically restricted to regions around the Mediterranean Sea (12). Two genes control sex-determination in garden asparagus, a female suppressor (*SOFF*) and male promoter (*aspTDF1*) (8, 9). Previous work showed that *aspTDF1* is male-specific in only a subset of Eurasian clade lineages, and no Mediterranean Basin clade lineages (9, 11), whereas *SOFF* is the most ancestral, male-specific, sex-determination gene and is associated with the origin of dioecy in the Eurasian clade (9). The updated *Asparagus* phylogeny (shown here, adapted from Bentz et al. (14)) highlights the stepwise and separate evolutions of extant sex-determining genes in the Eurasian clade, with *aspTDF1* evolving Y-linkage following *SOFF* and the origin of dioecy in the clade. *sex-linkage of *aspTDF1* not tested in species.

In garden asparagus (Eurasian clade), the presence of an ∼1 Mb nonrecombining, Y-specific region controls sex (hereafter, “sex-determination region”, or “SDR”), which was identified in the first reference genome of a double-haploid YY male (9). The Y-linked SDR (Y-SDR) in the YY garden asparagus reference included 13 annotated genes, two of which were shown to sufficiently control sex in experimental and spontaneous mutant genotypes: *SUPPRESSOR OF FEMALE FUNCTION* (*SOFF*), a *DUF247* gene that suppresses pistil development; and *TAPETAL DEVELOPMENT and FUNCTION 1* (*aspTDF1*), an *R2R3*-type *MYB* transcription factor and male-promoter gene influencing tapetal and pollen development (9–11). The presence of two sex-determining genes in garden asparagus supports a two gene hypothesis for the evolution of sex chromosomes with at least two linked mutations affecting female and male fertility, respectively (16, 17), rather than a single master-switch sex-determining gene as described in the Salicaceae (18). Previous work shows that a *SOFF* ortholog is conserved as male-specific (or Y-linked) across the entire Eurasian dioecy clade (9) (**Fig. 1**). However, PCR assays revealed that whereas *aspTDF1* is male-specific in garden asparagus, and its closest relatives, it is autosomal in other dioecious *Asparagus* species including *A. acutifolius, A. horridus,* and *A. cochinchinensis* (11). *Asparagus cochinchinensis* falls within a subclade that split from the subclade with garden asparagus early in the evolution of the Eurasian dioecious clade, while *A. acutifolius,* and *A. horridus* are species in the Mediterranean Basin clade (**Fig. 1**) (14); suggesting that *aspTDF1* evolved Y-linkage following the origin of dioecy within the Eurasian clade, and that the Mediterranean Basin clade may have independently evolved a distinct sex-determination system (**Fig. 1**).

In this study, we explore the origins of recombination suppression between ancestral autosomes that led to independent sex chromosome formation in the Mediterranean Basin and Eurasian dioecious clades of *Asparagus.* Specifically we leverage an updated phylogenomic framework (14) by comparing new haplotype-resolved, reference-grade genomes for taxa from the two dioecious clades: *A. horridus* (Mediterranean Basin) and *A. officinalis* (Eurasian). Both new genomes are of diploid male genotypes (2n = 2x = 20) (19). This study is the first to present a genome assembly for *A. horridus* and identify the sex chromosomes for the species. Our findings advance understanding of how dioecy can evolve between two closely related clades with unique sex chromosomes, expanding the utility of *Asparagus* as a model for the study sex chromosome evolution more broadly.

## RESULTS AND DISCUSSION

### Asparagus horridus genome and sex chromosomes

Here we present the first reference genome for *A. horridus* (**Fig. 2a**) and report an XY sex chromosome system for the species. The *A. horridus* genome assembly size was in-line with flow cytometry-based estimates (19), totalling ∼1.01-1.03 Gb per haplotype (**Table S1**). Presence of a larger Y chromosome (∼107 Mb) in haplotype 1, compared to the X (∼99 Mb), largely explains the size difference between the two haplotype assemblies (**Table S2**).

**Figure 2.**
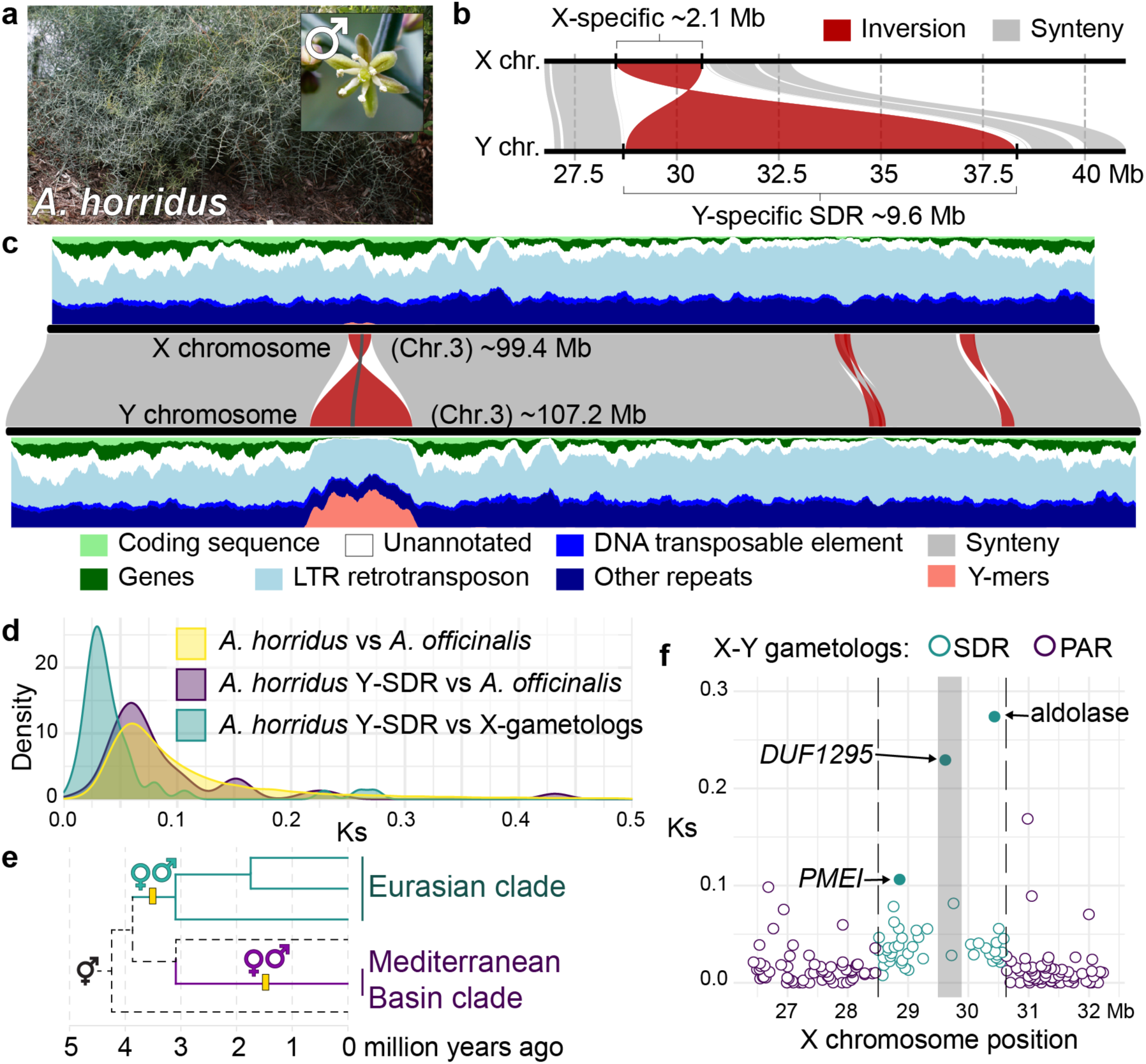
XY sex chromosomes evolved from chromosome 3 (ancestral autosomes) in the Mediterranean Basin dioecious clade with *Asparagus horridus*. **a)** *A. horridus* genome-line male plant (pb32m) producing species-typical staminate flowers with undeveloped pistil remnants. **b)** One large inversion marks the boundaries of the nonrecombining, Y-specific sex-determination region (Y-SDR) and inverted X-specific region. **c)** X-Y haplotype alignment (middle track) and structural annotation densities (X=top; Y=bottom) show significant increases in Y-mers (male-specific *k*-mers identified between 8 males – 7 females) and LTR retrotransposons within the Y-SDR corresponding to the large inversion boundaries. Dark grey region within the Y-SDR marks a nested inversion with three genes (highlighted in panel f). **d)** Synonymous substitutions (Ks) measured in three tests suggest that the *A. horridus* Y-SDR (teal curve) is younger than the divergence between *A. horridus* and *A. officinalis* (purple and yellow curves). Yellow = all *A. horridus* and *A. officinalis* orthologs. Purple = *A. horridus* Y-SDR genes and orthologs in *A. officinalis*. Teal = *A. horridus* Y-SDR genes and X-linked homeologs (gametologs). **e)** Dioecy evolved ∼1.13-1.81 Ma in the Mediterranean Basin clade, around one to two million years later than in the Eurasian clade. Dotted branches represent hermaphroditic lineages and the clade’s ancestral state (12, 14, 15). **f)** Compared to the pseudo-autosomal region (PAR), synonymous substitutions were consistently elevated across Y-SDR gametologs aside from three outlier genes (filled circles).

Assembly and annotation quality metrics for the *A. horridus* genome haplotypes were above standard reference thresholds (**Table S1, Fig. S1**). Fewer genes were predicted in *A. horridus* (31,194 to 31, 235 genes per haplotype), compared to the new *A. officinalis* genome annotation (**Table S1**), but were generally in-line with *A. officinalis* gene counts based on histone modification (ChIP-seq) data (20). According to pseudo-chromosome alignments and Y-mer (male-specific *k*-mers) mapping we found that the chromosome 3 pair correspond to the *A. horridus* XY chromosomes (**Figs. 2b-c, S2**). We identified an ∼9.66 Mb Y-SDR for *A. horridus,* corresponding to a single inversion distinguishing the pseudo-autosomal (PAR) boundaries between the X and Y chromosomes (**Fig. 2b**). The PAR-SDR boundaries were identified based on Y-mer mapping and a lower ratio of female:male read mapping depth/coverage (**Fig. S3**). A small inversion was found nested within the Y-SDR (**Figs. 2c, S4**), but since only three genes exist in that region, tests to assess whether that structural change occurred before or after the larger inversion lack statistical power. The Y-SDR in *A. horridus* is relatively gene poor (78-122 total non-TE gene predictions, see **Table S3** for functional annotations) and highly enriched in TE content, especially of *Ty3-* and *Ty1-*type LTR retrotransposons (**Fig. 2c, Table S2**). LTRs, and repeats more generally, are thought to lead to artificially inflated gene model estimates, especially of single-exon genes (mono-exonic) (21). In the *A. horridus* Y-SDR, 62 of the predicted genes were mono-exonic (**Tables S2**) and ∼71% of the mono-exonic genes lacked orthologs in other species, and therefore may include erroneous TE-derived gene models. The *A. horridus* Y-SDR corresponds to a collinear, but inverted, region on the X that spans ∼2.12 Mb (**Fig. 2b**) with 77 gene models (∼23% mono-exonic) and considerably lower repeat content than the Y-SDR (**Table S2**). Interestingly, nearly all genes annotated in the nonrecombining portion of the X chromosome have intact gametologs in the Y-SDR, suggesting limited or no gene degeneration in the region. Finally, and perhaps most importantly, no shared homologs were found between the *A. horridus* and *A. officinalis* Y-SDRs, supporting independent origins of genetic sex-determination and XY chromosomes in the two dioecious clades in the genus *Asparagus*.

### Origin of XY sex chromosomes in the Mediterranean Basin clade

Sex chromosomes evolved only once in the Mediterranean Basin clade, as indicated by the conservation of male-specific sequences shared between *A. horridus* and *A. acutifolius* (**Fig. S3**). The almost 10 Mb Y-SDR in *A. horridus* is syntenic with ∼2.1 Mb on chromosome 3 in *A. officinalis*, *A. setaceus*, and the *A. horridus* X. This suggests that proto-XYs in the Mediterranean Basin clade evolved following an ∼2.1 Mb inversion between the ancestral chromosome 3 pair, inhibiting recombination throughout the region and allowing for X-Y divergence. Expansion of the nonrecombining Y-SDR—as a consequence of TE accumulation—is thought to be a dominant process contributing to disparate sizes between the sex chromosomes of some lineages (22). Compared to the X-limited region, the Y-SDR in *A. horridus* exhibits a roughly four times greater ratio of repeats (**Table S2**) supporting TE enrichment (**Fig. 2c**) as a major driver of its expansion over time.

Little or no degeneration in the nonrecombining Y-SDR, as indicated by intact genic synteny (syntologs) shared with autosomes from other species, may suggest a young evolutionary age. To estimate the timing of dioecy evolution in the Mediterranean Basin clade relative to the total divergence from Eurasian clade lineages, we measured Ks between X-Y gametologs (teal curve in **Fig. 2d**, 0.034 median), and compared the X-Y Ks distribution with genome-wide Ks values for *A. officinalis–A. horridus* orthologs (yellow curve in **Fig. 2d**, 0.084 median), then *A. horridus* Y-SDR genes vs. *A. officinalis* orthologs (purple curve in **Fig. 2d**, 0.071 median). *Asparagus horridus* and *A. officinalis* diverged from a common ancestor approximately 2.78-3.78 Ma (12). We found that median Ks measurements between the *A. horridus* X-Y gametologs were approximately 47.9% (p-value 1.43e-10) and 40.5% (p-value 4.54e-09) less than the genome-wide and *A. horridus* Y-SDR comparisons with *A. officinalis* orthologs, respectively. Ks differences were not significantly different between the two comparisons with *A. officinalis* orthologs (p-value 0.10), which we used as an experimental control. Based on Ks comparisons and the estimated age of the most recent common ancestor (MRCA) of *A. horridus* and *A. officinalis*, the *A. horridus* X-Y nonrecombining regions began diverging ∼1.33-1.81 or 1.13-1.53 Ma (**Fig. 2e**), which is less than 1 million years after the origin of the Mediterranean Basin crown group (i.e., ∼1.9-2.9 Ma) (12). We posit that a single inversion led to the origin of recombination suppression between proto-XY chromosomes ∼1.13-1.81 Ma in the Mediterranean Basin clade, and rapid accumulation of LTR retrotransposons drove the expansion of the Y-SDR over time.

Candidate genes for sex-determination and sexual antagonism in *A. horridus* Testing for candidate sex-determining genes requires functional validation (e.g., genetic knockouts) and is thus out of the scope of this study, but comparisons of X-Y gene content, expression, and molecular evolutionary analysis can yield focused lists of gene candidates for sex-determination and other sex-specific phenotypes. Of the SDR-linked genes in *A. horridus,* we found that 20 were significantly up-regulated and 7 were down-regulated in male flowers compared to female flowers (across three combined developmental stages, see **Table S4**). Seven of those 27 genes also showed evidence of positive selection in the Y-SDR gametolog compared to their X-linked counterpart and orthologs from other species (**Table S5**).

Two Y-linked pectin methylesterase inhibitor (*PMEI*) genes exhibited interesting patterns linked to sex: one exhibited elevated Ks (Ks 0.11, **Fig. 2f**), positive selection (d_N_/d_S_ 68), and significantly lower expression in male flowers compared to female flowers (log2FC - 3.3) (**Table S5**); while the the other was highly up-regulated (log2FC +5.6) with no signs of positive selection (d_N_/d_S_ 1) (**Tables S6-S7**). Pectin methylesterases (*PMEs*) are posttranscriptionally regulated by *PMEIs* (23) and *PME/PMEI* activity plays important roles in many growth processes, including pollen tube development (24) and fruit ripening regulation (25) in *Arabidopsis thaliana* (hereafter, “Arabidopsis”) and the dioecious kiwifruit (*Actinidia deliciosa*) (26). In pear (*Pyrus bretschneideri*), *PMEIs* have also been shown to regulate pollen tube growth (27). Interestingly, *PMEIs* were also found in the Y-SDR of a dioecious night shade (*Solanum appendiculatum*) (28), thus potentially representing a gene family that commonly neofunctionalizes in male heterogametic (XY) systems, in association with loss of selection pressure to maintain function in fruit development and increased pressure for optimization of male-specific functions. We also found evidence of positive selection and increased expression in male vs. female flowers (log2FC +8.1) of a chalcone synthase (*CHS*) gene (**Table S5**). *CHS* genes are involved in pollen development and show sex-biased expression in several other dioecious systems (29–31). For example, chemically induced male sterility experiments in wheat (*Triticum aestivum*) revealed that *CHS* expression decreased in male-sterile plants (32), suggesting a functional role in pollen development and/or viability. Considering our observations of *PME/PMEI* and *CHS* genes in other plant species, and results from this study, Y-linked homologs in *A. horridus* may exhibit additionally derived, sexually antagonistic functions, potentially promoting male-specific functions rather than directly regulating sex-determination. The remaining Y-SDR genes with evidence of positive selection and sex-biased expression include a Fructose-bisphosphate aldolase (transcriptional activator of glycolytic enzymes) (Ks 0.27, **Fig. 2f**), Remorin C-terminal domain, Metallophos domain (*DUF4073*), Metallopeptidase domain, and an unannotated gene model (**Table S5**).

The other inferred Y-SDR genes under positive selection include a 3-oxo-5-alpha-steroid 4-dehydrogenase (*DUF1295*) (Ks 0.23, **Fig. 2f**), an unannotated gene, *APETALA2* (*AP2*)*, SLOW WALKER2* (*SWA2*) (**Table S5**). The Y-linked *AP2* and *SWA2* are evolving under strong selection pressures (d_N_/d_S_ 121 and 111, **Table S5**) and are especially interesting due to their functions in other flowering plants. *AP2* is an ethylene-responsive transcription factor and A-class homeotic gene in the ABC model of flower development in Arabidopsis (33–35). Down-regulation of an *AP2* homolog in rice (*Oryza sativa*) leads to reduced stamens, fused anthers, additional pistils, lower seed efficacy, and decreased pollen viability and germination altogether suggesting a major role in male function (36). A distantly related *AP2* homolog was also found in the *A. officinalis* Y-SDR, but was not specifically implicated in sex-determination for the species (9). *SWA2* is required for coordinated cell cycle progression during female gametophyte and pollen development in Arabidopsis (37, 38). Mutant *swa2* genotypes in Arabidopsis exhibit arrested female gametophyte development and aborted ovules (37, 38).

Pollen cell cycles were also disrupted in Arabidopsis *swa2* mutants—though to a lesser extent compared to the impaired development of female gametophytes—which led to defective pollen development in a small percentage of mutants (38). Further, differential expression analysis among sterile vs. fertile ovules in *Pinus tabuliformis* revealed significantly lower expression of an *SWA2* homolog in sterile ovules compared to the latter, suggesting a conserved functional role in ovule development (39). Considering the broadly conserved role of *SWA2* in female function, and the apparent late abortion of pistil development in male flowers of *A. horridus* (see flower with vestigial pistil development in **Fig. 2a**) this gene may be evolving different functions between the sexes, as indicated by an elevated d_N_/d_S_ ratio. No signs of differential expression of *SWA2* were observed between the sexes, but expression comparisons at additional and distinct developmental stages may reveal sex-specific patterns not detected here.

### Asparagus officinalis genome and sex chromosomes

Publication of a YY double-haploid genome, for *A. officinalis,* in 2017 (9), and an XX double haploid genome in 2020 (8), along with analyses of experimental mutants in both studies, documented a Y-linked two-gene sex-determination system for the species. Recent improvements in genome sequencing and assembly technologies have enabled generation of more accurate and contiguous, diploid-phased reference genomes. The *A. officinalis* genome presented here meets standard quality metrics for reference assemblies and annotations (**Table S1, Fig. S5**) and yielded more complete pseudo-chromosomal assemblies compared to the previous work (9) (**Table S8, Fig. S6**). Total gene content in the new *A. officinalis* reference (34,316 to 34,681 genes per haplotype) was higher than predicted for the earlier *A. officinalis* genomes (8, 9) and the hermaphroditic species *Asparagus setaceus* (40); but are closer to updated gene counts based on ChIP-seq data that revealed as many as 4,640 additional protein coding genes missing from the earlier genome annotation (20).

We used Y-mer mapping and haplotype synteny to delimit the X-Y nonrecombining regions in the new *A. officinalis* diploid-phased assembly (**Figs. 3a-b, S7**). We also used gene trees with X-Y homologs to precisely define PAR-SDR boundaries. Comparison of the two haplotypes revealed a fully hemizygous region spanning ∼1.87 Mb on chromosome 1 of haplotype 2 corresponding to the Y-SDR, and an ∼0.13 Mb X-specific region in haplotype 1 (**Table S2**, **Fig. 3b**). Technological advancements in genome sequencing and assembly explain the different size estimates of the *A. officinalis* sex-limited regions in this study compared to previous work (8, 9), as no scaffolding was required throughout those regions in the new assembly. In the YY double-haploid *A. officinalis* assembly, 6 of the 13 Y-SDR genes were identified on sex-linked contigs that were not anchored to the physical genome map, raising the possibility of misplacement (9). Two of those 6, originally unanchored, gene models were collapsed into a single gene model and 3 others were reassigned to the PAR in our haplotype-resolved assemblies (**Table S9**). Ten total genes (non-TE-associated) were predicted in the updated *A. officinalis* Y-SDR, including male-specific copies of *SOFF* (**Fig. 3c**) and *aspTDF1* (**Fig. 3d**). As expected given the hemizygosity of the Y-SDR, no gametologs were found between the Y- and X-specific regions.

**Figure 3.**
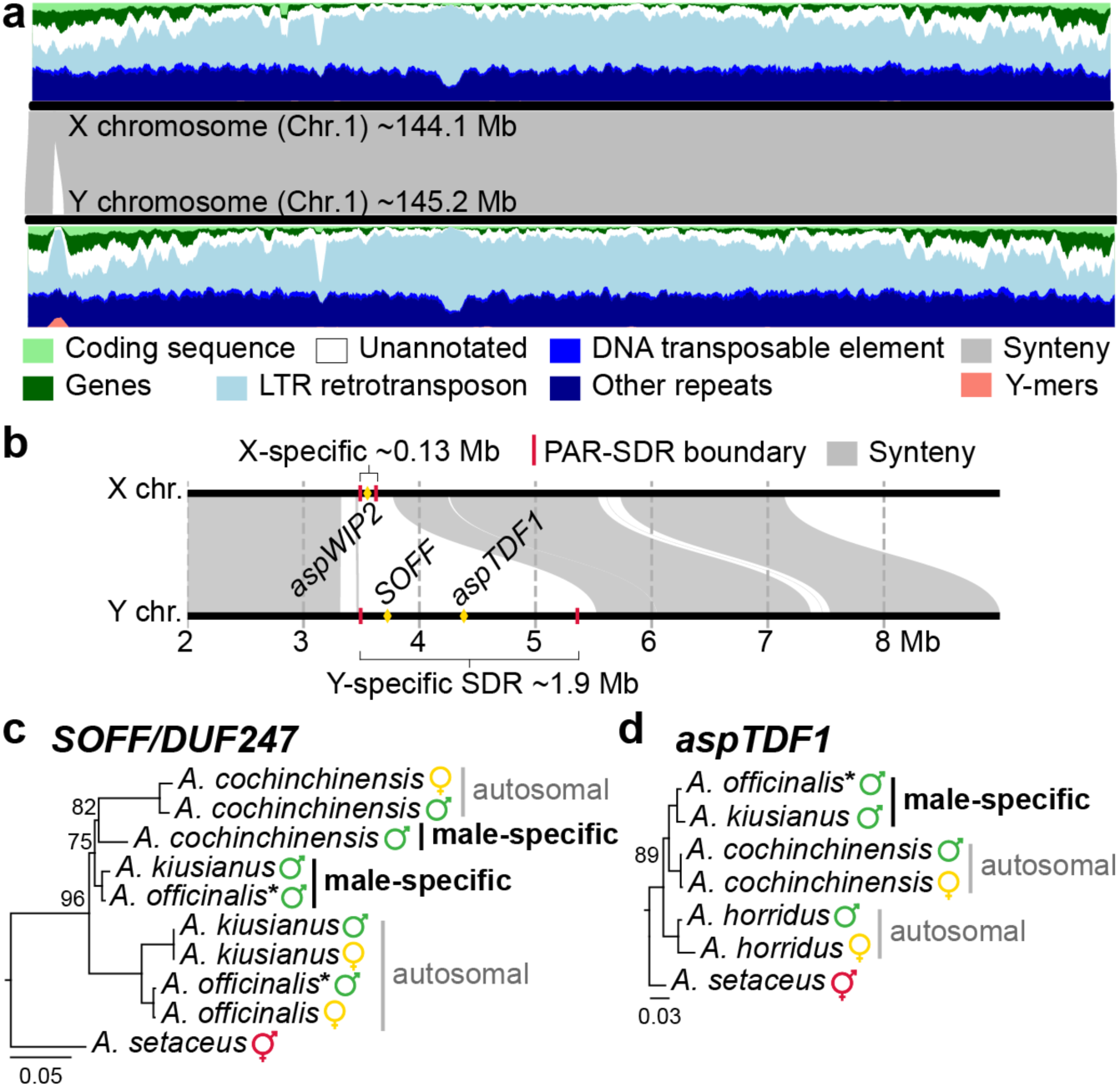
XY sex chromosomes, in the new haplotype-resolved garden asparagus (*Asparagus officinalis*) genome, correspond to chromosome 1 (9)**. a)** X-Y haplotype alignment (middle track) and structural annotation densities (X=top; Y=bottom) show support for a hemizygous Y-specific sex-determination region (Y-SDR) where Y-mers (male-specific *k*-mers identified between 3 males - 3 females) and LTR retrotransposons are elevated compared to the surrounding PARs (pseudo-autosomal regions). **b)** Hemizygous regions between the XY pair mark the nonrecombining Y-SDR and X-specific region. The Y-SDR contains 10 genes, including two with sex-determining functions: *SOFF* and *aspTDF1* (diamonds) (8, 9), whereas the X-specific region only contains one gene (*aspWIP2*) and shares no gametologs with the Y. **c)** A *SOFF* (*DUF247* gene family) phylogeny revealed strong support for separate clades with either male-specific or autosomally-linked homologs from *A. officinalis* (chromosome 5) and *A. kiusianus*; and uncertain placement of the *A. cochinchinensis* male-specific homolog. The *SOFF* tree supports a recent duplication of an ancestral *DUF247* gene preceded neofunctionalization of sex-determining function in the most recent common ancestor (MRCA) of the Eurasian clade; which may have been lost in a common ancestor of *A. officinalis* and *A. kiusianus,* but maintained in *A. cochinchinensis.* **d)** The *aspTDF1* tree supports the stepwise recruitment of *aspTDF1* into the Y-SDR of a common ancestor of *A. officinalis* and *A. kiusianus*, following divergence from the MRCA shared with *A. cochinchinensis*. *Asparagus setaceus* is a bisexual species, representing the ancestral condition for the genus. *Agave* orthologs were used to root both gene trees (see Fig. 4d). Bootstrap branch support shown when <100% support. **\***gene models from the new *A. officinalis* genome.

Only one gene (*aspWIP2*) was found in the X-specific region (**Fig. 3b**). In Arabidopsis, *WIP2* is a zinc finger transcriptional regulator (*C2H2*-type) required for normal pollen tube growth and transport to ovules for fertilization (41). The function of *aspWIP2* in *A. officinalis* has not been tested, but its specificity on the X leaves open the possibility of a dosage advantage in females (two copies) relative to males (one copy) and potential sexually antagonistic function (8). A second gene model, an *outer envelope protein 80* ortholog, showed evidence of linkage with both the X and Y nonrecombining regions in earlier work (8, 9), but results from our analyses were inconclusive because none of its exons contained a Y-mer although separate clades of X-vs. Y-linked orthologs were moderately or poorly supported, respectively (**Fig. S8**). We placed this *outer envelope protein 80* homolog in PAR2 of both haplotypes (**Table S9**); however, the germplasm used here differs from the previous assemblies (8, 9), so it is possible that the boundary of the nonrecombining SDR varies within the species. In sum, the new genome for *A. officinalis* provides improved assembly of the X-Y nonrecombining regions and sex-limited gene annotations, due its increased contiguity enabled by PacBio HiFi+Omni-C sequencing. Additionally, by applying Y-mer mapping and phylogenetic methods, we found increased resolution of the PAR-SDR boundaries in *A. officinalis* (**Table S2**).

### Sex chromosome evolution in the Eurasian clade

Investigation into the evolutionary origin of the *A. officinalis* Y-SDR has been difficult due to the hemizygous nature of the X- and Y-limited regions (8, 9), leaving inference of the genomic mechanism(s) responsible for the origin of proto-XY recombination suppression unresolved for the Eurasian clade. We leveraged recently published chromosome-scale reference genomes representing two additional Asparagaceae subfamilies (Agavoideae and Nolinoideae) (42–44) to investigate the Y-SDR origin in the *Asparagus* Eurasian clade. Inference of syntologs vs. lineage-specific structural rearrangements (summarized in **Fig. 4a**) revealed no structural variation associated with the PAR-SDR boundaries in *A. officinalis.* However, PAR-linked regions, immediately adjacent to the *A. officinalis* Y-SDR on chromosome 1, exhibited large blocks of syntologs on one autosome (chromosome 5) in *Asparagus* (Asparagoideae), two in *Dracaena* (Nolinoideae) (**Fig. 4b**), and three in*Yucca* (Agavoideae) (**Table S10**). One *SOFF* homolog was located on chromosome 5 in *A. officinalis*, but not in a syntenic block. To that end, no syntologs were identified for any of the Y-SDR-linked genes in *A. officinalis,* altogether suggesting that these genes entered in the Y-SDR in a stepwise manner following the establishment of a nonrecombining *SOFF* locus on an ancestral proto-Y. Interestingly, we found syntologs of the X-specific *aspWIP2* on chromosome 5 in all analyzed *Asparagus* species (**Table S10**), thus we hypothesize that the ancestral Y-linked allele was lost sometime following the origin of dioecy in the Eurasian clade. We then tested whether the observed relationship between *Asparagus* chromosomes 1 and 5 could be traced back to a whole genome duplication (WGD) or a smaller, segmental duplication, and if either were associated with the origin of dioecy in the Eurasian clade. Analysis of synonymous substitutions indicates that many syntologs on the *Asparagus* chromosomes 1 and 5 arose from an ancient WGD shared with other Asparagaceae subfamilies >41 Ma (Asparagoideae-Nolinoideae Ks tests shown in **Figs. 4c, S9**), well before the origin of dioecy in *Asparagus.* This inference agrees with previous analysis of copy number variation (paralogs vs. orthologs) in de novo transcriptome comparisons of Agavoideae and Asparagoideae taxa (9).

**Figure 4.**
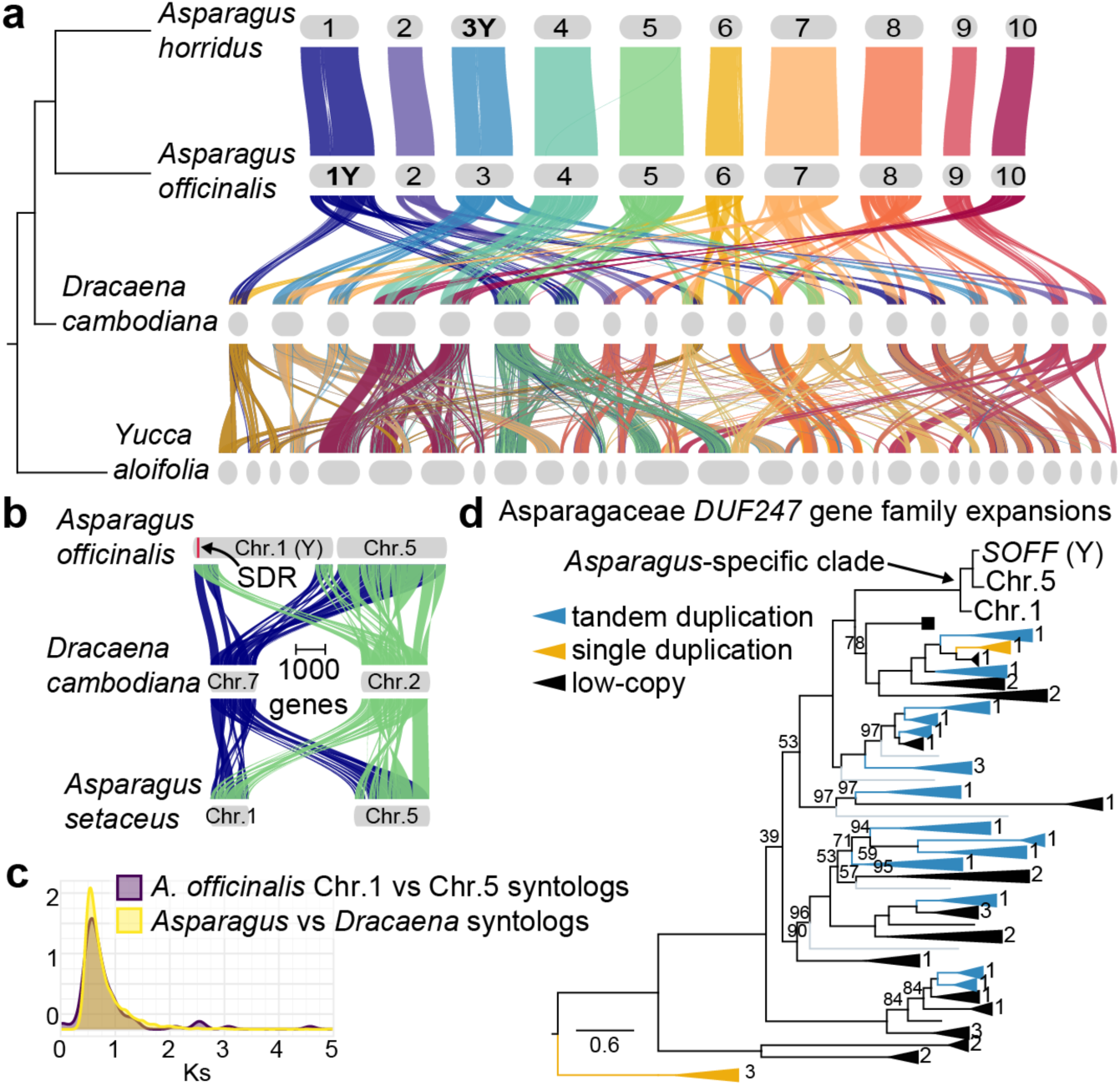
Two XY sex chromosome systems evolved from different ancestral autosomes in *Asparagus*, a genus in the Asparagaceae subfamily Asparagoideae. Chromosome 1 represents the XYs in *A. officinalis* and chromosome 3 represents the *A. horridus* XYs (only Y chromosome haplotypes shown). **a)** Syntolog relationships across three Asparagaceae subfamilies illustrate the rampant genome rearrangements that occurred across >41 million years of lineage divergence. *Dracaena*=Nolinoideae; *Yucca*=Agavoideae. **b)** The *A. officinalis* sex-determination region (SDR) on the Y is nested between syntologs shared with chromosome 5 of *Asparagus* and two others from *Dracaena*. *Asparagus setaceus* is a hermaphroditic species. **c)** Plot illustrates overlapping synonymous substitution (Ks) distributions, measured between *A. officinalis* chromosome 1-5 syntologs from SDR surrounding regions (purple curve), and compared to that of all *Asparagus*-*Dracaena* syntologs (yellow curve). This suggests that an ancient genome duplication, predating the origins of dioecy in *Asparagus* by at least 38 million years, is responsible for the observed chromosome 1-5 homology in *Asparagus.* **d**) Lineage-specific *DUF247* (*SOFF*) gene family expansions are common across Asparagaceae taxa and usually occur via tandem duplications (blue clades). The Y-specific *SOFF* in *A. officinalis* arose in an *Asparagus-*specific clade with autosomal homologs from chromosomes 1 and 5. Single gene duplications=yellow clades. Clades were collapsed and labeled with the number of Asparagaceae subfamilies sharing the indicated duplication pattern. Black box tip marks the *SOFF* gene tree root used in Fig. 3c. Grey branches are rice homologs used as a control for major clades (46). Bootstrap branch support shown when <98%.

Analysis of the *DUF247* gene family across multiple Asparagaceae taxa revealed no closely related *SOFF* orthologs outside of *Asparagus* (**Fig. 4d**, SupplementalFile10.pdf), nor were any identified in a separate analysis with wider sampling (46). Phylogenetic analysis of *SOFF/DUF247* homologs from the hermaphroditic *A. setaceus* and three Eurasian dioecious species (*A. officinalis, A. kiusianus* and *A. cochinchinensis*) supports the hypothesis that a male-specific *SOFF* arose following a more recent single or tandem gene duplication in the MRCA of the Eurasian dioecy clade and that the *SOFF/DUF247* homolog on chromosome 5 likely represents an older paralog (**Fig. 3c**). The less well supported placement of *A. cochinchinensis SOFF/DUF247* homologs in **Fig. 3c** implies an independent set of duplications in the *A. cochinchinensis* lineage, but understanding the timing and nature of those duplications will require genome assemblies for *A. cochinchinensis* and relatives (**Fig. 1**). As seen in earlier work (46), phylogenetic analysis of *DUF247* genes shows many instances of gene family expansions by tandem duplications and variation in copy number across Asparagaceae lineages (**Fig. 4d**). Rampant copy number variation across *DUF247* homolog clades in Asparagaceae may also explain the absence of a closely related *SOFF* ortholog in *A. horridus*.

### Two independent XY sex chromosome systems evolved in Asparagus

In this study, we use genomic and evolutionary analysis to test for support for two independent origins of dioecy and sex chromosomes in *Asparagus* (14). We show that each origin of dioecy in the genus involved different ancestral autosomes: chromosome 1 in the Eurasian clade and chromosome 3 in the Mediterranean Basin clade (see bolded Y chromosomes in **Fig. 4a**). In *A. horridus*, the nearly 10 Mb Y-SDR is considerably larger than the almost 2 Mb Y-SDR in *A. officinalis*, despite being ∼1.7-2 million years younger; which supports the hypothesis that expansion of recombination suppression, as a measure of age and total SDR size, is not a linear relationship in plants (6) although both regions appear to have expanded over time. Aside from the presence of a non-orthologous *AP2* gene and higher ratio of repetitive sequences, compared to the X and autosomes, the only common patterns observed between the *A. officinalis* and *A. horridus* Y-SDRs included the secondary recruitment of LTR retrotransposons and other gene content (shown as duplications in **Figs. S10-S11**) driving their stepwise expansions. Intriguingly, however, both dioecy origins occurred within ∼1-2 million years of each other (**Fig. 2e**), within the same major clade in the genus (**Fig. 1** shows the “Asparagus clade” in the genus *Asparagus*), and in association with long-distance dispersals out of southern-central Africa to Eurasia (14, 15). Considering ancestral biogeography (southern-central Africa) and timeliness (∼1.1-3.8 Ma) of dioecy evolution, it is plausible that founder events associated with historical climate oscillations across central-northern Africa helped set the stage for both independent transitions from hermaphroditism to dioecy in the genus (12).

The origin of dioecy in the Eurasian clade is marked by the evolution of a male-specific *SOFF* co-opted for sex-determination, which was followed by the stepwise recruitment of additional genes including *aspTDF1* in some lineages (**Fig. 1**). The ancestral Y-linked *SOFF* may have evolved from a tandem duplication of an autosomal *DUF247* gene, which have since been lost in *A. officinalis,* but may still be present in *A. cochinchinensis* (**Fig. 3c**). Therefore, a single-gene hypothesis may explain the origin of dioecy in the Eurasian clade, since early diverging lineages (i.e., the *A. cochinchinensis* subclade) exhibit a male-specific copy of *SOFF* (**Fig. 3c**) but not *aspTDF1* (**Fig. 3d**) (9, 11)*. SOFF* knockouts in *A. officinalis* males result in functioning hermaphroditic flowers, (*aspTDF1* knockouts are male-sterile) (8, 9), indicating that *SOFF* expression does not impact pollen development in the species. Thus, experimental investigation of *SOFF* function in the *A. cochinchinensis* subclade is necessary to elucidate the ancestral sex-determination mechanisms in the Eurasian group. A single-gene model for the origin of dioecy in the Eurasian clade would require that the ancestral *SOFF* had some function in pollen development which was lost following the co-option of *aspTDF1* into the Y-SDR, in an ancestor of *A. officinalis*. Sexually antagonistic genes are predicted to accumulate in nonrecombining sex-limited regions over time and are thought to lead to sexual dimorphism (47). The stepwise recruitment of *aspTDF1* into the Y-SDR may have been a consequence of sexually antagonistic selection (i.e., removal of *aspTDF1* from the autosomes ensures that females can no longer produce pollen) in the MRCA of *A. officinalis* and *A. filicinus* or *A. verticillatus* (see **Fig. 1**).

Works presented here, together with phylogeographic analysis of the origin of dioecy in *Asparagus* (14) and functional work on other dioecious plant species (48), indicate that there are many potential molecular mechanisms for the shift from hermaphroditism to dioecy in flowering plants. Continued work on dioecious lineages of *Asparagus* offers opportunities for improved understanding of the ecological drivers of the origin and persistence of dioecy. For instance, two of the eight independent range expansions out of southern Africa were associated with dioecy origins in *Asparagus* (14) suggesting that long-distance dispersals and inbreeding avoidance has contributed to dioecy transitions, but specialization on male or female function may have contributed to its maintenance across time. In any case, integrated phylogenetic, genomic, and functional investigations of dioecy in model taxa such as *Asparagus* will continue to yield deeper understanding of the origins and evolution of separate sexes across the tree of life.

## METHODS AND MATERIALS

### Biological materials

Male (XY) plants were selected for genome assembly, to capture both sex chromosomes. Individual plants were selected for sampling based on tissue availability. Fresh cladodes were collected and flash frozen with liquid nitrogen for all DNA-seq experiments. We sampled several tissue types, in triplicates, at different developmental stages, for transcriptome sequencing from the *A. officinalis* and *A. horridus* genome lines, which we used for genome structural annotations (**Table S4**). We also sampled male and female flowers across different developmental stages (five replicates each), from a wild population of *A. horridus* identified at Capo Rama reserve WWF - Terrasini (PA) Italy for differential expression analysis between the sexes (**Table S4**). Tissue for transcriptome sequencing was flash frozen with liquid nitrogen immediately after sampling and all tissue was sampled at the same time, if used together in downstream analysis. Details about all samples from this study can be found in **Table S11**.

### DNA and RNA data generation

PacBio HiFi, Omni-C, and Illumina (PE150) libraries were all prepared at HudsonAlpha Institute for Biotechnology (Huntsville, Alabama, USA) using SMRTbell^®^ Prep Kit v2.0 (Pacific Biosciences, Menlo Park, California, USA), Dovetail Genomics Omni-C^®^ Kit (Cantata Bio, Scotts Valley, California, USA), and NEBNext Ultra II DNA PCR-free Library Prep Kit (New England Biolabs Inc., Ipswich, Massachusetts, USA), respectively. PacBio HiFi reads were sequenced on the SEQUEL II platform, whereas Omni-C and all other short-read DNA-seq data were sequenced on the Illumina (San Diego, California, USA) NovaSeq 6000. High molecular weight DNA extraction was performed using the Takara NucleoBond^®^ HMW DNA kit (Takara Bio USA, Inc., San Jose, California, USA) prior to PacBio HiFi library preparation. DNeasy Plant Mini kit (Qiagen, Hilden, Germany) was used for DNA isolation prior to Illumina library preparation. For the *A. horridus* and *A. officinalis* genome lines, total RNA was extracted from the various tissue types using RNeasy Plant Mini Kit (Qiagen), libraries were prepared using Illumina TruSeq Stranded mRNA Library Prep kit, and then sequenced on the Illumina NovaSeq 6000 platform at HudsonAlpha. All RNA replicates for each genome line were also pooled for PacBio HiFi Iso-Seq library preparation and long-read sequencing on the SEQUEL II platform at HudsonAlpha. Additional RNA-seq data sets were generated for male and female *A. horridus* from an Italian population (**Table S4)** as follows: 1) total RNA extraction with RNeasy Plant Mini Kit, 2) shipped from Italy to the U.S. on GenTegraRNA (GenTegra, Pleasanton, California, USA) columns to ensure RNA stability, 3) mRNA libraries prepared by Novogene Corporation Inc. (Sacramento, California, USA) using in-house protocols, 4) sequencing on the Illumina NovaSeq X-Plus (10B, PE150) platform.

### Genome assembly

We used HIFIAsm+HiC v0.16.1 (49) to build initial contigs and YaHS v1.1 (50) to scaffold those contigs into chromosome-scale assemblies. Prior to scaffolding, we used BWA-MEM v0.7.17 with the flag *-5SP* (51), SAMBLASTER v0.1.24 (52), and SAMtools v1.16.1 (53) to map Omni-C reads to contigs, mark duplicate alignments, and remove those duplicates, respectively.

HIFIAsm contigs <50,000 nt were also removed prior to scaffolding. We ordered and oriented contigs/scaffolds using the JUICER v1.6 (54) and Juicebox v1.11.08 (55) pipelines to match the Aspof.V1 (9) genome (https://phytozome-next.jgi.doe.gov/info/Aofficinalis_V1_1). Final assembly completeness was assessed using BUSCO v6.0.0 (viridiplantae_odb12) (56) and Merqury v1.3/Meryl v1.4.1 for *k*-mer tests (57).

### Genome structural annotation

We annotated repetitive elements and protein coding genes using repeat-soft-masked haplotype assemblies after generating repeat libraries de novo using RepeatModeler v2.0.2 (58) and soft-masking with RepeatMasker v4.1.2 and the options *-cutoff 250* and *-nolow.* Curated repeats from the Repbase database for monocots (59) were combined with our de novo library prior for the RepeatMasker analysis. We used the long-read protocol ( https://github.com/Gaius-Augustus/BRAKER/blob/master/docs/long_reads/long_read_protocol.md) for BRAKER v3.0.3 (60–62) and its many dependencies (63–72) for gene prediction based on extrinsic evidence from short- and long-read transcriptome sequencing of various tissue samples (**Table S4**) and published protein sequences from *Asparagus officinalis* (9), *Asparagus setaceus* (40), and Viridiplantae (OrthoDB v11) (73). Illumina transcriptome reads were aligned to soft-masked haplotypes with STAR v2.7.10 (74). Full-length (non-concatemer) consensus Iso-Seq reads were mapped to soft-masked haplotypes using pbmm2 v1.3.0 (75), then isoforms were collapsed in the mapped transcripts. Gene predictions were parsed and filtered using AGAT v1.1.0 (76) and GffRead v0.12.7 (77), discarding genes with 1) in-frame stop codons (or adjusting the CDS phase when possible), 2) single-exon transcripts when absent on the opposite strand, 3) missing start codons, or 4) total CDS <300-nt. EnTAP v1.0.0 (78) was used to further assess gene prediction accuracy via reciprocal functional annotation based on the UniProt/Swiss-Prot (79) and EggNOG v5.0 (80) databases. Mono:multi-exonic ratios were also used for gene prediction quality control assessment (21). TE prediction was performed with EDTA v2.2.2 (flags: *--sensitive 1 --anno 1 --evaluate 1*) (81) using the RepeatModeler library of classified elements, BRAKER gene predictions, and the ‘out’ file from RepeatMasker. TRF v4.09.1 (82) was used to annotate tandem repeats across each haplotype. SDR gene predictions were further curated by removing models that were assigned TE-associated annotations or >90% soft-masked. Completeness of gene predictions were assessed using BUSCO v6.0.0 (viridiplantae_odb12) proteins.

### Delimitation of sex chromosome nonrecombining regions

To identify X/Y haplotypes, we mapped male-specific *k-*mers (Y-mers) from Illumina short-read datasets for *A. officinalis* and *A. horridus* (**Table S11**) to each haplotype, separately for each species. All 21-bp *k*-mers, present in Illumina reads for both species, were counted using JELLYFISH v2.3.0 (83) and filtered by removing 21-mers present at low (<10) or high (>250) frequencies. Y-mers were subset by selecting for 21-mers conserved across all male samples but absent in females from each species (84). We used BWA-MEM (flags: *-k 21 -T 21 -c 10 -a*) to map Y-mers, then delimited Y chromosomes and SDRs according to scaffolds and regions with the highest Y-mer coverage peaks, respectively (84). In a separate analysis, Y-mers from *A. acutifolius* were processed with those from *A. horridus* to test for a shared SDR. Normalized Illumina reads from all *A. horridus* samples (**Table S11**) were mapped to both haplotypes for the species, to perform coverage comparisons between the sexes for SDR delimitation (85), using BWA-MEM, requiring a 35 bp minimum seed length, <10 multi-mapped alignments, and >30 mapping quality. BBMap v38.93 *reformat.sh* (86) was used to normalize read depth by random down-subsampling to ∼30x. Using rough SDR coordinates based on Y-mer mapping density, we assessed gene tree topologies for the relative placement of X-Y gametologs/orthologs for genes near putative PAR-SDR boundaries. Y-SDRs and adjacent PARs (PAR1 and PAR2) were defined according to the first and last Y-specific gene/allele, as indicated by strongly supported clades of either Y-linked or X-linked orthologs (e.g., genes in a clade of only Y-linked orthologs were assigned to the Y-SDR). Orthologs/gametologs were identified using OrthoFinder v2.5.5 (87) and GENESPACE v1.3.1 (88) with both new haplotypes from *Asparagus officinalis* and *Asparagus horridus* as well as other monocot relatives: *Asparagus setaceus* (40)*, Asparagus kiusianus* (45)*, Dracaena cambodiana* (44), *Yucca aloifolia* (43), and *Ananas comosus* (pineapple) (89). Chromosome labels from the *A. setaceus* genome were renamed here to match those from this and previous studies (8, 9).

GENESPACE results were also used to identify structural variants among syntenic, orthologous gene blocks. Haplotype-haplotype alignments were also generated to test for SVs between the XYs within each species, using minimap2 v2.26 (75) and SyRI v1.6.3 (90), respectively. SVs were plotted with plotsr v1.1.0 (91).

### Sex chromosome evolution in A. horridus (Mediterranean Basin clade)

We performed pairwise comparisons of Ks between Y-SDR + flanking PAR genes vs. X-gametologs (primary transcripts). We tested for a linear correlation between X chromosome position and Ks to test for large step-wise SDR expansion events and rule out the presence of evolutionary strata in the *A. horridus* Y-SDR. X chromosome position should be more preserved over time compared to the Y-SDR, due to no meiotic crossing over in the latter.

Correlation tests were performed using base R (v4.2.2) (96) to calculate Pearson’s product-moment correlation coefficient, Kendall’s rank correlation (tau) coefficient, and Spearman’s rank correlation (rho) coefficient (*p*-value <0.05 cut-off). Ks was also measured between genome-wide homologs from *A. officinalis* and *A. horridus*, to estimate total species divergence. Three separate Wilcoxon signed-rank tests were then performed, because data were not normally distributed, to test for significant (*p*-value <0.5) differences in Ks for the following comparisons: 1) *A. horridus* Y-SDR genes vs. X-gametologs (N=47); 2) *A. horridus* Y-SDR genes vs. *A. officinalis* orthologs (N=41); and 3) *A. horridus* vs. *A. officinalis* genome-wide, single copy orthologs (N=12,646). All Ks estimates were calculated with KaKs_Calculator 3.0 (97) from protein alignments converted to nucleotide codon alignments with MAFFT v7.505 (98) and pal2nal v14 (99), respectively. To estimate the absolute timing of sex chromosome origins in the Mediterranean Basin clade, we used previous divergence time estimates for the MRCA of *A. horridus* and *A. officinalis* and multiplied the 95% confidence intervals by the percentage of test 1 to tests 2 and 3 from above. We then tested for signs of positive selection among Y-SDR genes in *A. horridus* by calculating d_N_/d_S_ ratios with PAML v4.8 (100) CODEML branch-sites model (M2a) and two nulls: the M1a sites model for nearly neutral selection; and the M2a branch-sites null which fixes omega (d_N_/d_S_) to 1 (neutral) for the foreground branch and is more stringent than the former (101). New ML gene trees were estimated for CODEML and consisted of only *Asparagus* orthologs identified by OrthoFinder. Gene trees were inferred using IQ-TREE v1.6.12 (93) with 1000 ultrafast bootstraps and the best fit substitution model (94). Trees with only three tips were manually constructed based on species relationships (14). M1a sites model results were assessed using a likelihood ratio test (LRT) and chi-squared distribution to compute *p-*values (*P*) and a sequential Bonferronitype procedure to control the false discovery rate by computing the expected rate of false rejection (*Q*), requiring *P*<*Q* (102). To compare the nested M2a models, LRT and chi-squared critical value thresholds were applied based on 1 degree of freedom (i.e., 3.84 LRT = *P* of 0.05; 6.63 LRT = *P* of 0.01) to compute relative *P* (<0.05 cut-off).

### Asparagus horridus male vs. female gene expression

To test for sex-specific expression patterns were compared between male and female flower tissues sampled from different developmental stages from an Italian population of *A. horridus* at the same time and location. RNA extractions with sufficient yield were sequenced (**Table S4**), then transcriptomes were assembled de novo with StringTie v2.2.1 (101), using *A. horridus* HAP1 transcript alignments generated with STAR. Differential expression analysis was performed using DESeq2 (102) with the StringTie gene count matrix. Expression similarities across flower (Italian population) sampling treatments were assessed based on PCA and clustered heatmaps (**Figs. S12-S13**). We tested whether sex (i.e., male vs. female) explained significant differential expression patterns among the floral sampling treatments, rather than comparing individual developmental stages, because expression profiles from each treatment were too overlapping among the successful RNA-seq libraries (**Figs. S12-S13**). Significantly different expression profiles were assessed based on adjusted *p*-value (>0.05) and log2 fold change indicating significant up-regulation (>0.99) or down-regulation (<-0.99).

### Sex chromosome evolution in A. officinalis (Eurasian clade)

Previous work proposed that a segmental duplication and translocation, including an autosomal ancestor of *SOFF,* from chromosome 5 to chromosome 1, drove the origin of proto-XY recombination suppression via hemizygosity (8, 9). To test this, we tested whether the origin of the hypothesized duplication event aligned with both the Asparagaceae and *Asparagus* phylogenies. First, we used GENESPACE and OrthoFinder results to infer homologous, syntenic gene blocks among *A. officinalis, A. horridus, A. setaceus, Dracaena,* and *Yucca*— with a focus on the Y-SDR + flanking PARs near the left end of chromosome 1 in *A. officinalis*. Then we measured Ks using wgd v2 (92) and compared results from two different treatments: 1) between paralogs from the *A. officinalis* chromosomes 1 and 5 to estimate the relative timing for the duplication event in question; 2) between genome-wide orthologs from *Asparagus* and *Dracaena* to estimate the relative timing of species divergence. Homologs/paralogs from the *A. officinalis* chromosomes 1 and 5 included only those identified from the first 400 gene models on chromosome 1 of haplotype 1 (i.e., selecting genes that span the PAR1-SDR-PAR2 borders) that exhibited syntenic homologs on chromosome 5 (86/400 genes fit these criteria).

*SOFF* and *aspTDF1* multiple sequence alignments were inferred using MAFFT v7.490 (flags: *--maxiterate 1000 --localpair*), then trimmed to remove poorly aligned regions with trimAl and the flag *-automated1.* Gene trees were inferred with IQ-TREE v1.6.12 (93), using 1000 ultrafast bootstraps, and the best fit substitution model (94). Homologs were identified in published assemblies for *Asparagus setaceus* (104), a male and female of *Asparagus kiusianus* (45), a female of *Asparagus officinalis* (8), and the outgroup species *Agave tequilana* (43) that was used for rooting. The *Asparagus kiusianus* assemblies lacked *SOFF* annotations, but we identified those using a local BLAST search. In *A. setaceus,* the *SOFF* homolog on chromosome 1 was split into two gene models that we concatenated according to BLAST alignments. The *SOFF* and *aspTDF1* trees were rooted with the *Agave tequilana* gene models *AgateH1.23G025400.1.v2.1* and *Agave_AgateH1.26G044700.1.v2.1*, respectively. The root for the *SOFF* tree was determined based on a larger phylogenetic analysis of the *DUF247* gene family (see SupplementalFile11.pdf) across *Asparagus, Agave, Dracaena*, and rice clade references from Zhu et al. (2025). All *DUF247* gene predictions were selected for phylogenetic analysis, based on inference of gene functions by EnTAP (performed as previously described), from all four *Asparagus* haplotypes from this study and the cited *Asparagus setaceus, Agave tequilana,* and *Dracaena cambodiana* assemblies. Prior to running EnTAP, genome annotations from other studies were re-filtered using AGAT and GffRead, as executed previously in this study. *The DUF247* gene family tree was inferred as described for *aspTDF1* and *SOFF*, but using amino acid alignments instead of nucleotides due to increased sequence diversity across the *DUF247* family. *Asparagus kiusianus* genes were not included in the broader *DUF247* analysis because the published genomes were missing protein predictions for those genes.

## ACKNOWLEDGEMENTS

We thank Laura Genco and Davide Bonaviri (WWF Italy, Capo Rama reserve), Tony Avent (Juniper Level Botanical Garden, Raleigh, NC, USA), and Mason McNair (Michigan State University, East Lansing, MI, USA) for their support with plant sampling. We thank Zach Stansell for providing seed collections for *Asparagus officinalis* from US Department of Agriculture’s Germplasm Resources Information Network (GRIN). We thank the University of Georgia and HudsonAlpha for computational, wet lab, and greenhouse support.

## AUTHOR CONTRIBUTIONS

P.C.B. wrote the manuscript and performed computational analysis. P.C.B., S.B.C., A.H., and J.L.-M. conceived the study and performed early analysis. P.C.B., F.M., V.R., and H.H. conducted field and/or laboratory experiments. P.C.B., F.M., and F.S. collected and curated biological samples. All authors reviewed manuscript drafts.

## DATA AVAILABILITY STATEMENT

Sequencing reads and genome assemblies/annotations are available under the NCBI BioProjects (TBD) and (TBD). Supporting scripts and additional files are available on GitHub (url) and Zenodo (url/doi).

## SUPPLEMENTAL FIGURES

**Figure S1.**
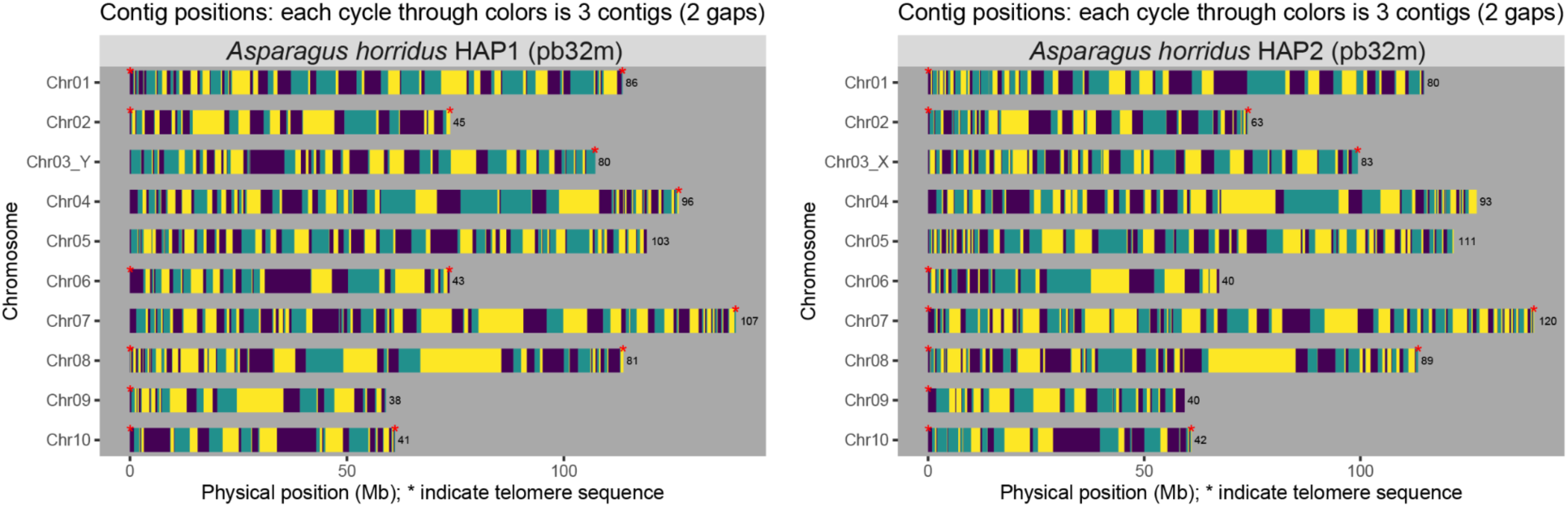
Contig (alternating color blocks) and telomere (red asterisk) maps from each haplotype assembly for *Asparagus horridus* (pb32m). Plots generated with GENESPACE (v1.3.1). Haplotype 1 = left panel. Haplotype 2 = right panel.

**Figure S2.**
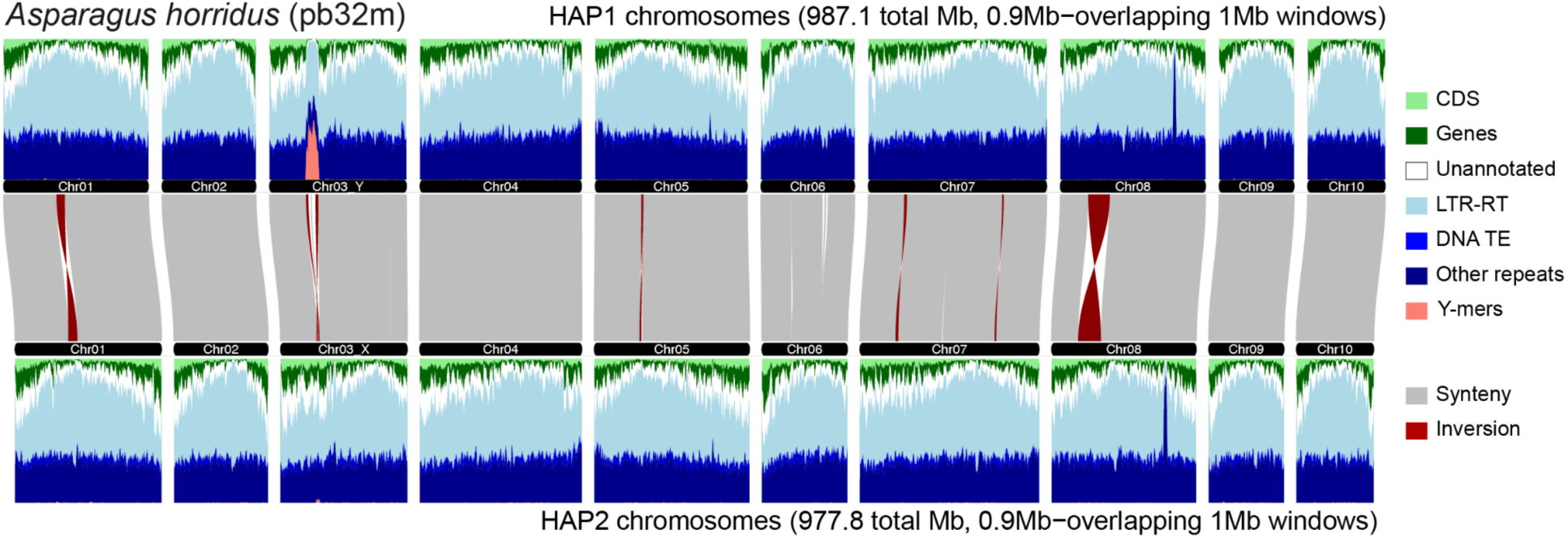
*Asparagus horridus* (pb32m) pseudo-chromosome alignments and annotation densities between each haplotype. Male-specific *k-*mer (Y-mer) mapping density is also shown (salmon color) and corresponds to the nonrecombining sex-determination region on chromosome 3. Plot generated using GENESPACE (v1.3.1) plot_2genomes.

**Figure S3.**
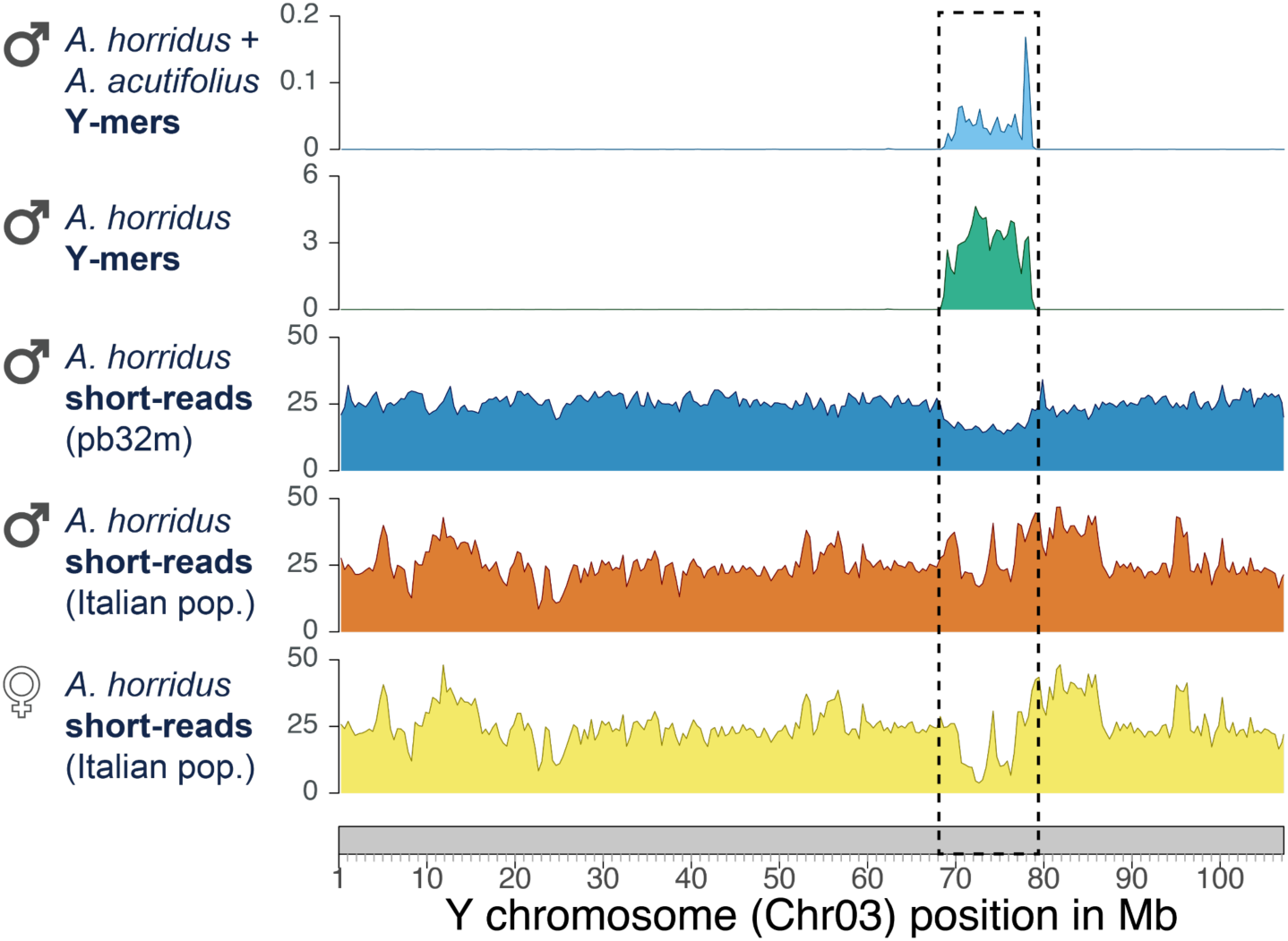
Evidence from read mapping supports a ∼9.6 Mb Y-linked sex-determination region (Y-SDR) on chromosome 3 (dotted box) of *Asparagus horridus*, corresponding to a single large inversion. Top two panels show read mapping depth of male-specific *k-*mers (Y-mers). Top track = Y-mers conserved in *A. acutifolius* + *A. horridus.* Second track from top = Y-mers counted in *A. horridus*. Bottom two tracks show male vs. female Illumina read mapping (see significant drop in female read density in the Y-SDR), compared to the reference male (middle track: see density decrease in the Y-SDR). The Italian samples are from a different population relative to the reference male (from Spain) and are examples of possibly population-specific variation in Y-SDR-linked content/structure. All tracks show mapping depth in 500-Kb sliding windows.

**Figure S4.**
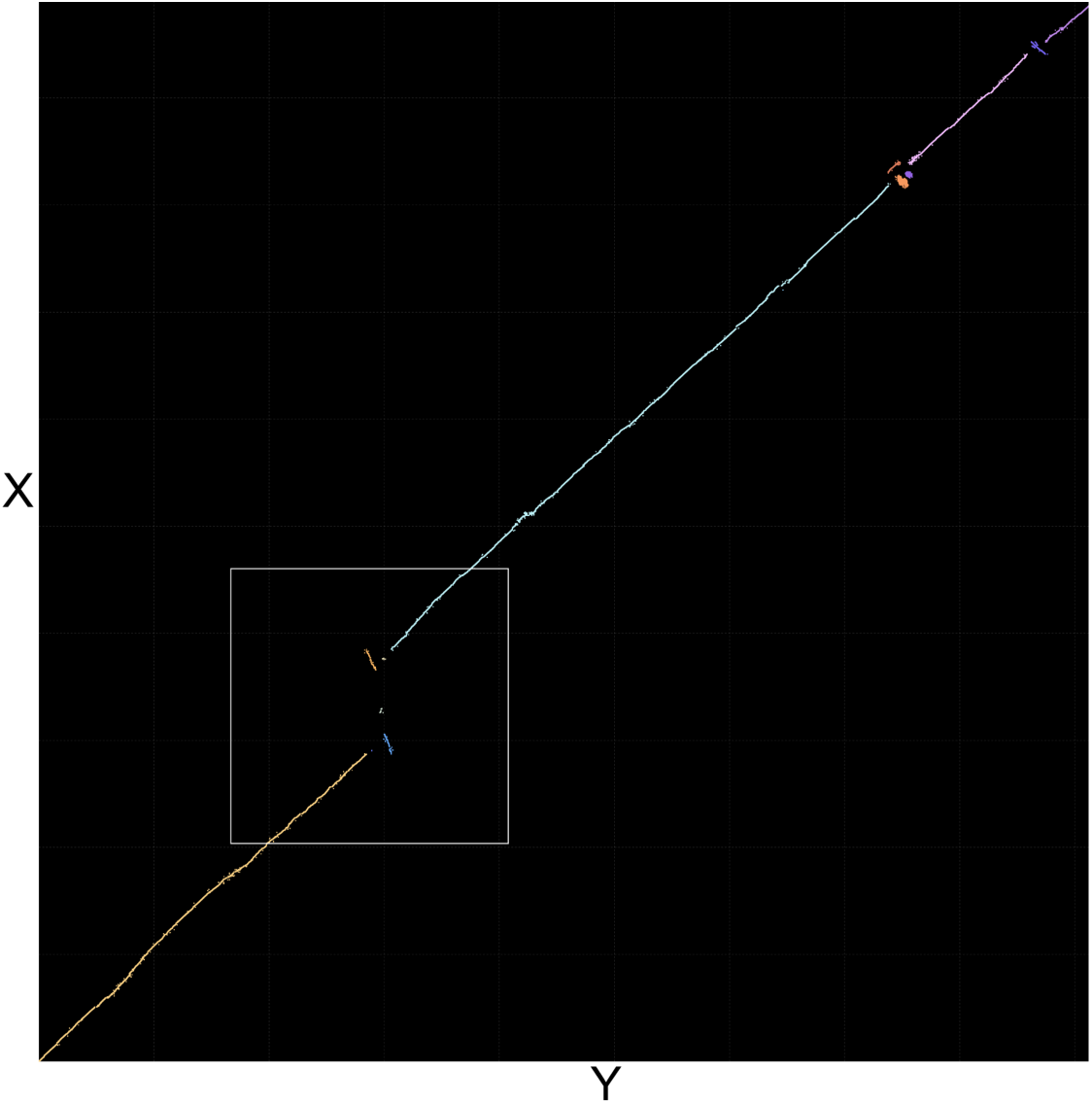
*Asparagus horridus* X-Y alignment dot plot (nucleotide alignment) showing evidence of a small nested inversion within a larger inversion. The larger inversion marks the nonrecombining region boundaries.

**Figure S5.**
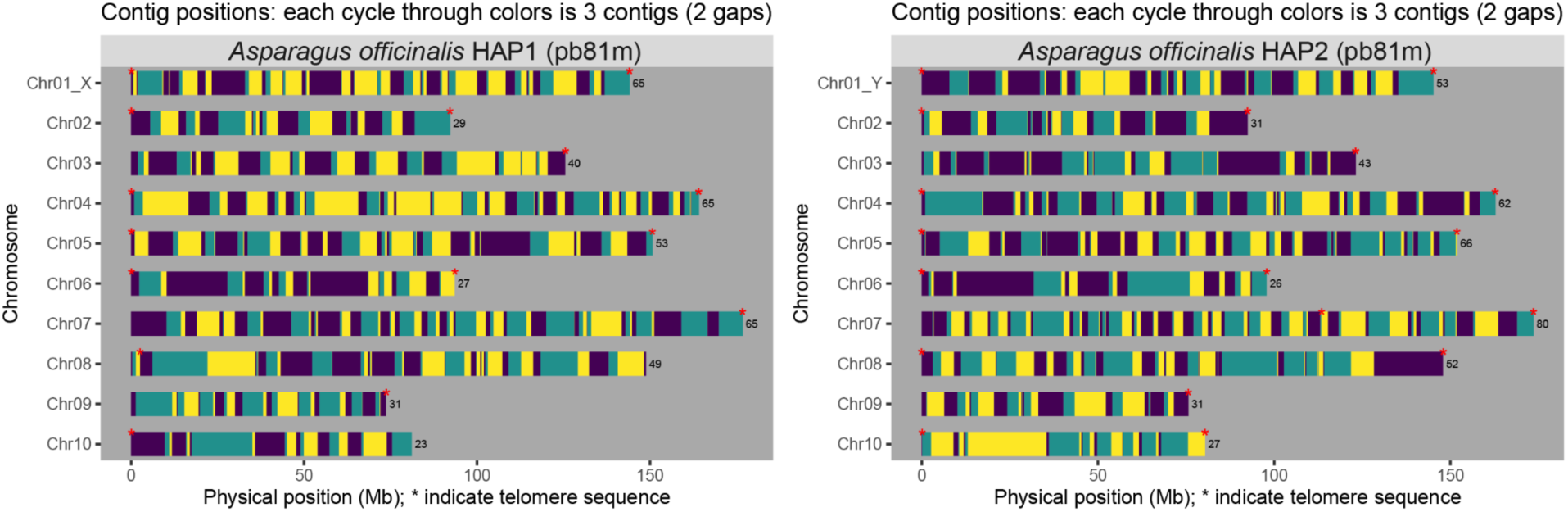
Contig (alternating color blocks) and telomere (red asterisk) maps from each haplotype assembly for *Asparagus officinalis* (pb81m). Plots generated with GENESPACE (v1.3.1). Haplotype 1 = left panel. Haplotype 2 = right panel.

**Figure S6.**
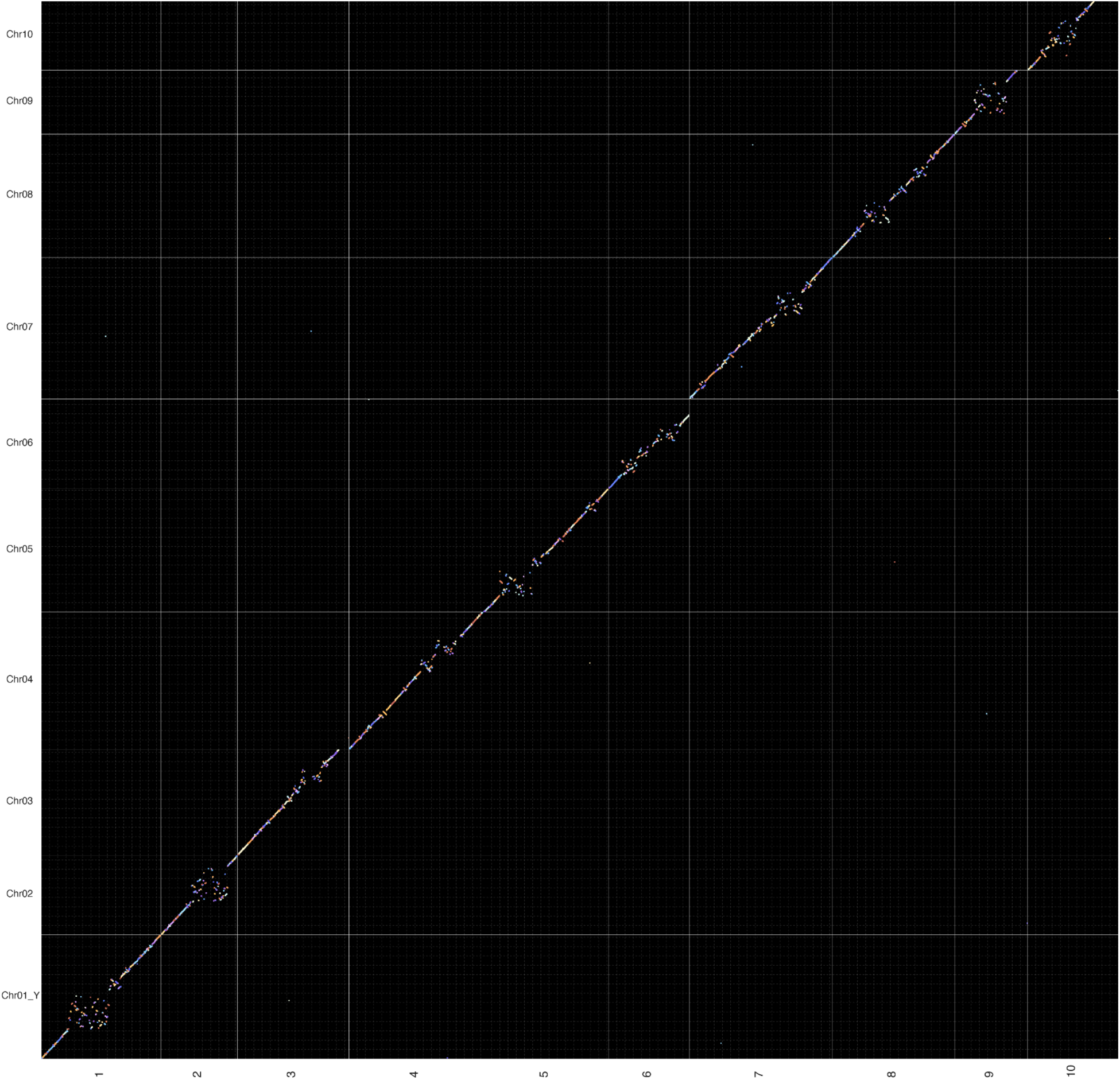
Dot plot alignment between the *Asparagus officinalis* YY double haploid assembly from Harkess, et al. (2017) (X-axis) and haplotype 2 (pb81m) from this study (Y-axis). Chromosome 1 = the Y chromosome in both assemblies. Plot generated using the GENESPACE (v1.3.1) clean_windows().

**Figure S7.**
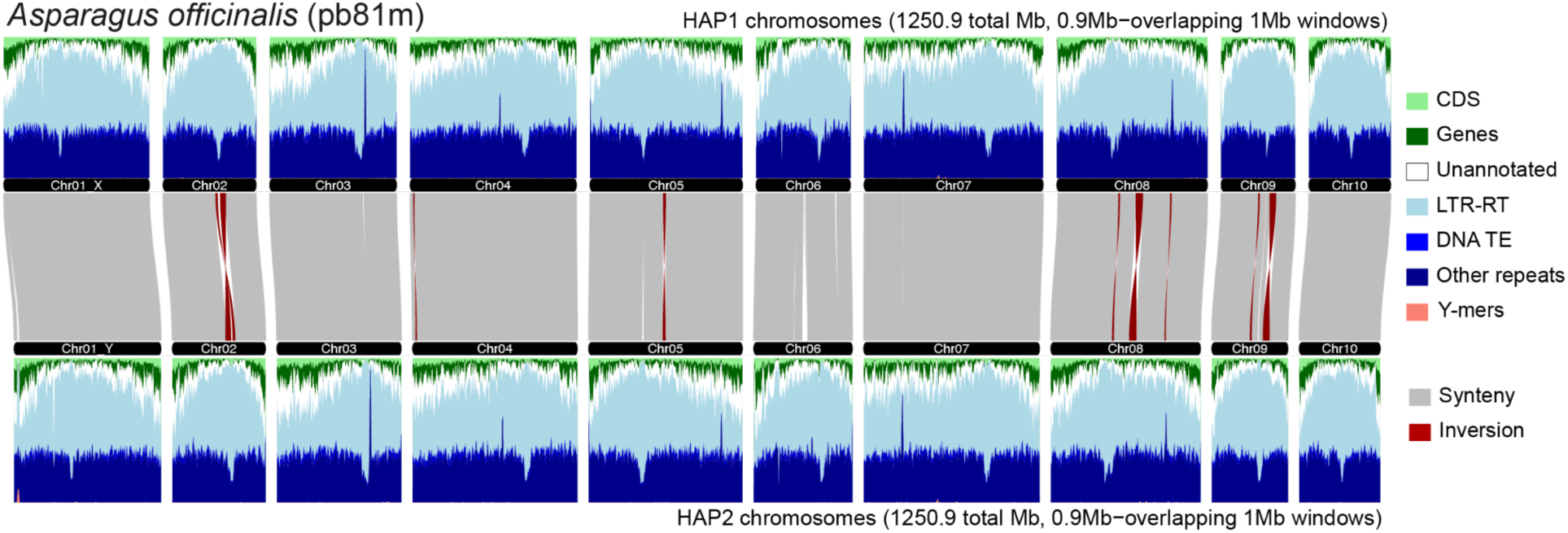
*Asparagus officinalis* (pb81m) pseudo-chromosome alignments and annotation densities between each haplotype. Male-specific *k-*mer (Y-mer) mapping density is also shown (salmon color) and corresponds to the nonrecombining sex-determination region on chromosome 1. Plot generated using GENESPACE (v1.3.1) plot_2genomes.

**Figure S8.**
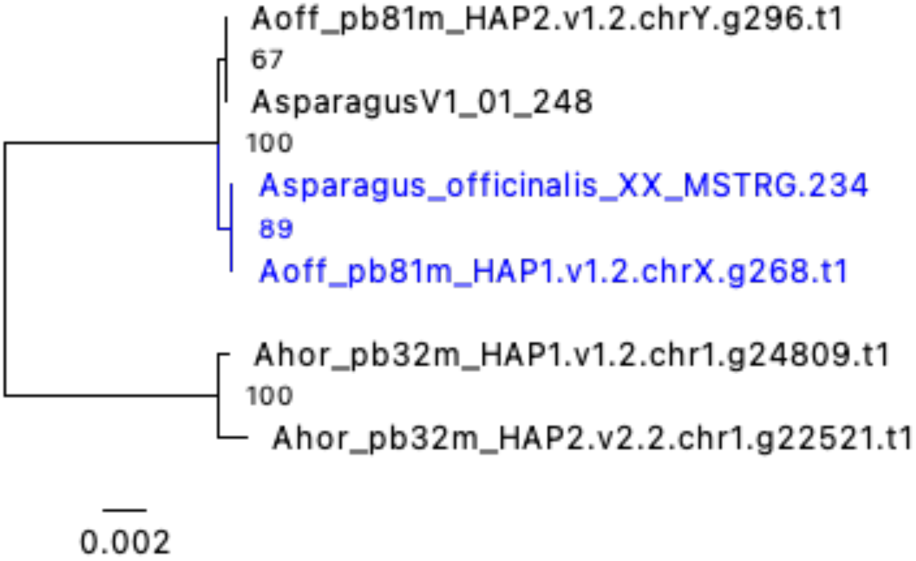
Maximum likelihood gene tree of an *outer envelope protein 80* gene that was found to be in the nonrecombining region on the X (*Asparagus_officinalis_XX_MSTRG.234*) and Y (*AsparagusV1_01_248*) in previous studies (Harkess et al., 2017, 2020) but was placed in the pseudoautosomal region (PAR2) of this study, due to poor support for a Y-specific clade (67% bootstrap support) and lack of Y-mers mapping to exons. Blue clade = X-linked homologs. The tree is rooted with *Asparagus horridus* orthologs.

**Figure S9.**
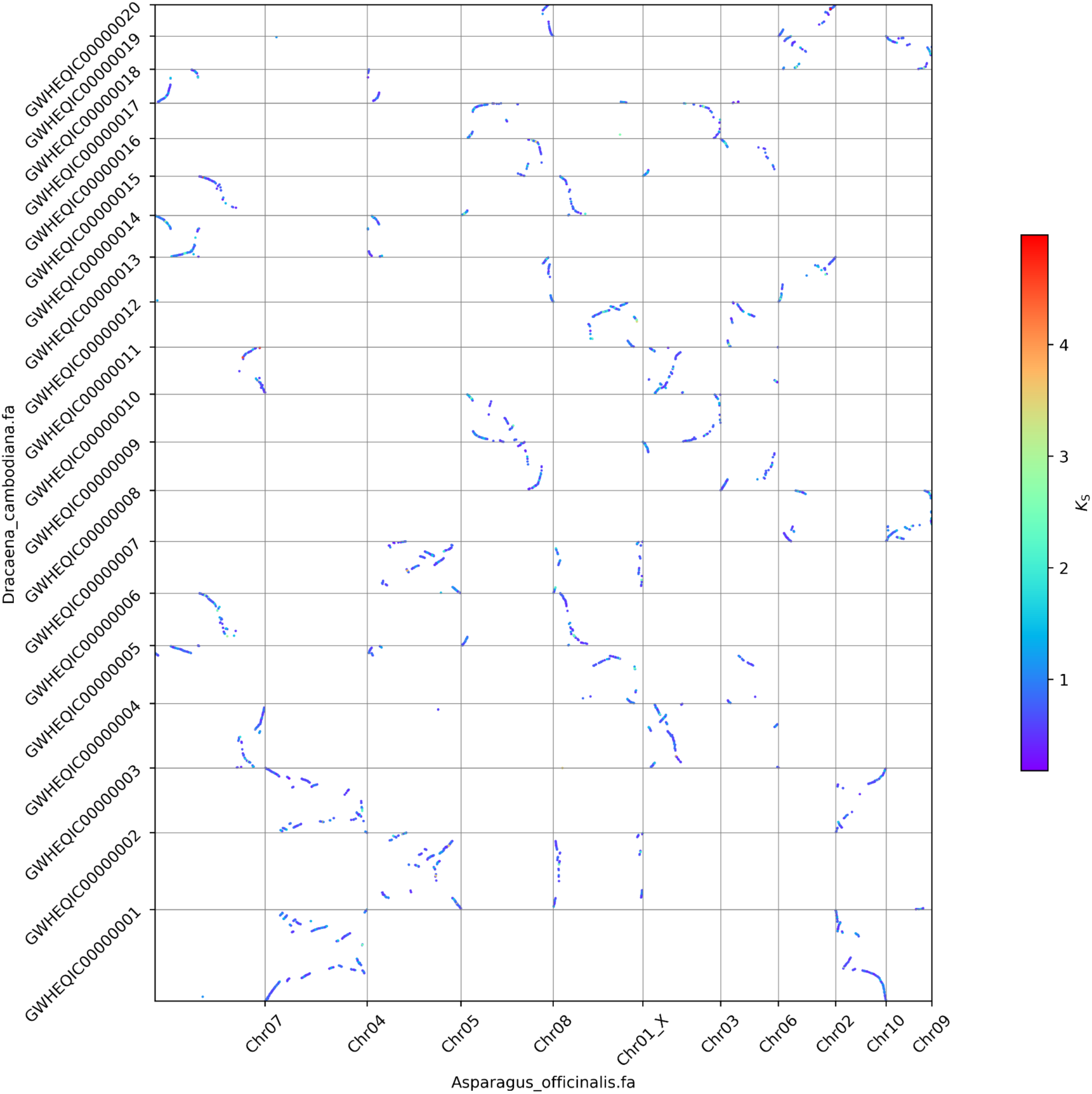
Syntenic gene block (syntolog) alignments between *Asparagus* (Asparagoideae) and *Dracaena* (Nolinoideae) colored according to Ks (synonymous substitution measurements) showing signatures of a shared genome duplication event with syntologs present on ∼2 chromosomes in each species. The Ks distribution peaked at ∼0.5 between the species. Plot generated from analysis with wgd v2 (Chen et al., 2024).

**Figure S10.**
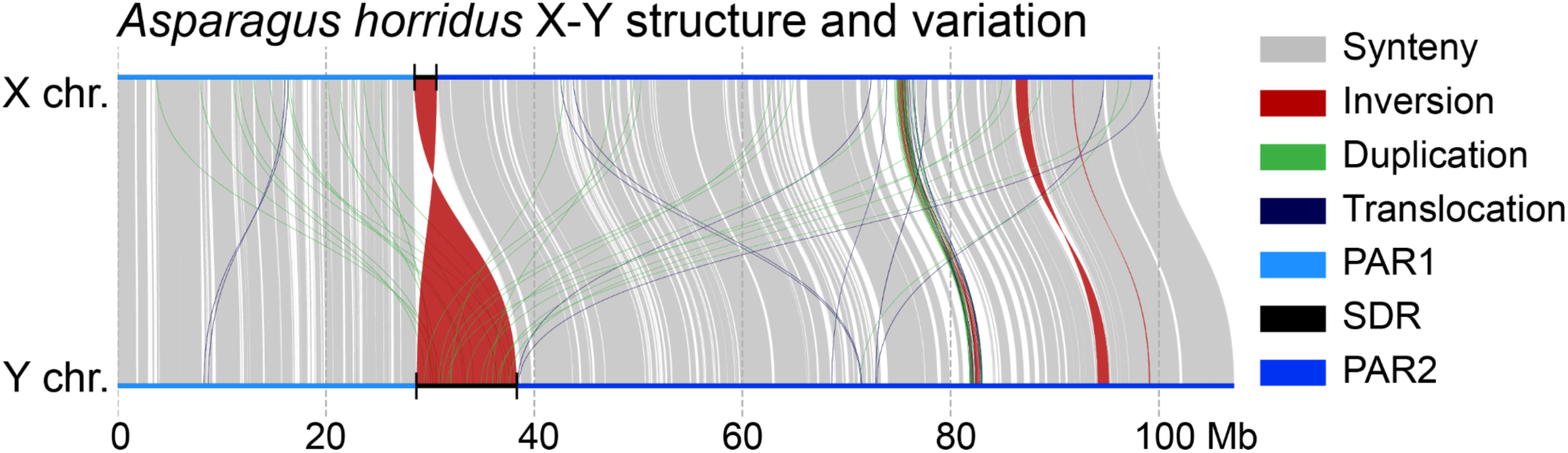
*Asparagus horridus* (pb32m) X-Y haplotype (chromosome 3) alignment showing structural variation and overall sex chromosome structure: the nonrecombining sex-determination region (SDR) is marked by black region with thin vertical bars marking boundaries with psuedo-autosomal regions (PARs) on either side of the SDR (marked by different shades of blue). Minimap2 (v2.26) and syri (v1.7.0) were used for haplotype alignment and SV detection, respectively. Plot generated using plotsr (v1.1.0).

**Figure S11.**
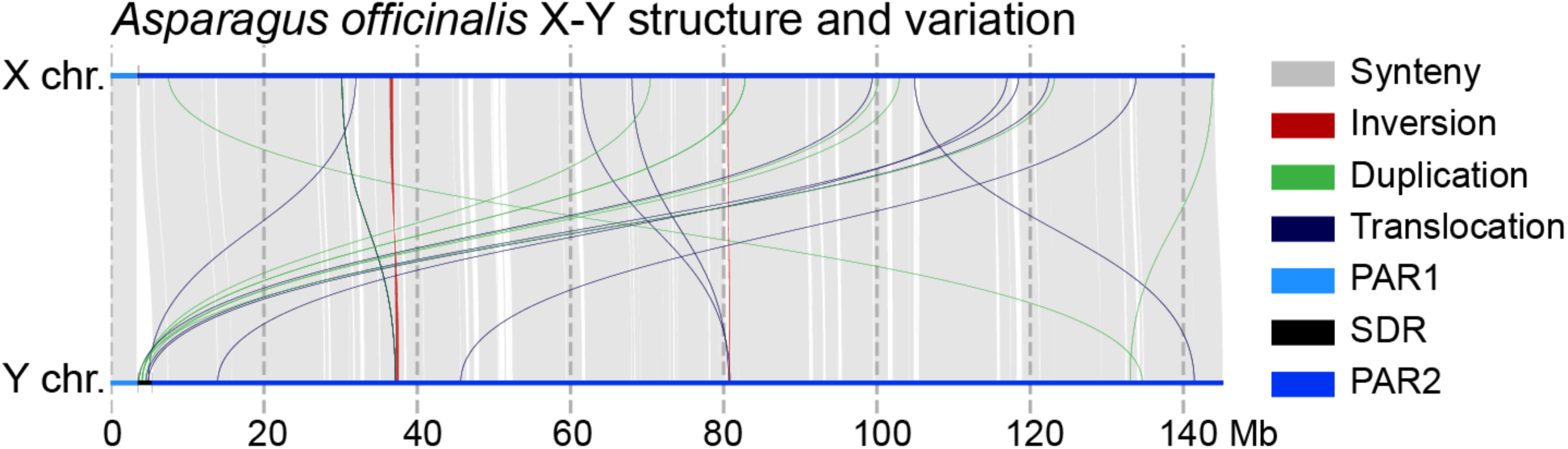
*Asparagus officinalis* (pb81m) X-Y haplotype (chromosome 1) alignment showing structural variation and overall sex chromosome structure: the nonrecombining sex-determination region (SDR) is marked by black region with thin vertical bars marking boundaries with psuedo-autosomal regions (PARs) on either side of the SDR (marked by different shades of blue). Minimap2 (v2.26) and syri (v1.7.0) were used for haplotype alignment and SV detection, respectively. Plot generated using plotsr (v1.1.0).

**Figure S12.**
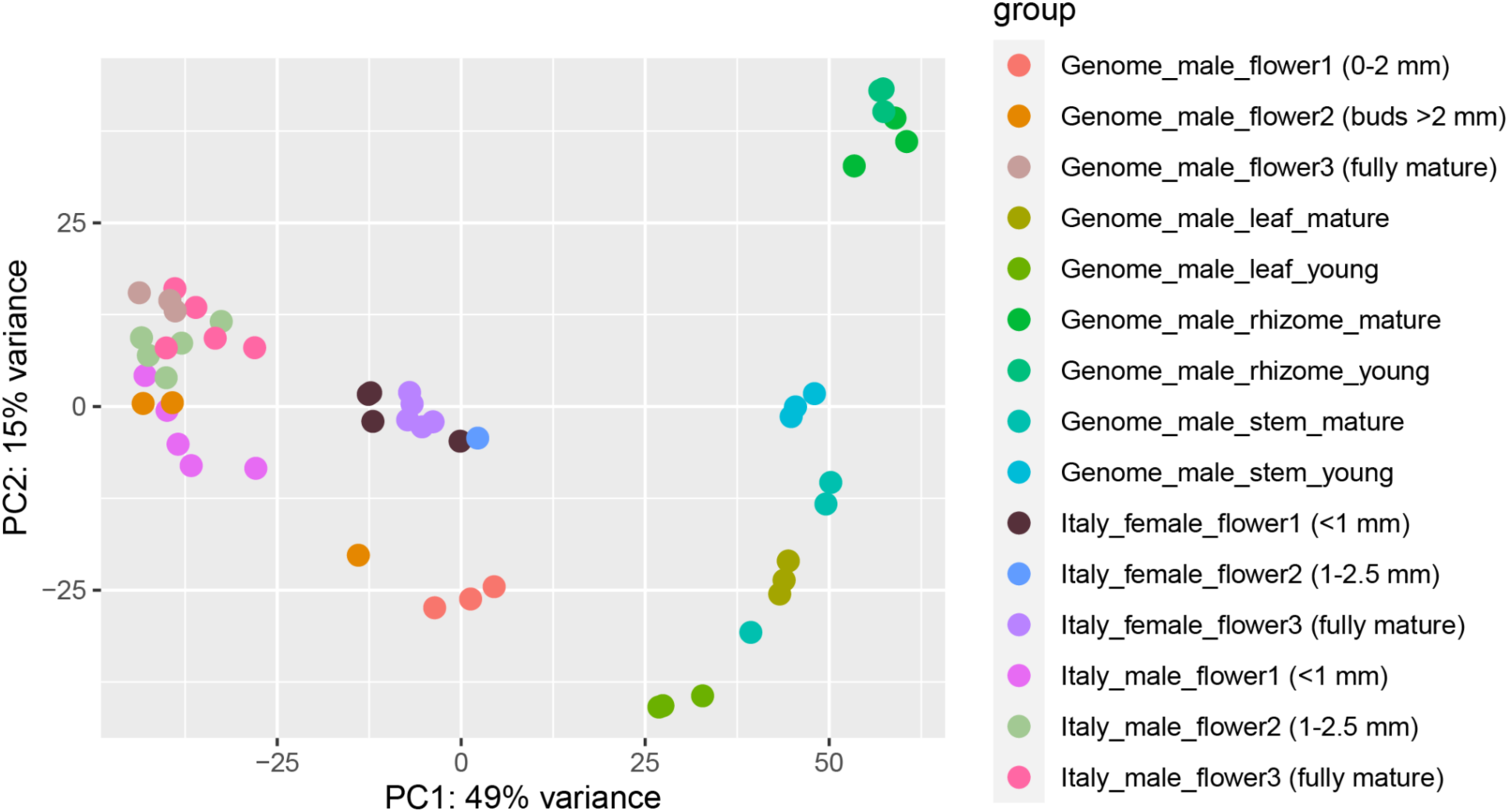
Principal component analysis of expression profiles from transcriptome sequencing of various tissues at different developmental stages from *Asparagus horridus.* Sample labels starting with “Genome_” are from the genome line plant (pb32m) and were used for structural annotation predictions. Sample labels starting with “Italy_” are tissues sampled at the same time and place from a wild population in Italy, which were used for differential gene expression analysis between the sexes. Floral stages for each sex, from the Italian population, were combined to measure expression differences between male and female flowers overall, because replicates did not consistently cluster together.

**Figure S13.**
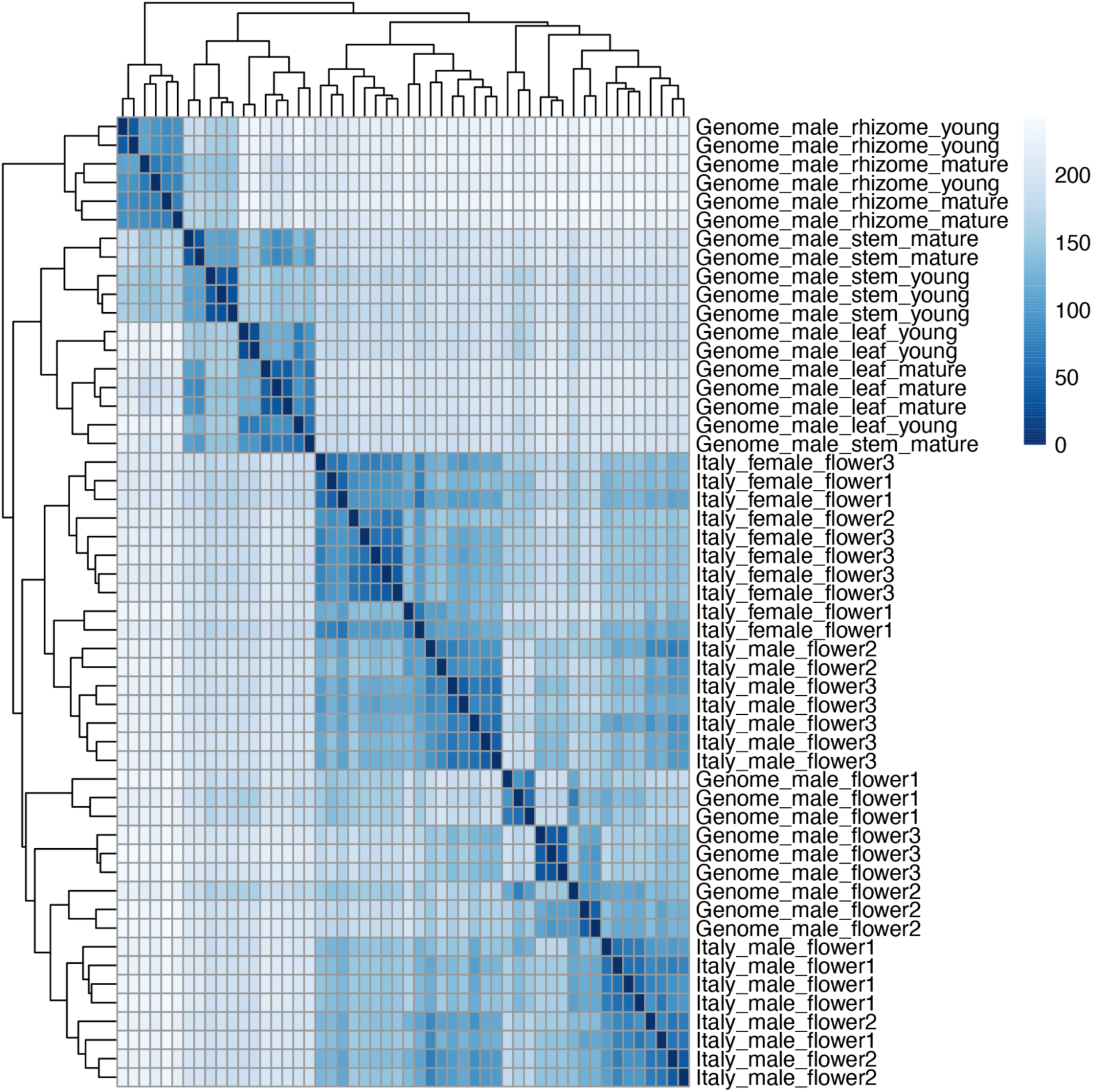
Clustered heat map of expression profiles for each tissue type sampled for transcriptome sequencing for *Asparagus horridus*.

## SUPPLEMENTARY MATERIALS

SupplementalTables.xlsx Spreadsheet with all supplemental data tables (Tables S1-S11) referred to in the main text

SupplementalFile1.pdf Transposable element divergence plot for Asparagus officinalis (pb81m) haplotype 1

SupplementalFile2.pdf Transposable element divergence plot for Asparagus officinalis (pb81m) haplotype 2

SupplementalFile3.pdf Transposable element divergence plot for Asparagus horridus (pb32m) haplotype 1

SupplementalFile4.pdf Transposable element divergence plot for Asparagus horridus (pb32m) haplotype 2

SupplementalFile5.pdf Transposable element annotation density plots for Asparagus officinalis (pb81m) haplotype 1

SupplementalFile6.pdf Transposable element annotation density plots for Asparagus officinalis (pb81m) haplotype 2

SupplementalFile7.pdf Transposable element annotation density plots for Asparagus horridus (pb32m) haplotype 1

SupplementalFile8.pdf Transposable element annotation density plots for Asparagus horridus (pb32m) haplotype 2

SupplementalFile9.pdf Riparian plot of gene synteny plot among all new Asparagus haplotypes/genomes from this study

SupplementalFile10.pdf Phylogenetic analysis (gene tree inferred from amino acid alignments) of the DUF247 gene family across Asparagaceae

