## Supplementary figures and images for "Two independent origins of XY sex chromosomes in *Asparagus*"

### SupplementalFile1

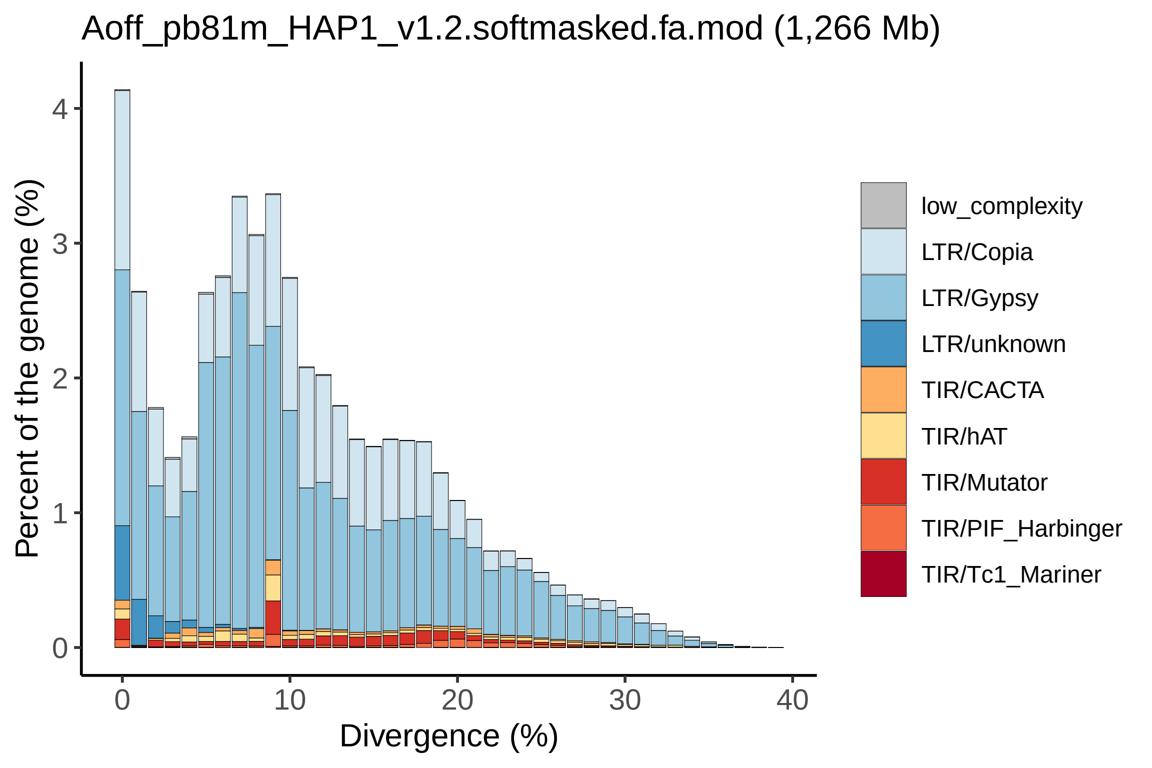

### SupplementalFile2

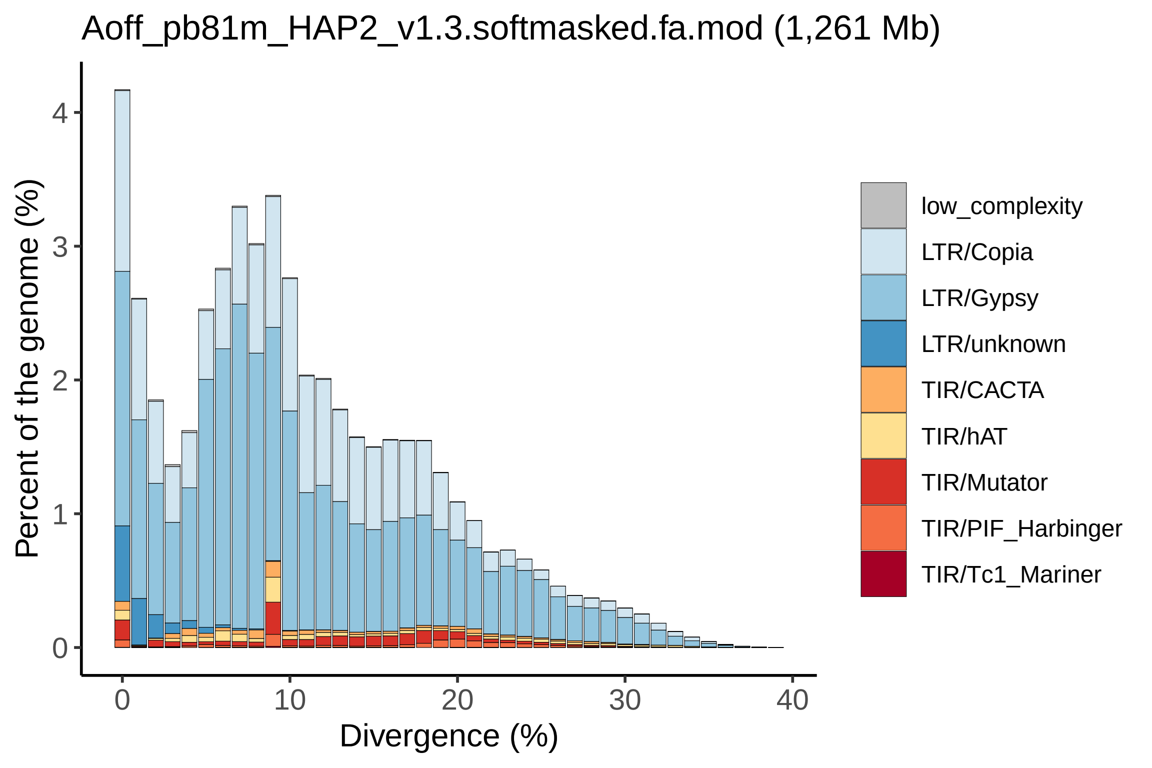

### SupplementalFile3

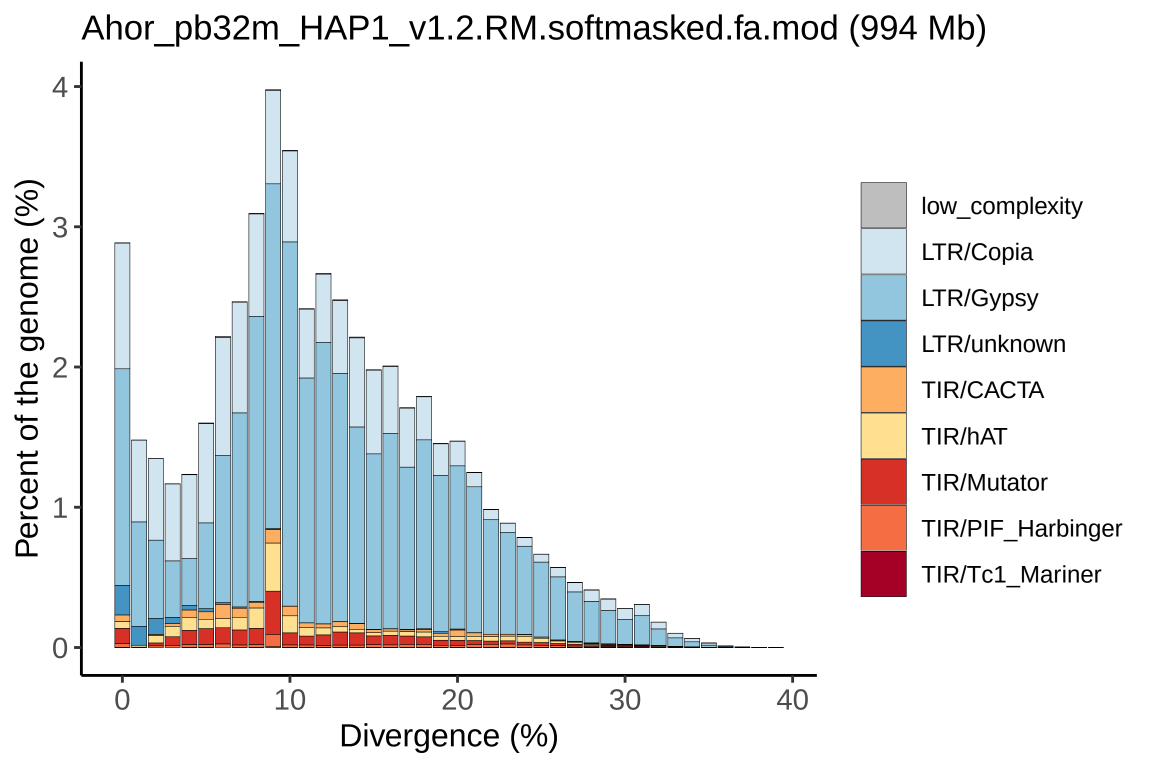

### SupplementalFile4

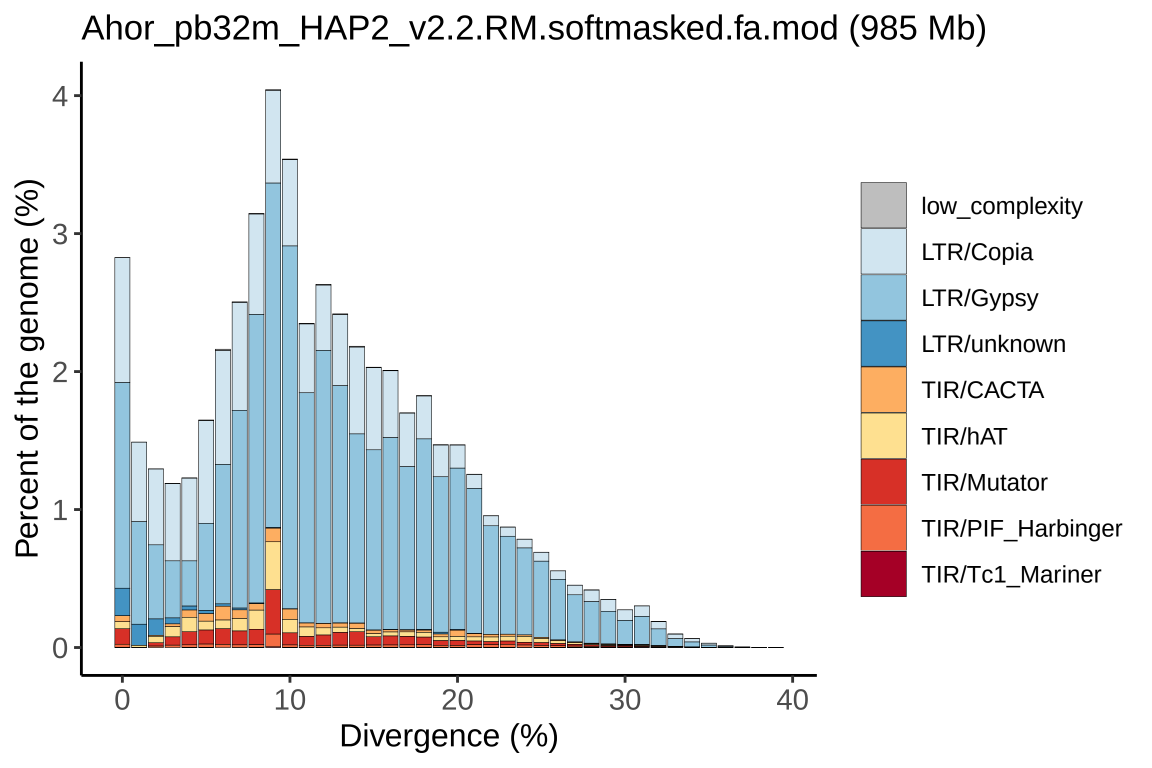

### SupplementalFile5

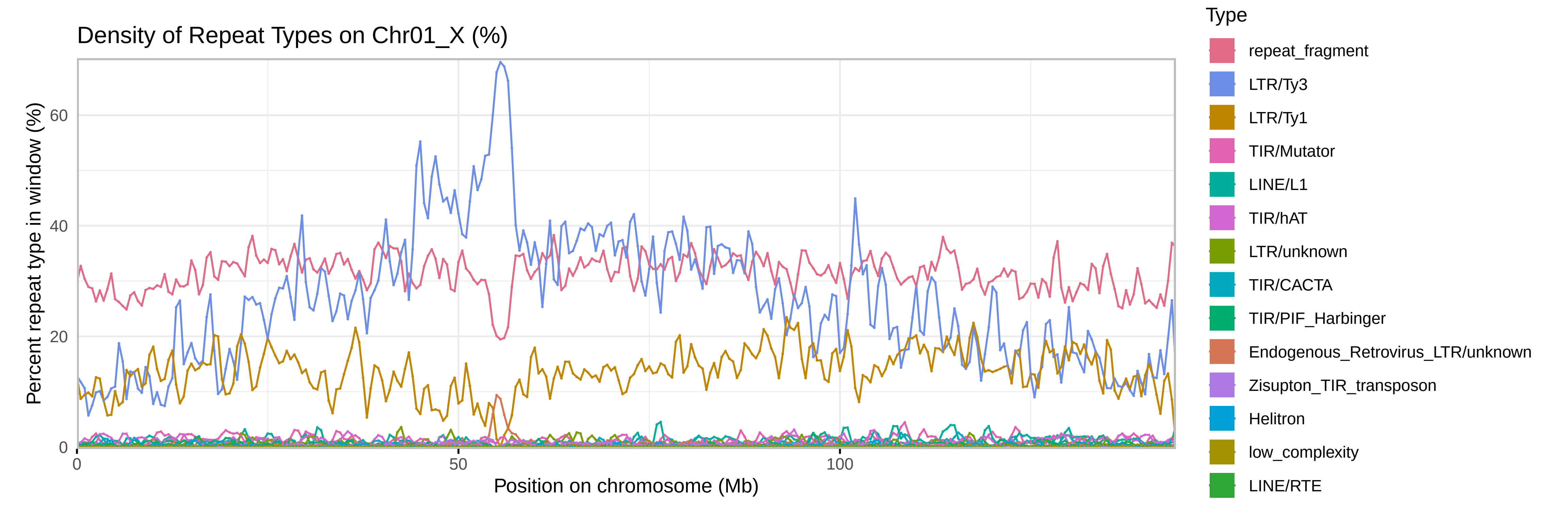

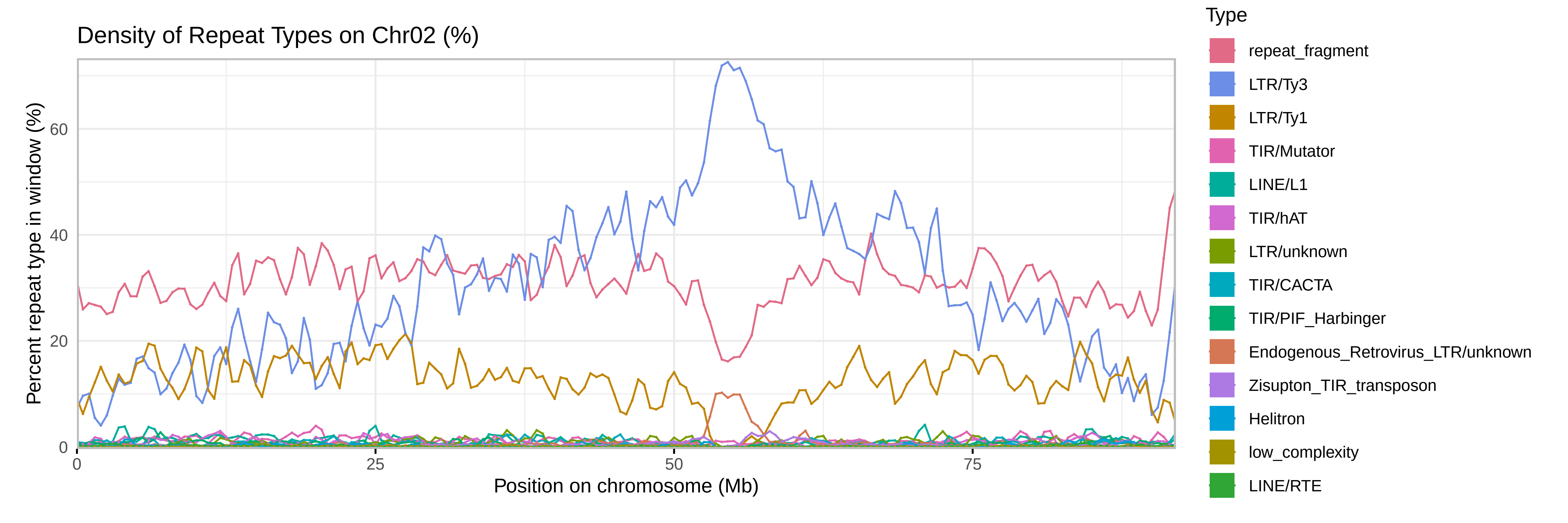

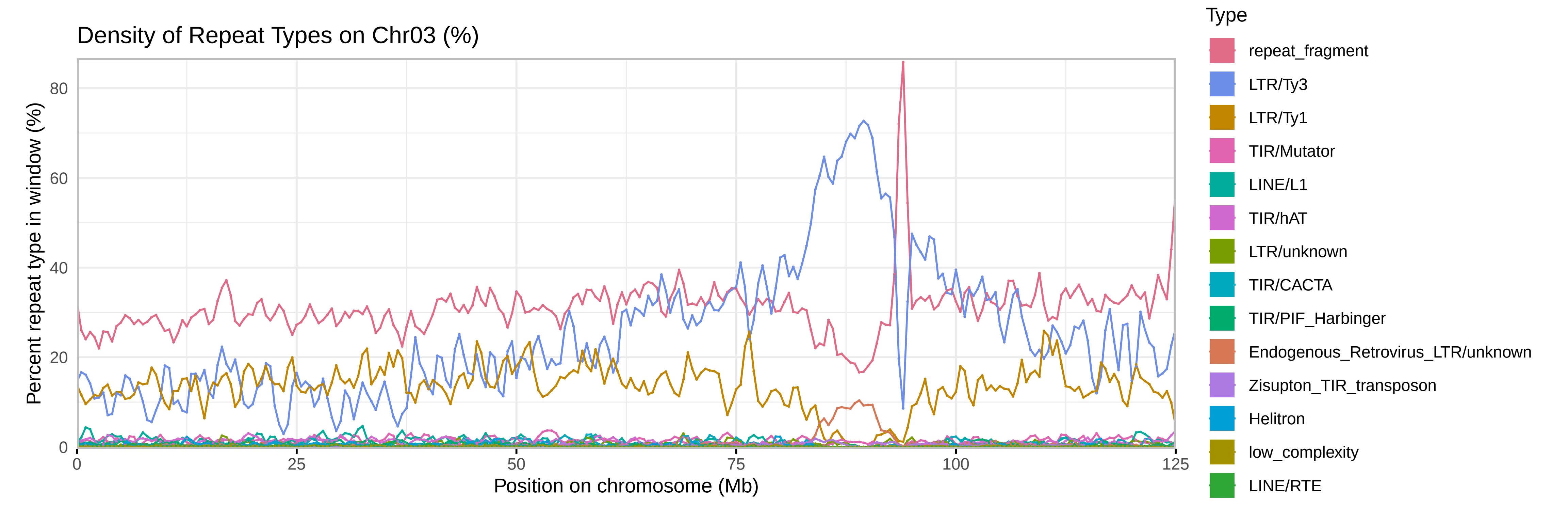

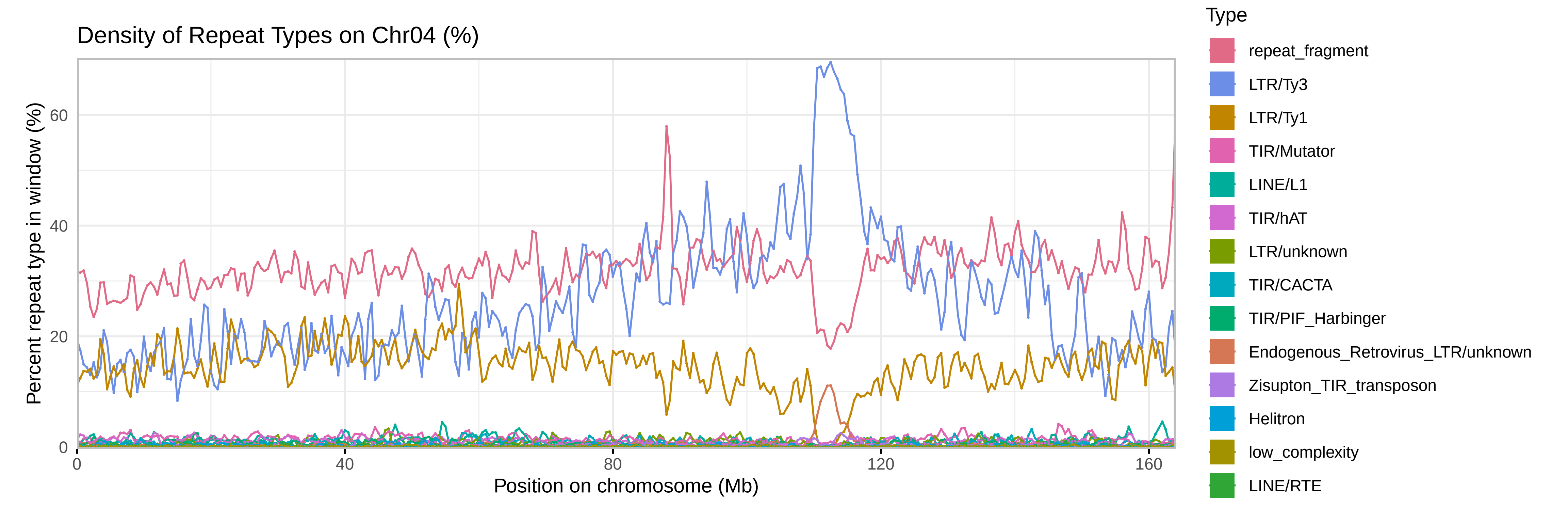

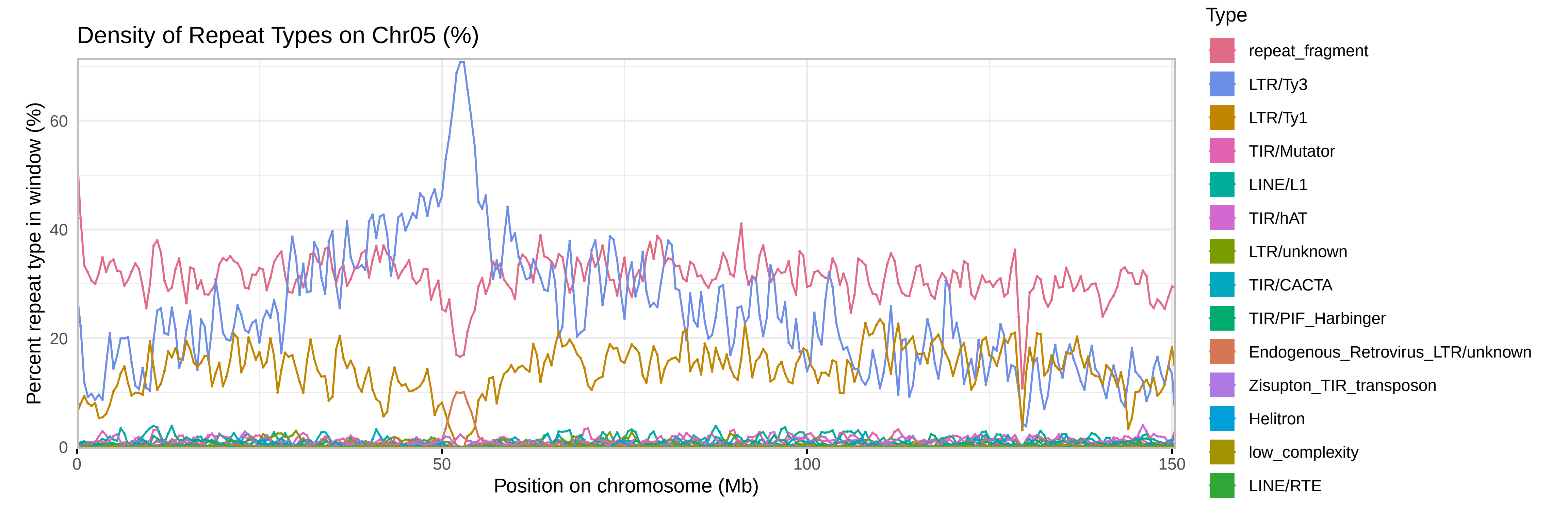

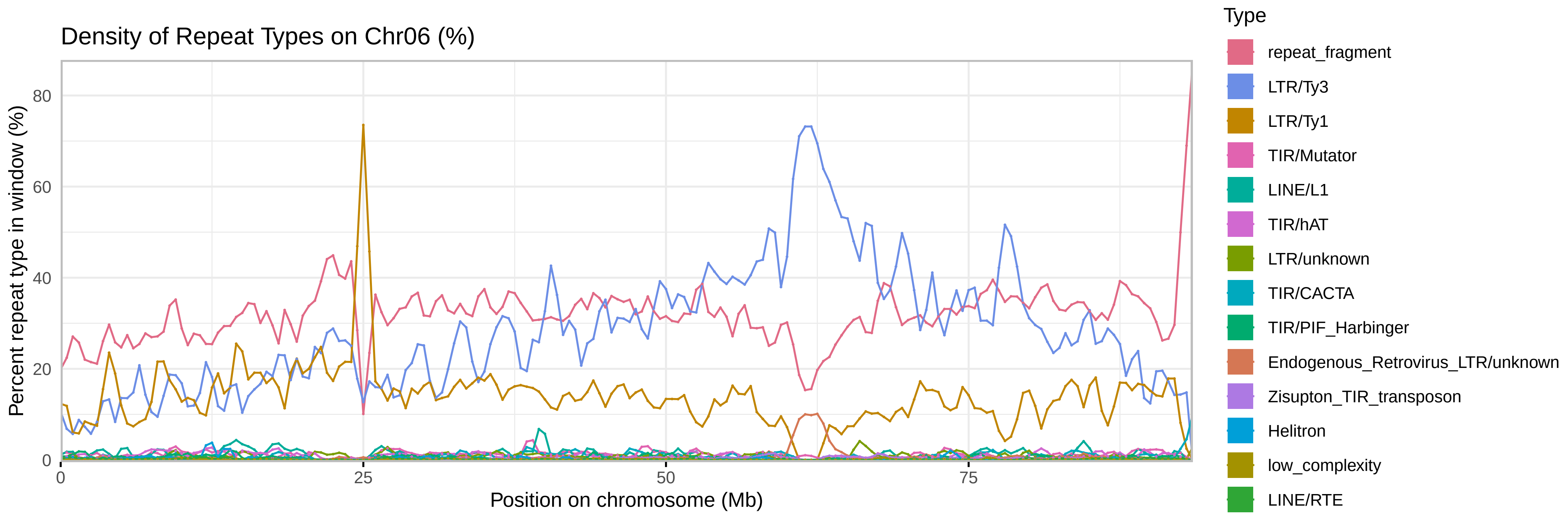

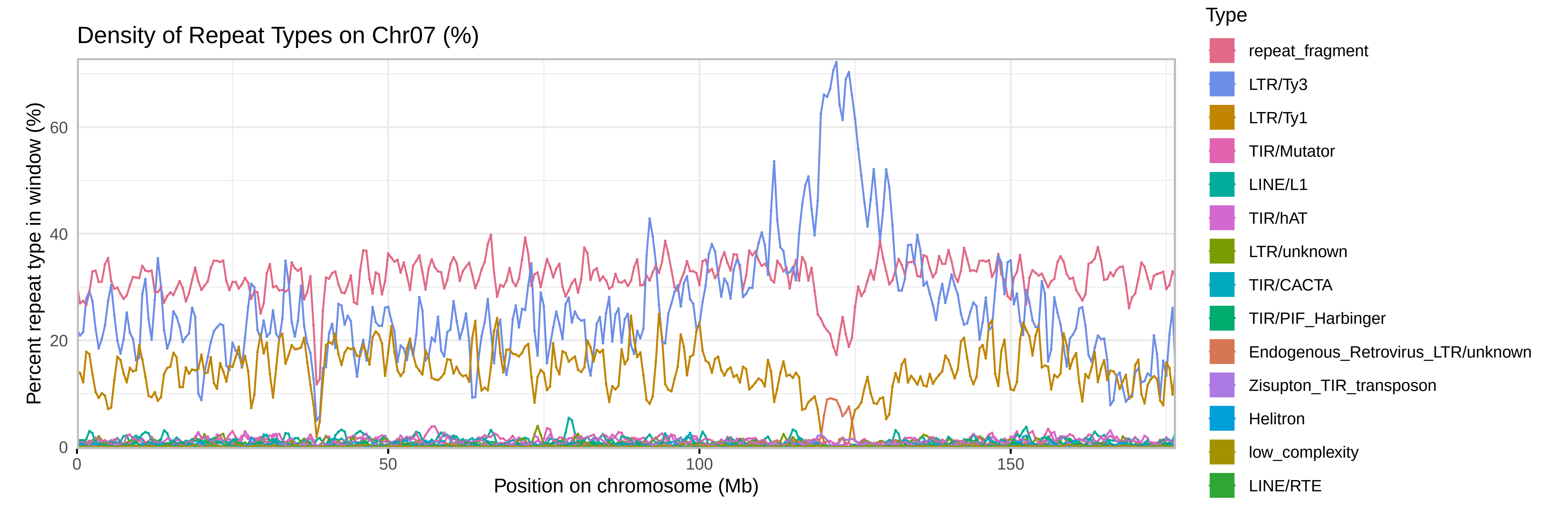

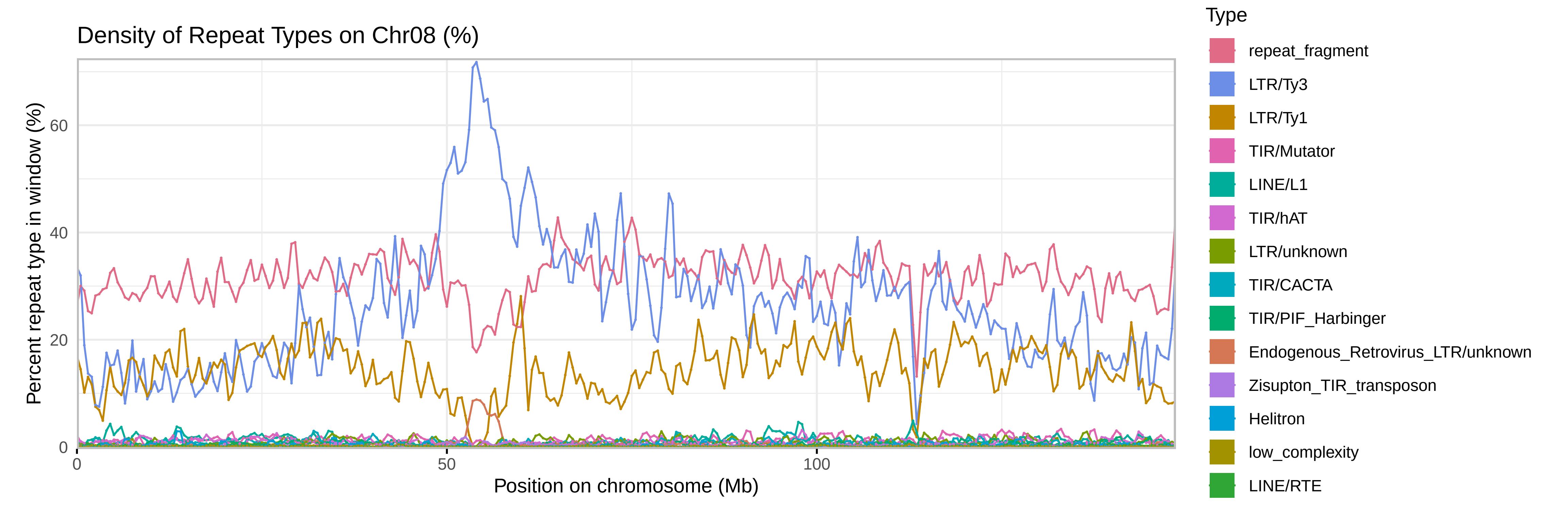

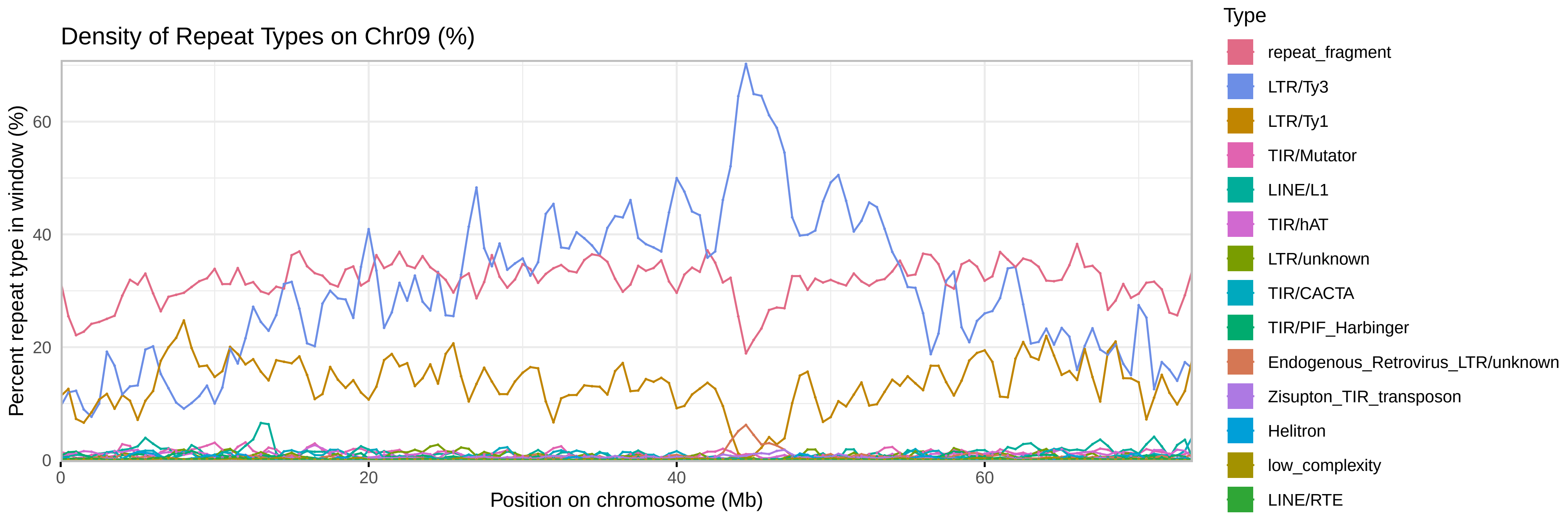

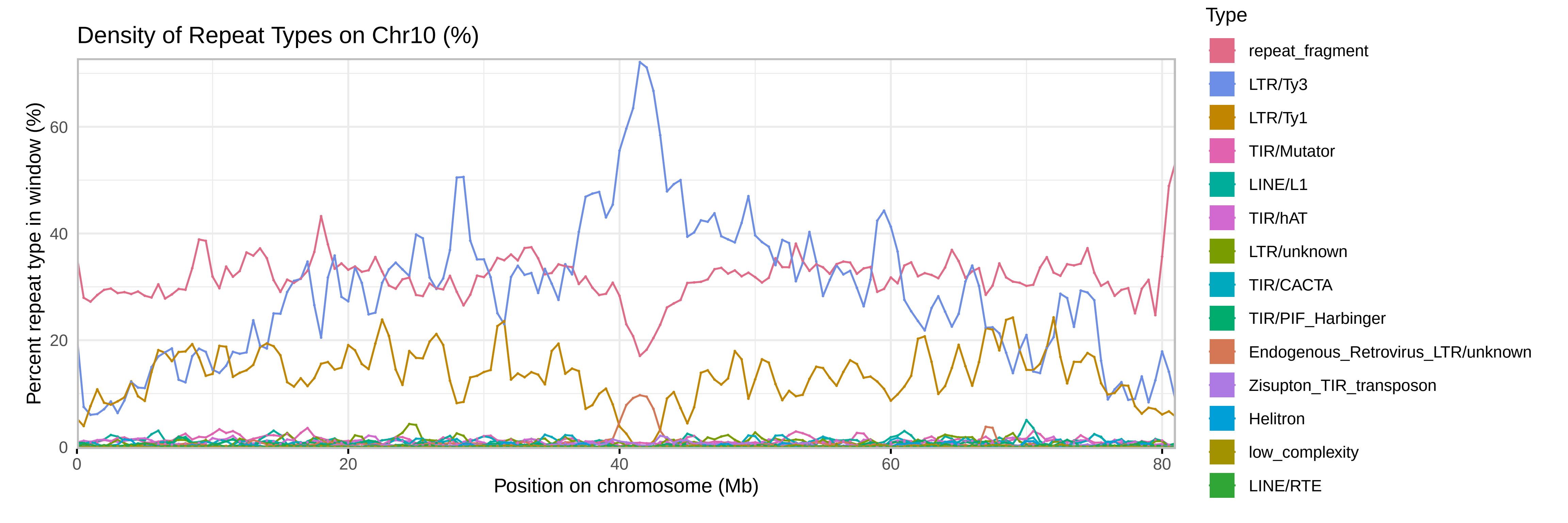

### SupplementalFile6

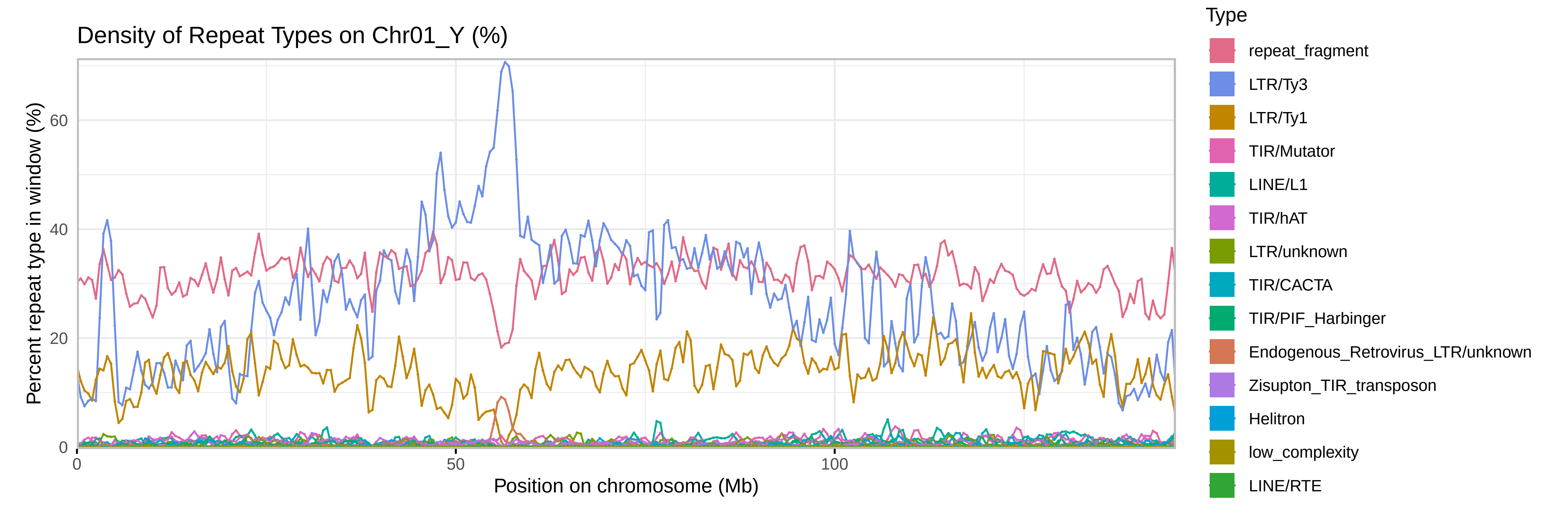

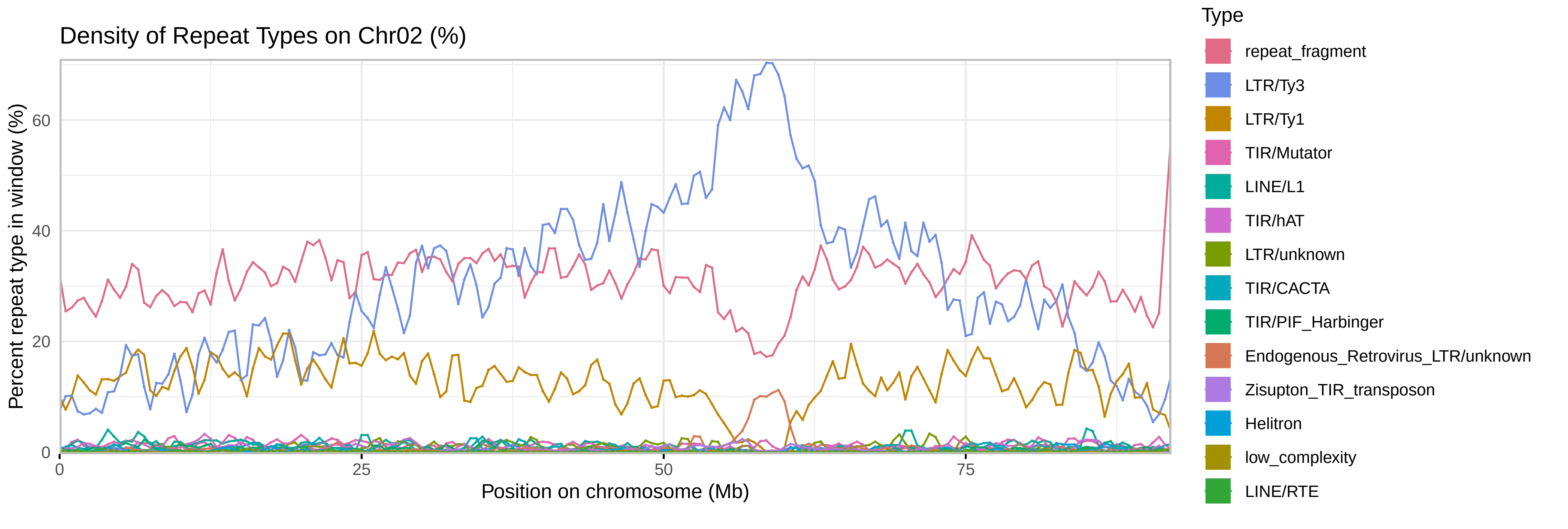

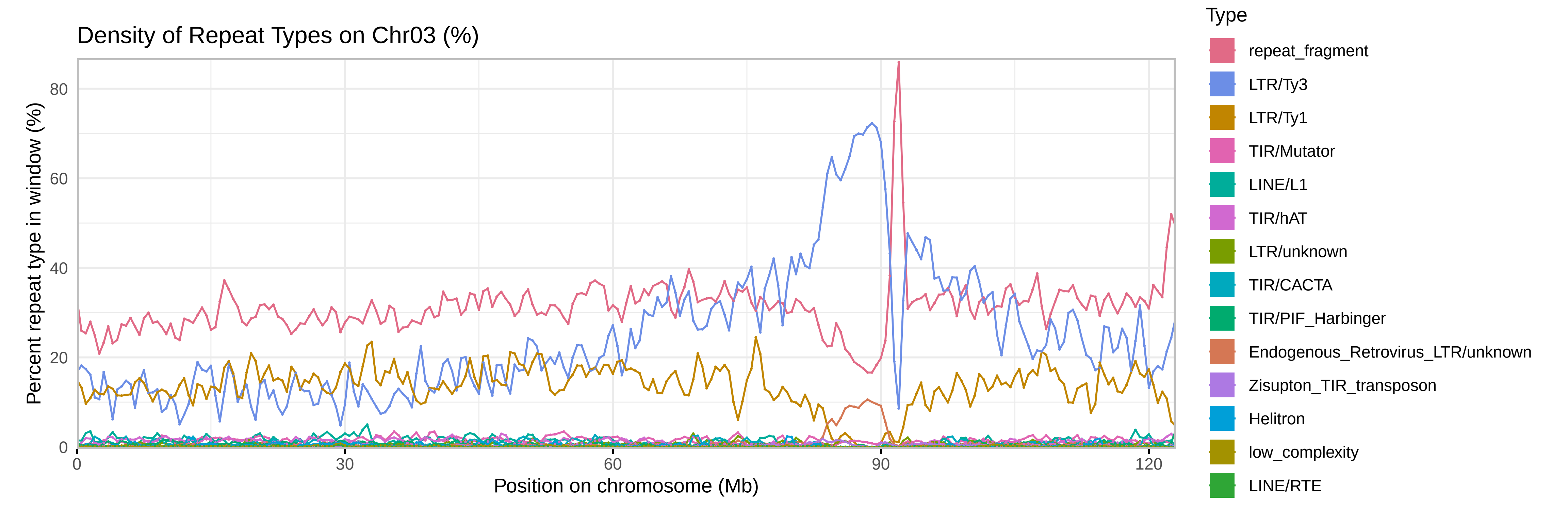

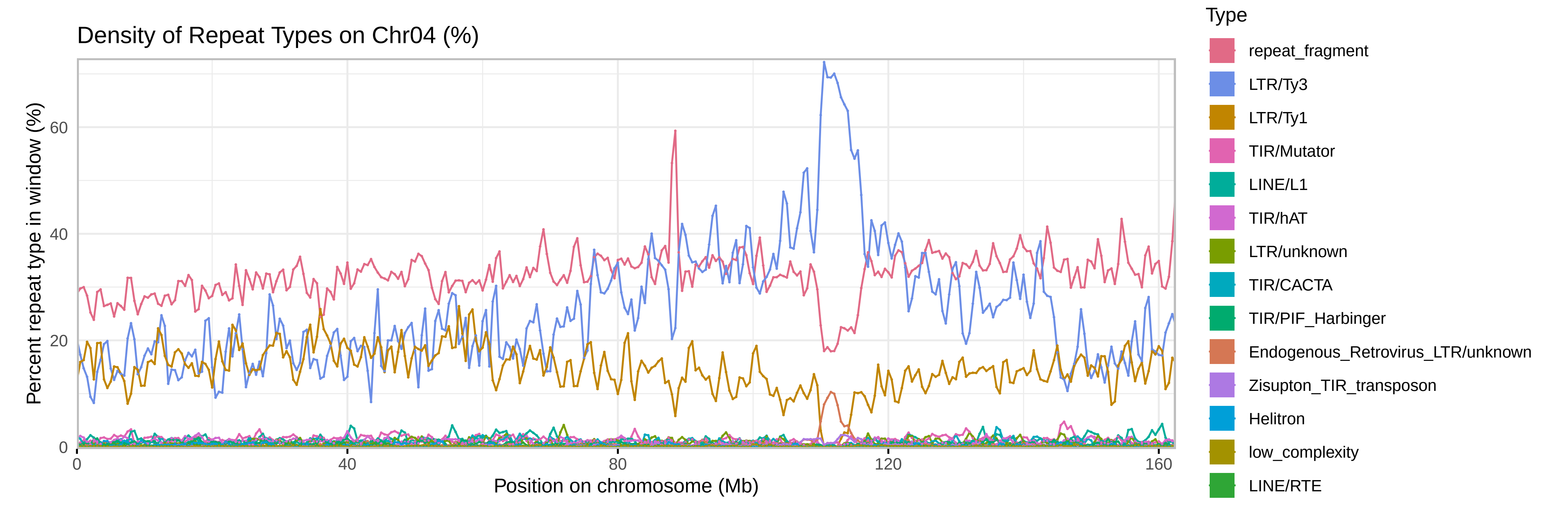

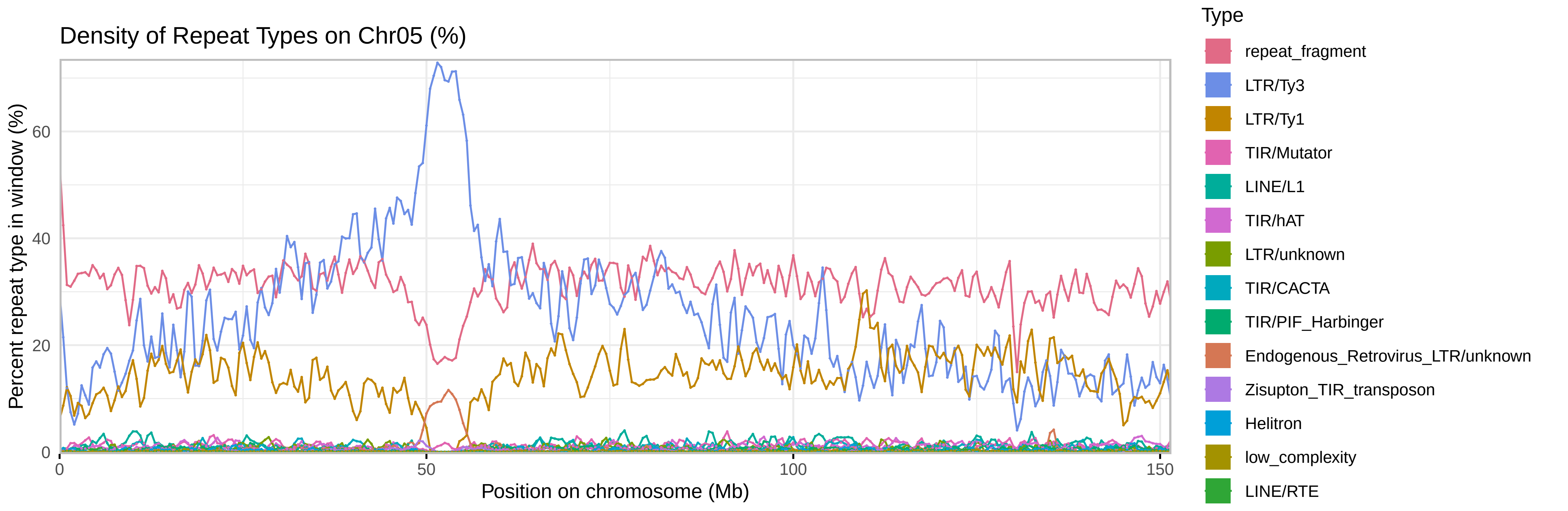

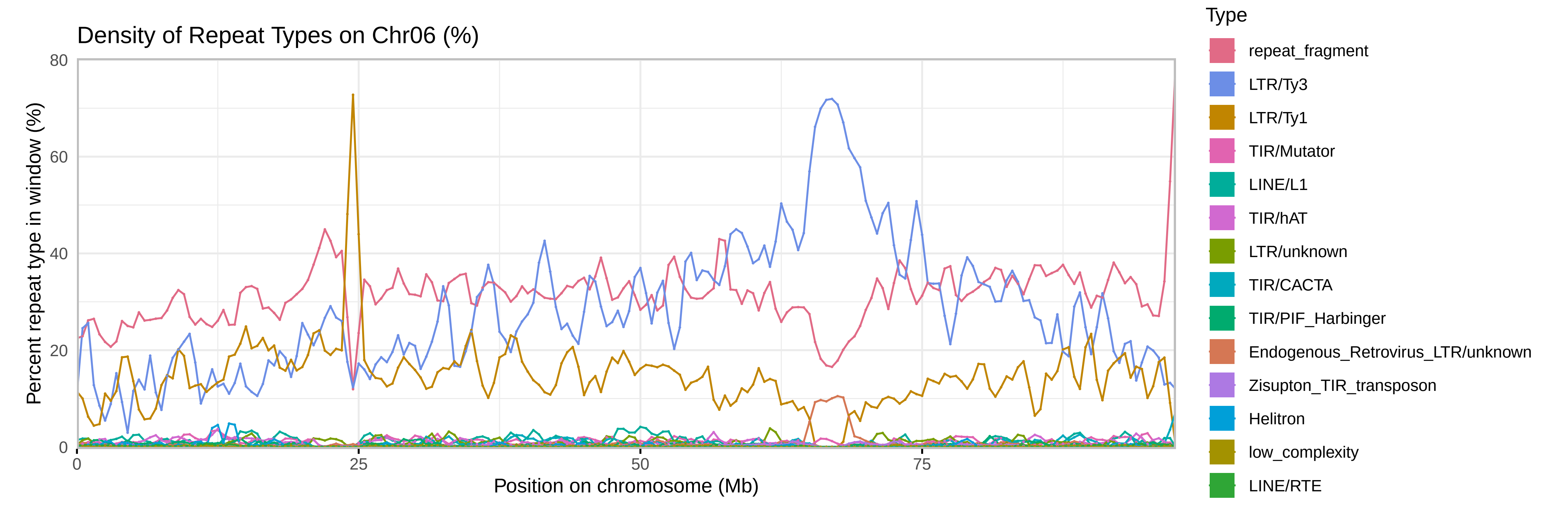

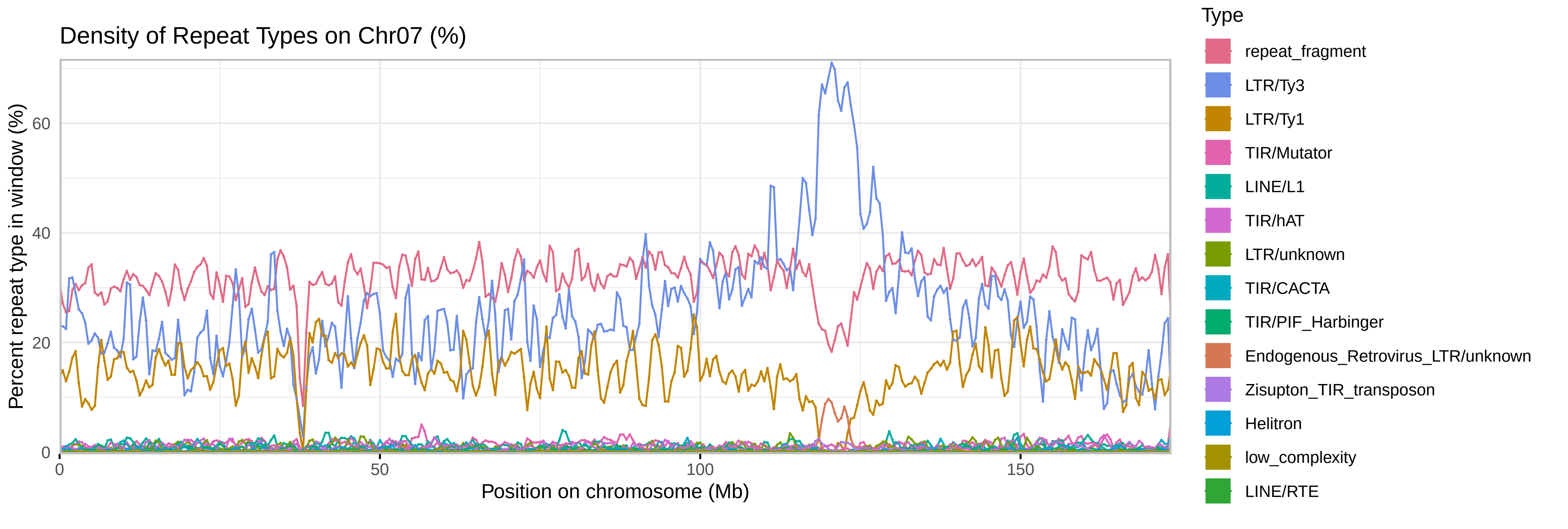

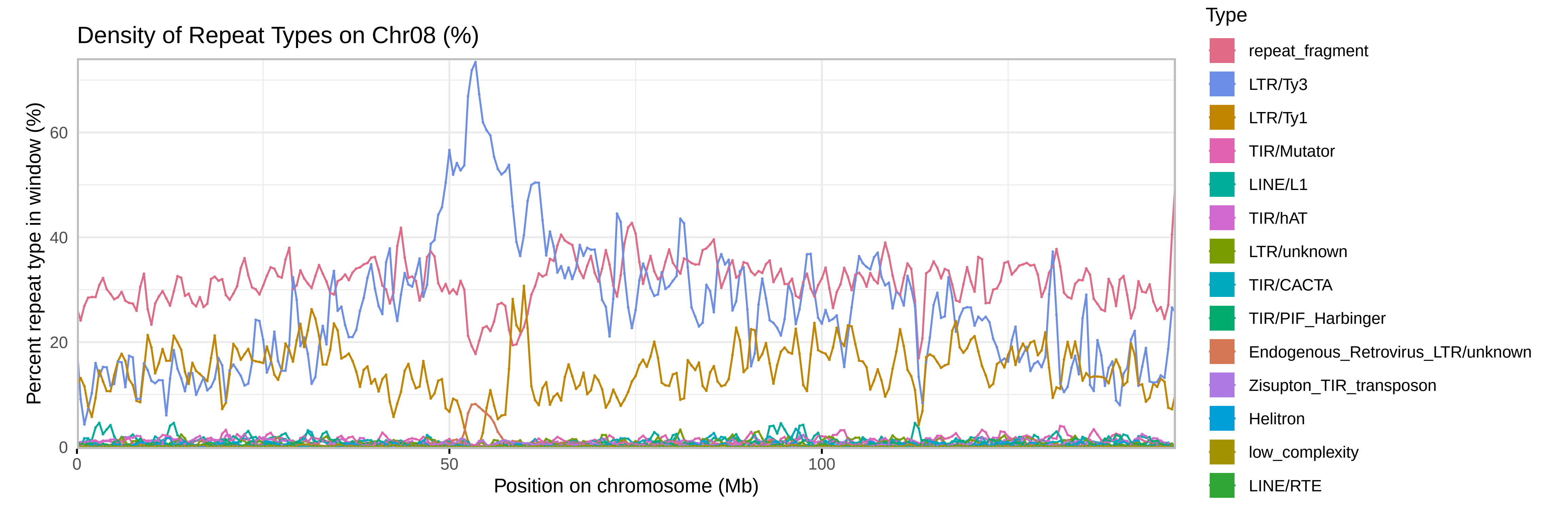

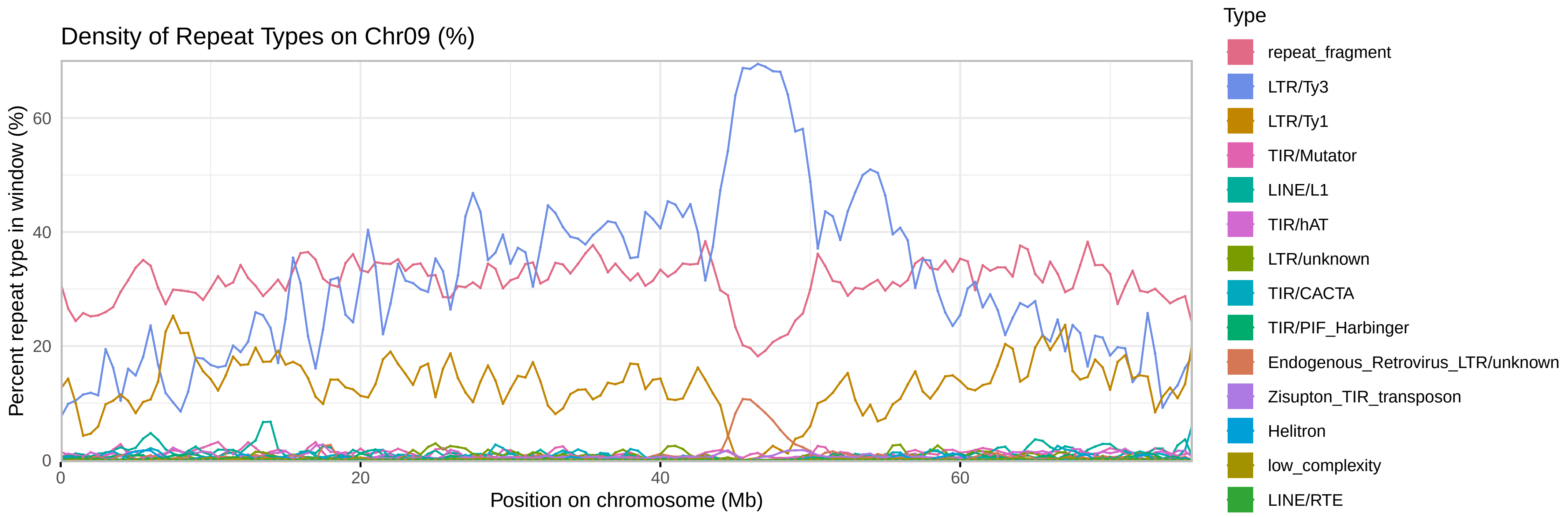

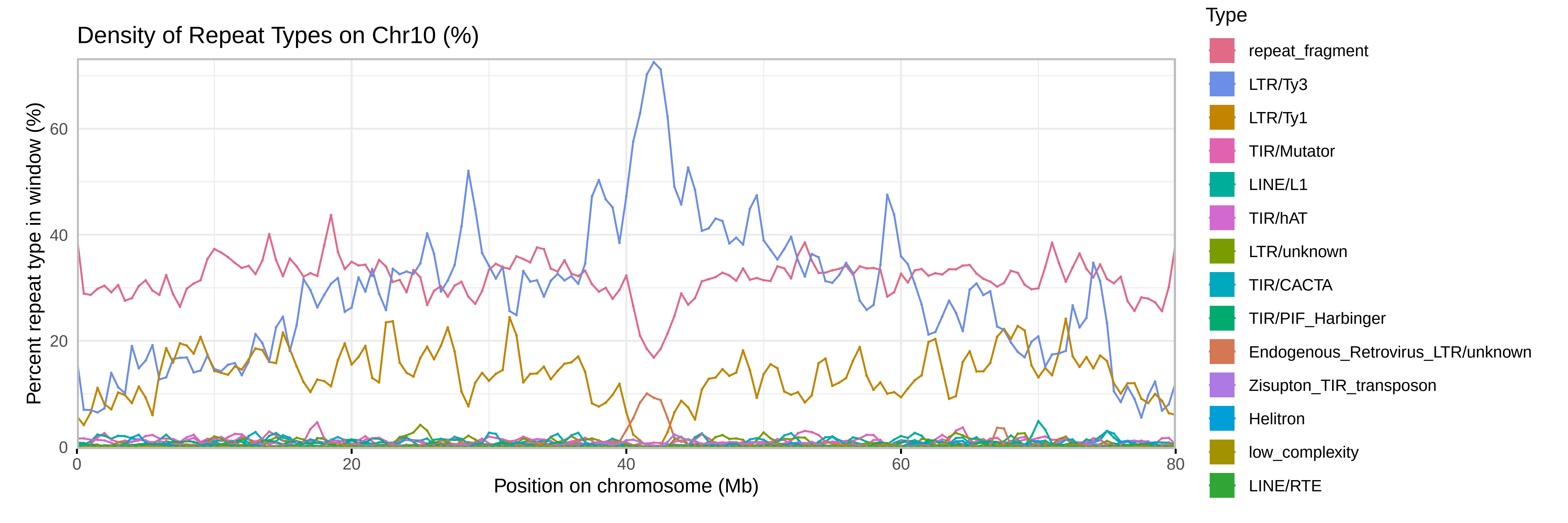

### SupplementalFile7

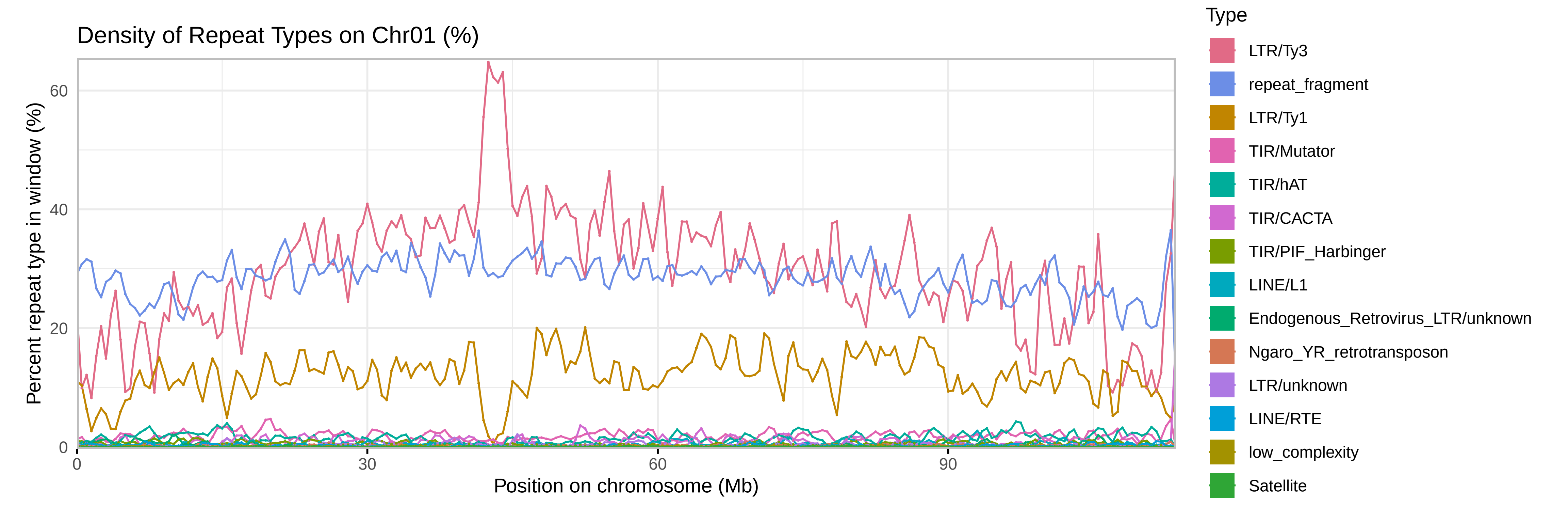

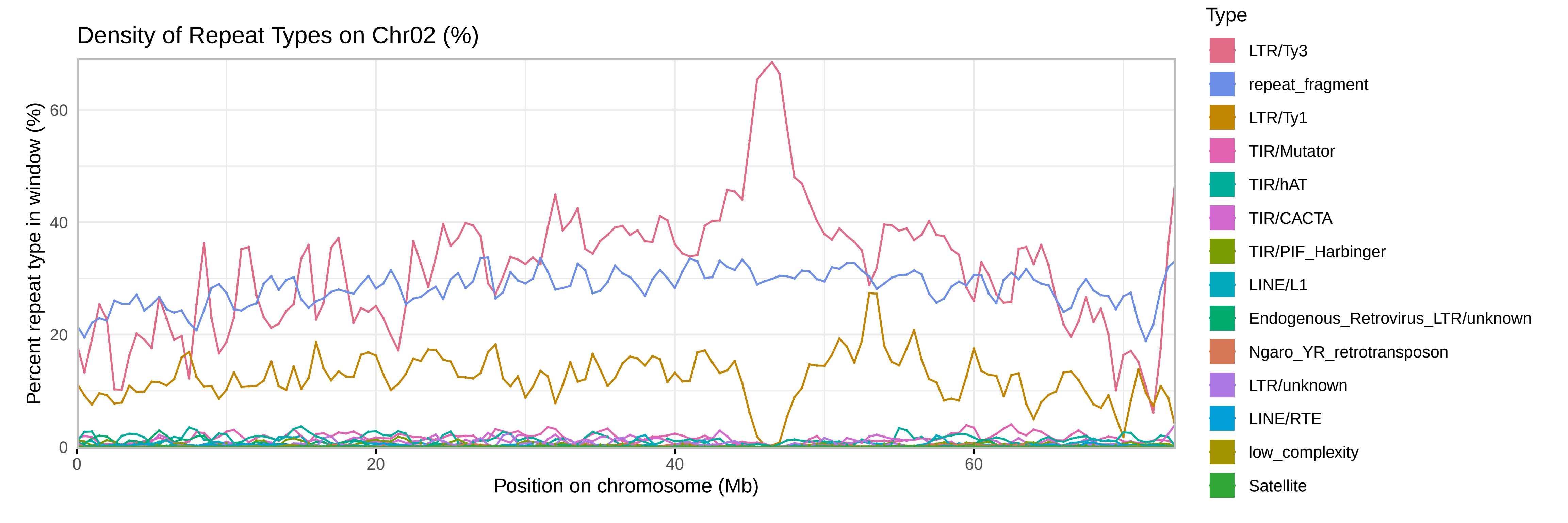

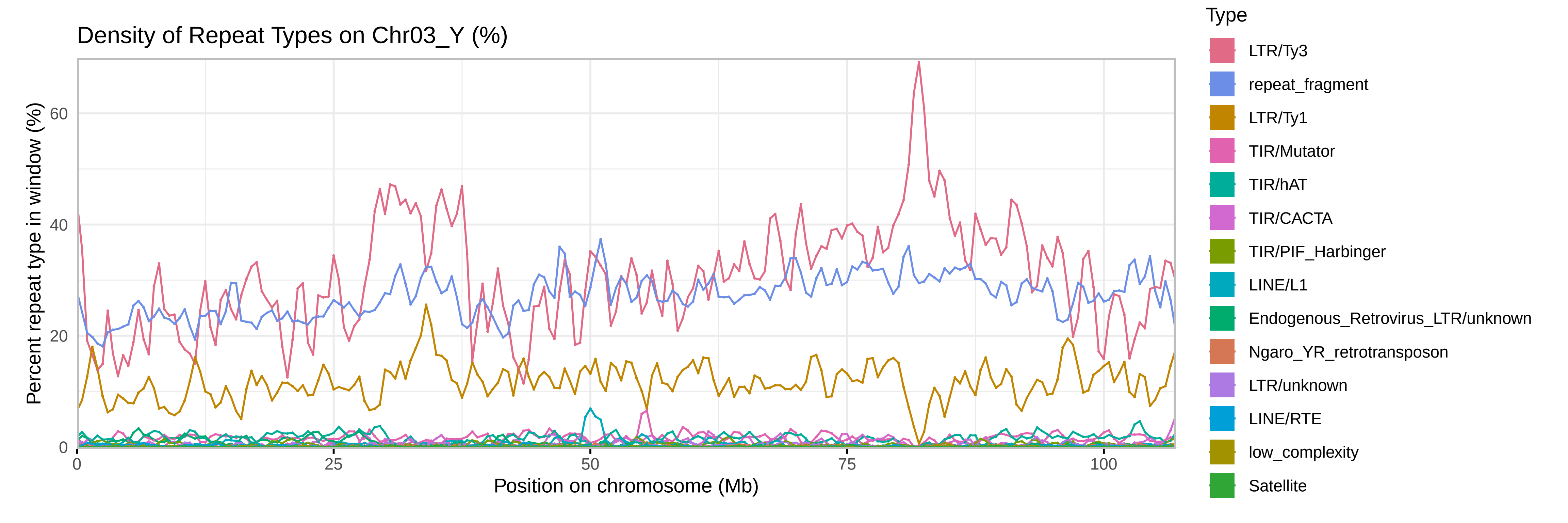

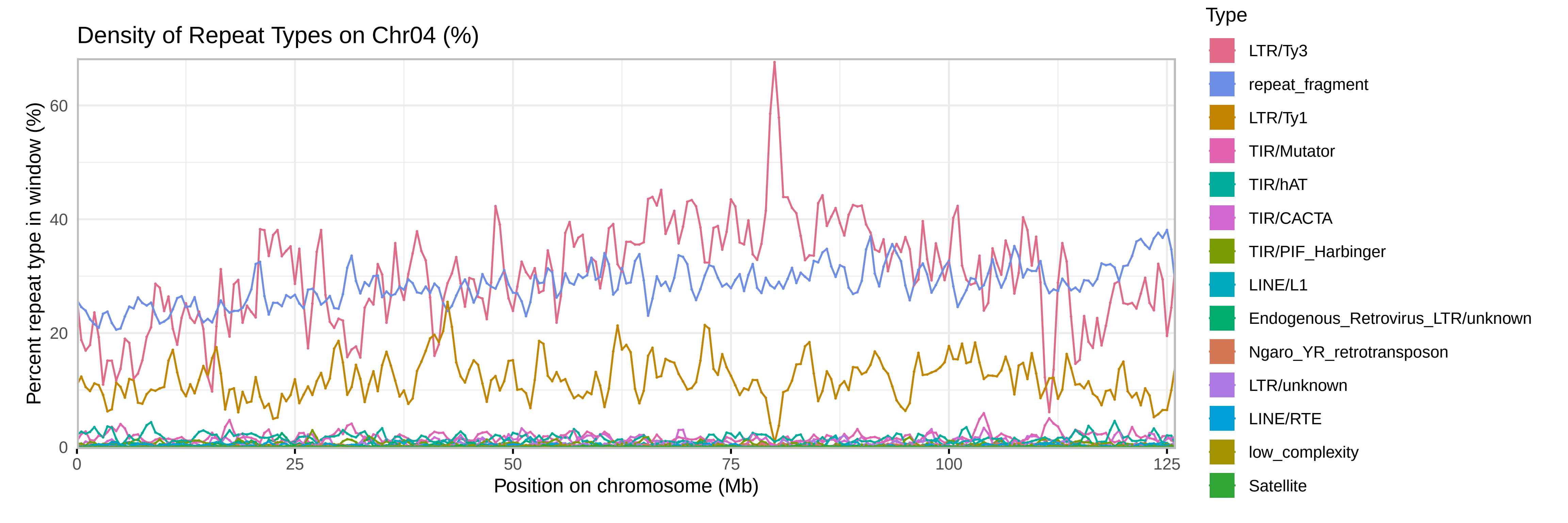

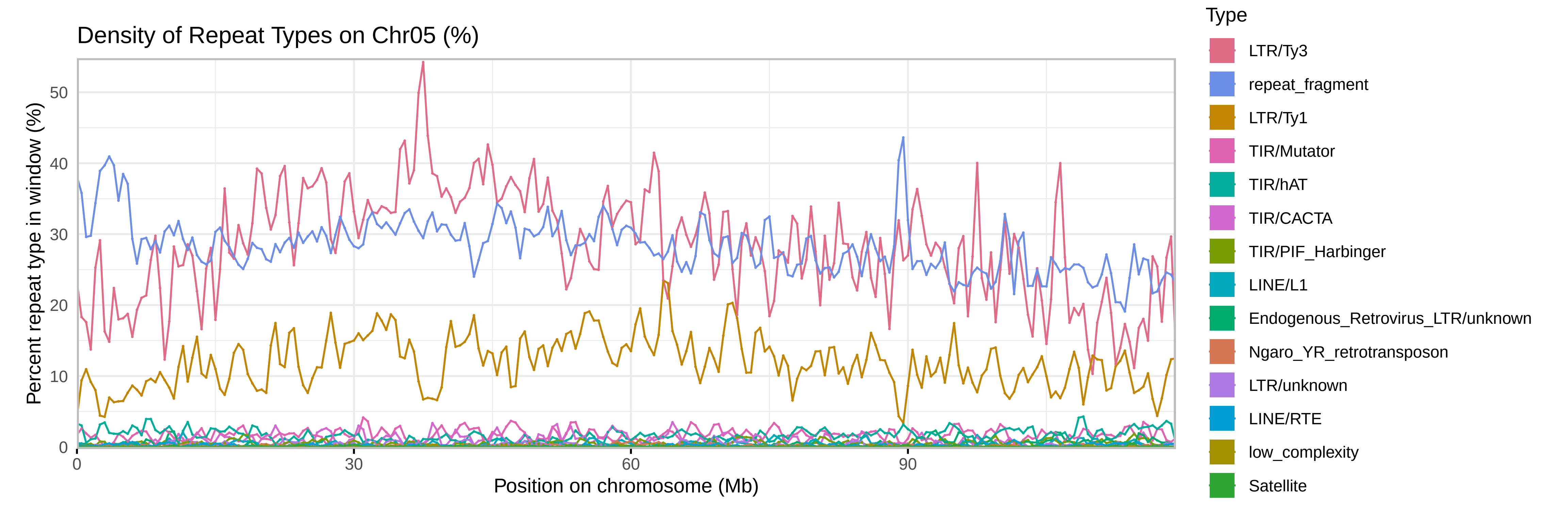

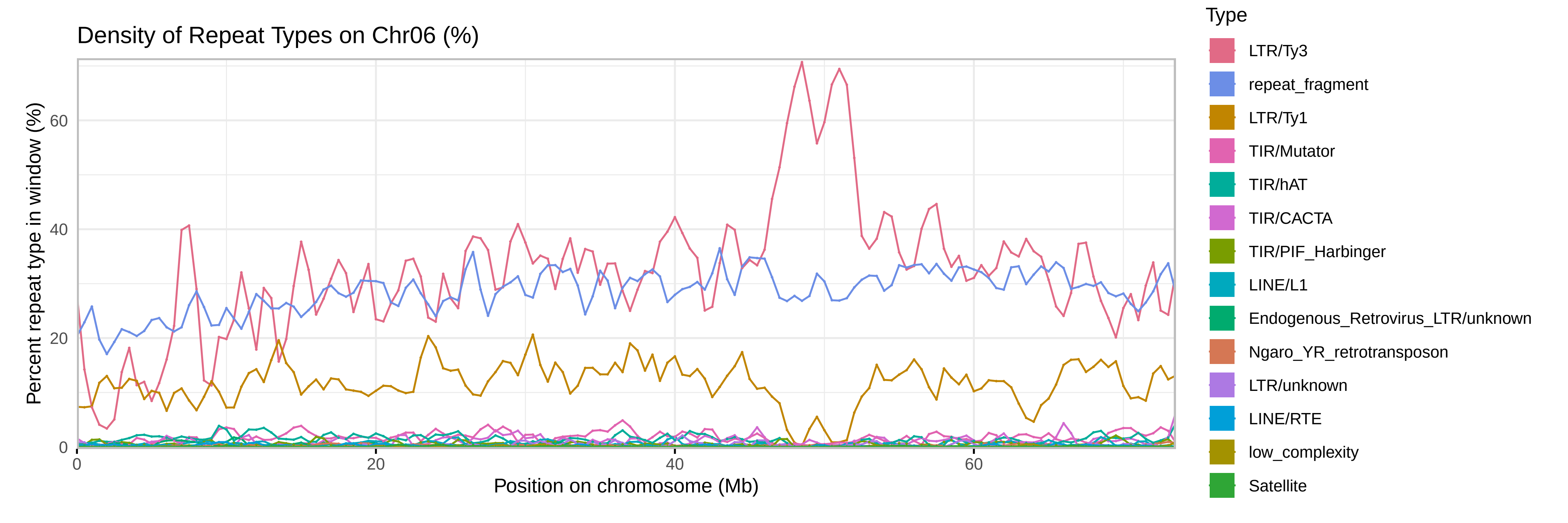

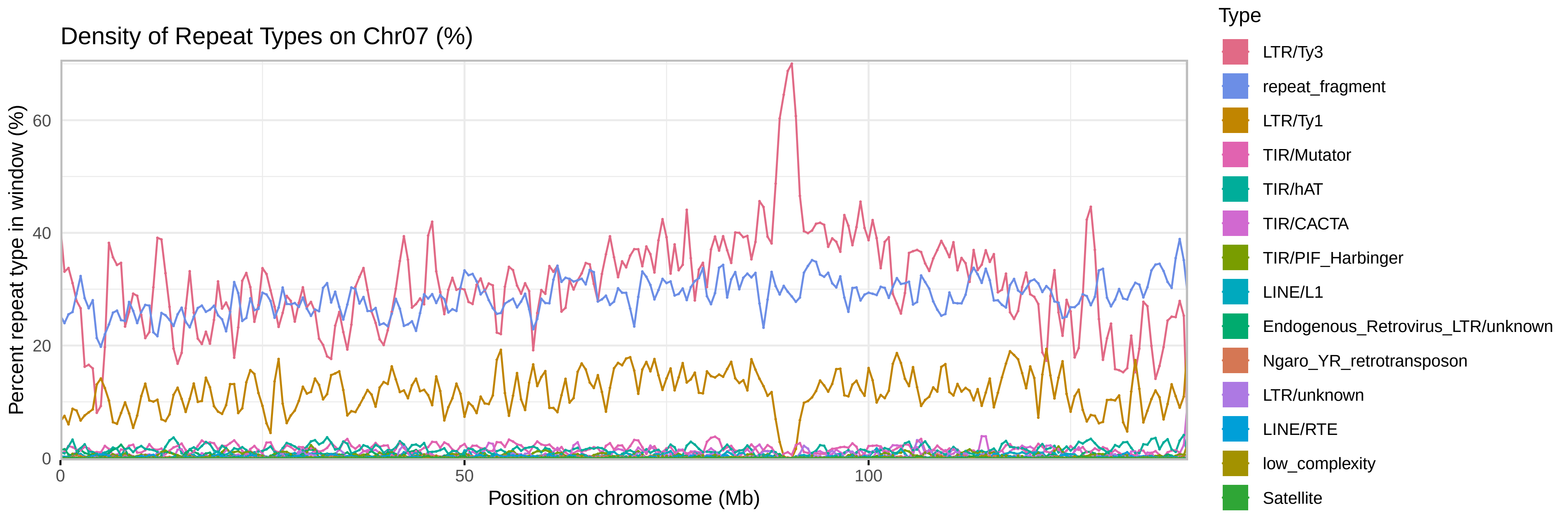

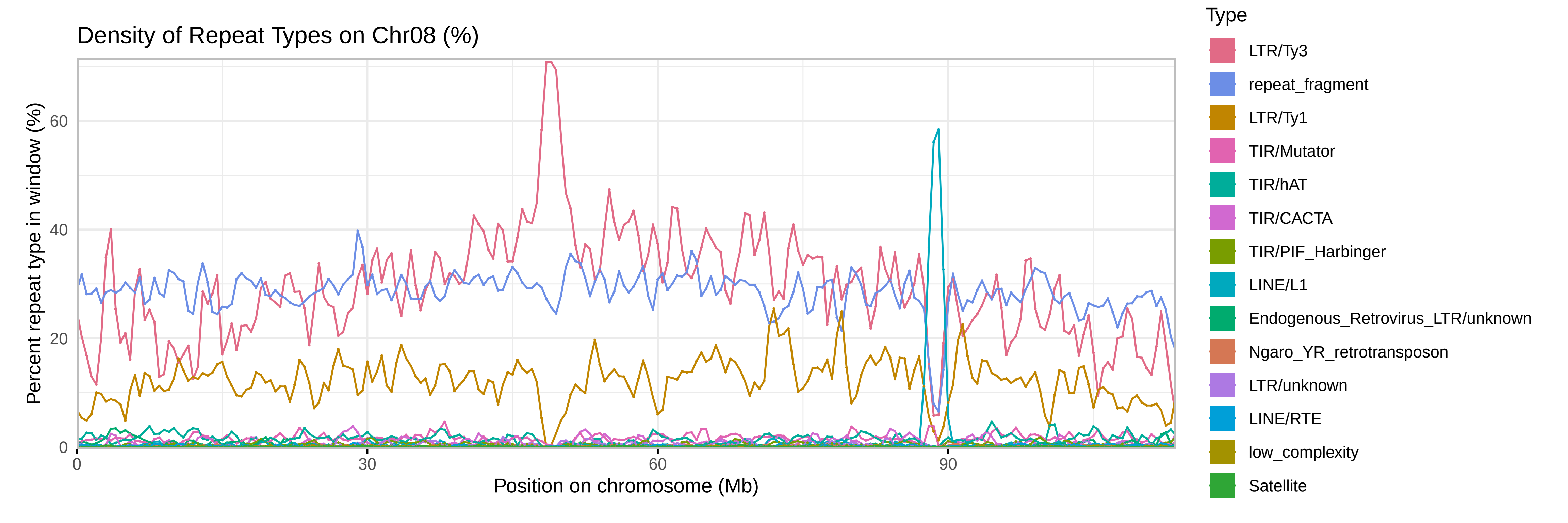

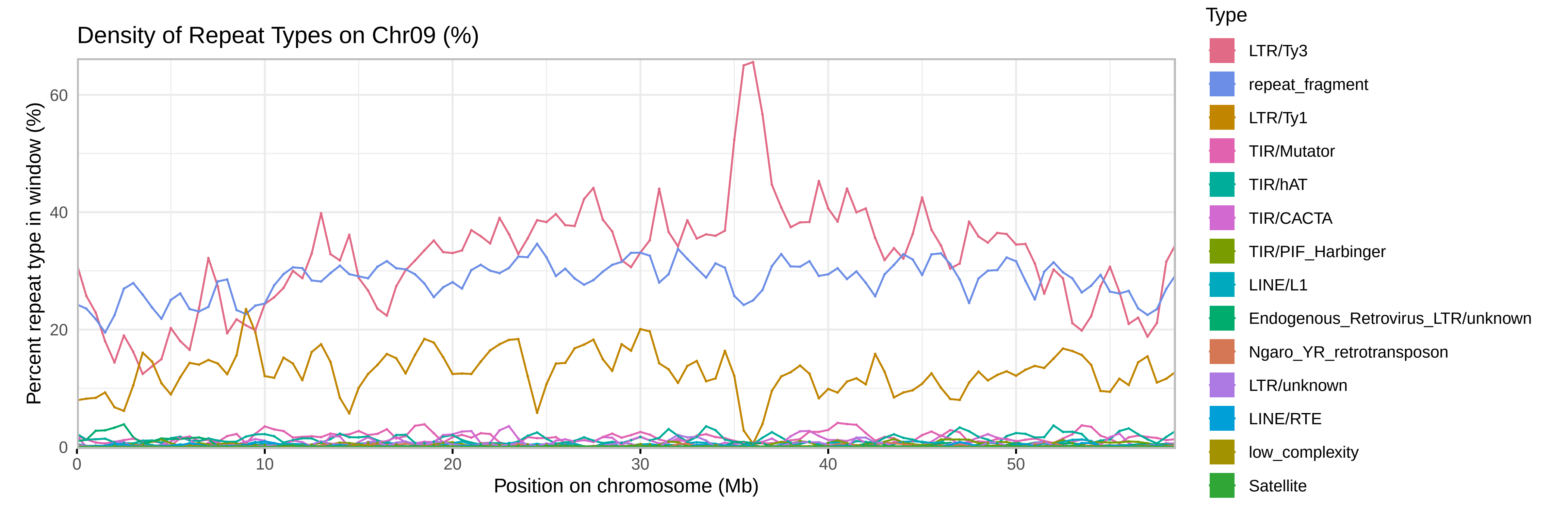

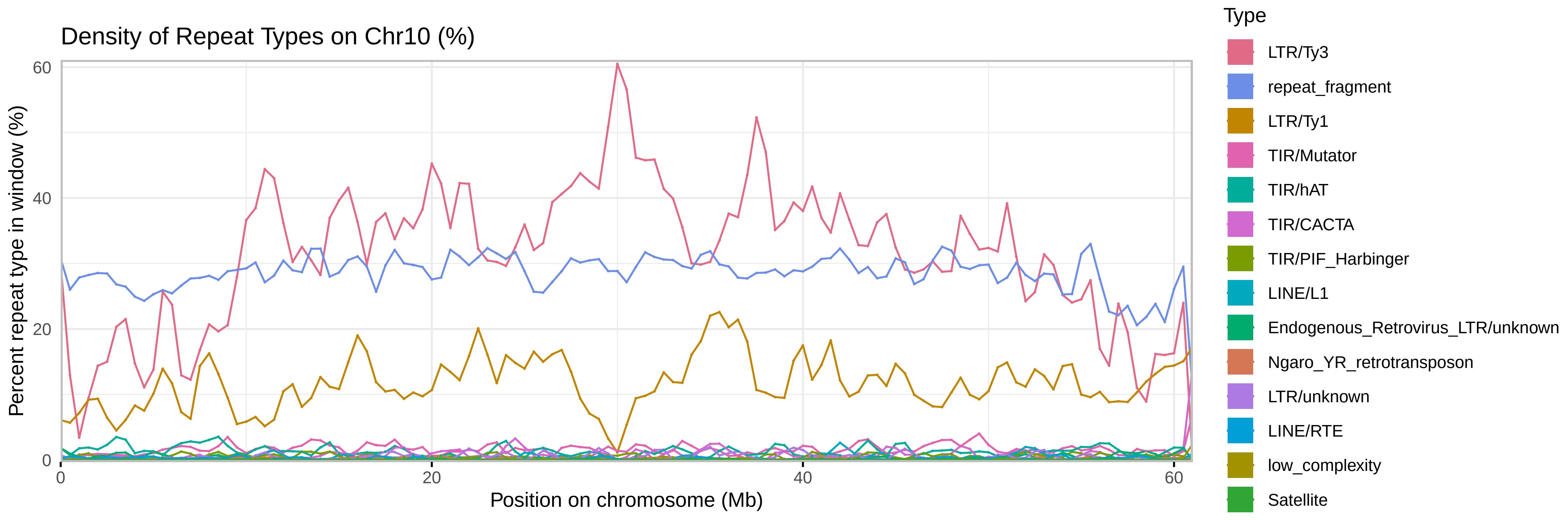

### SupplementalFile8

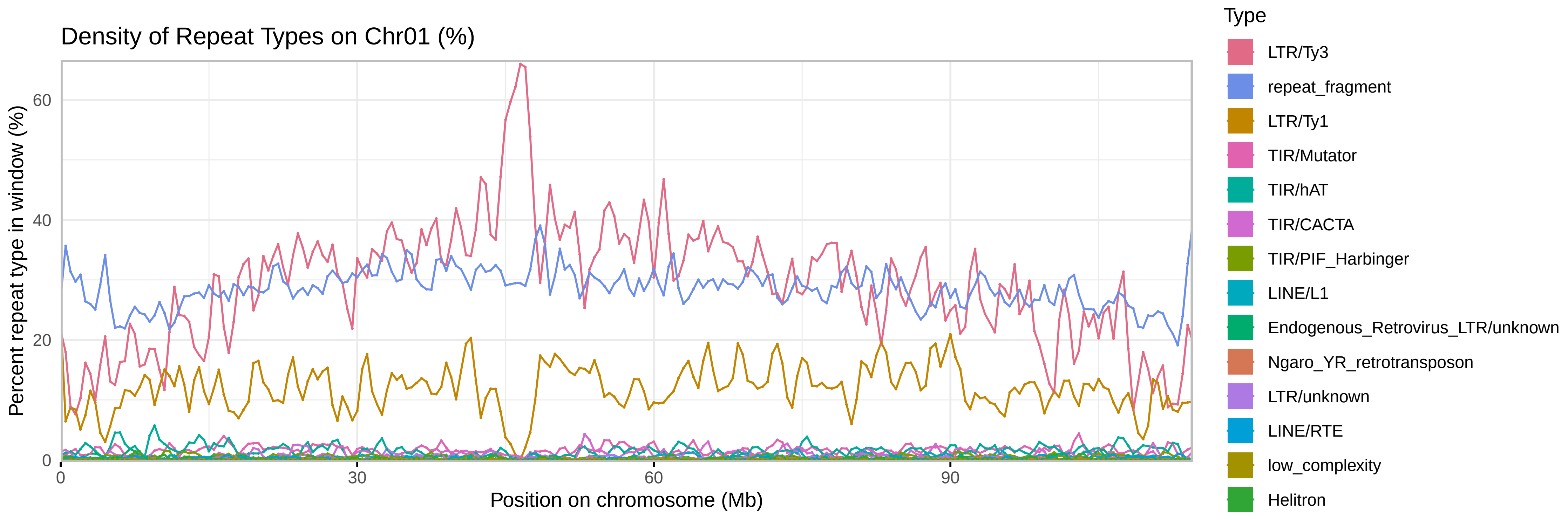

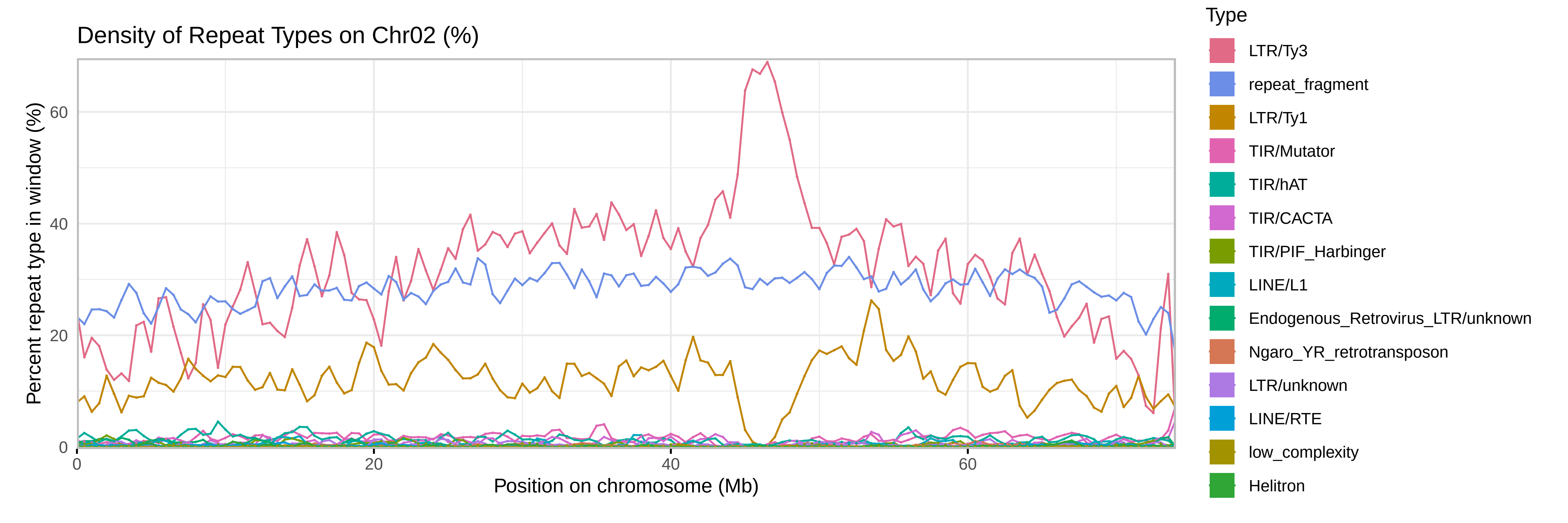

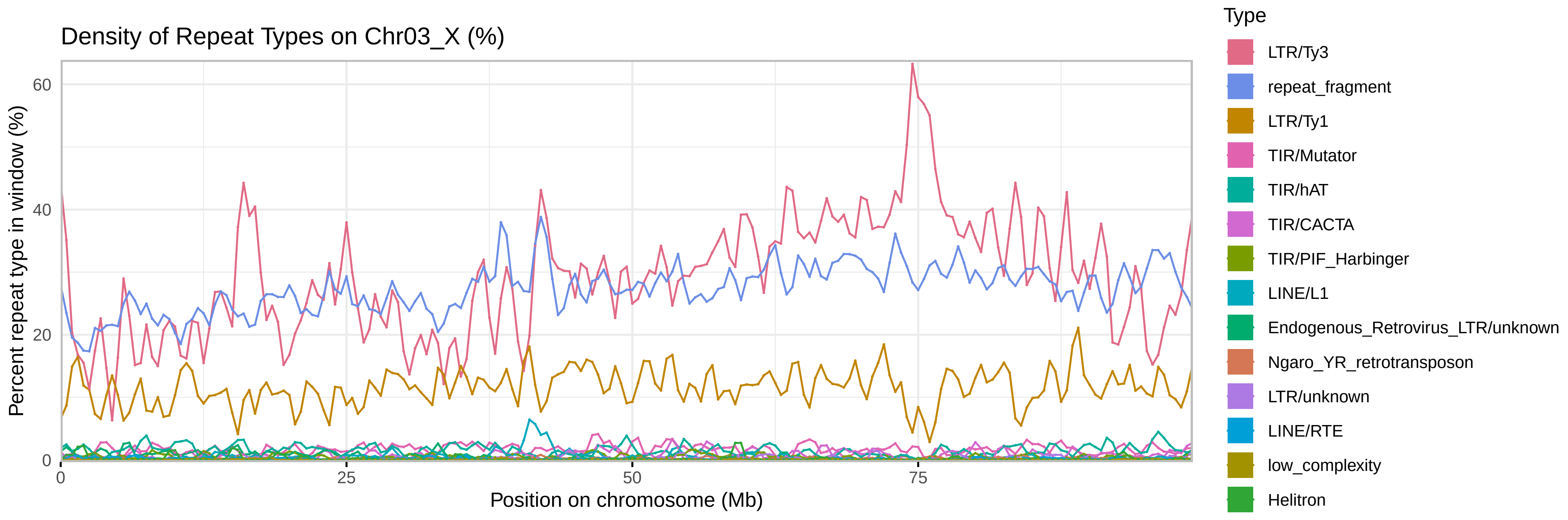

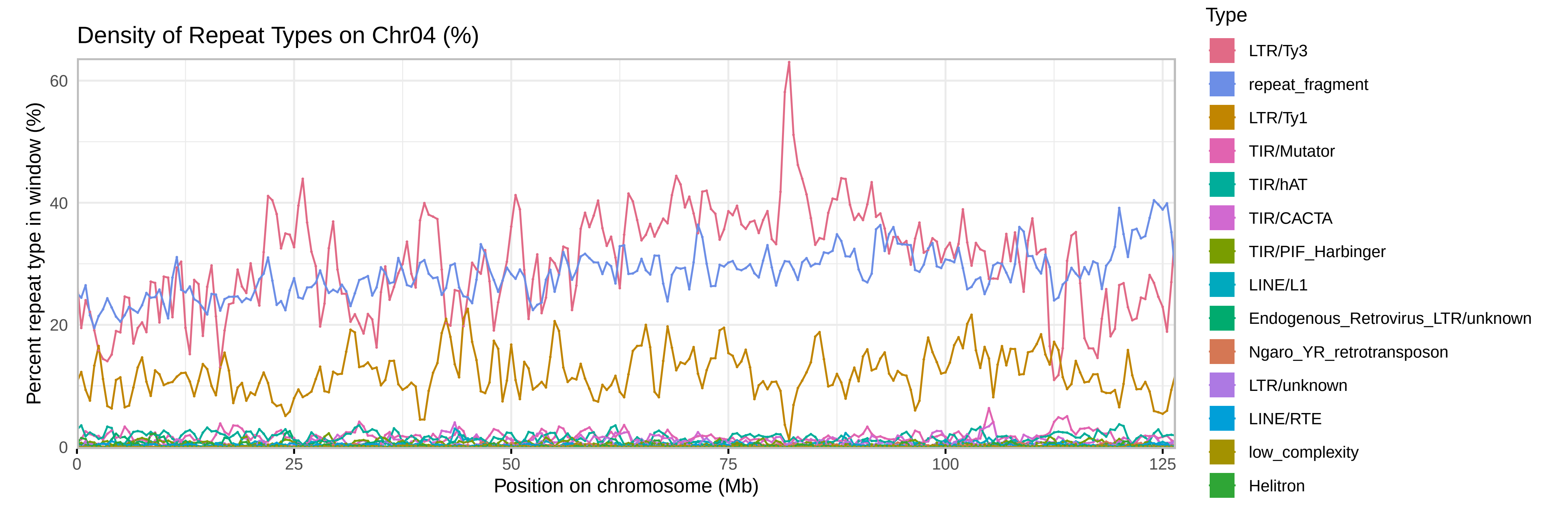

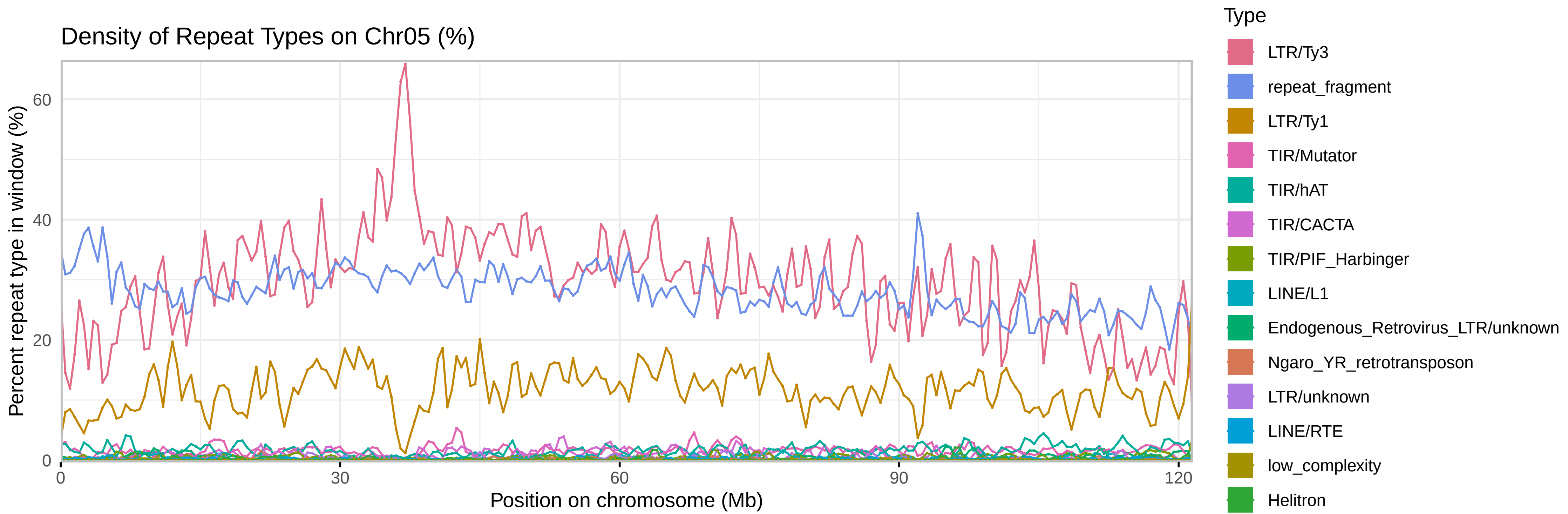

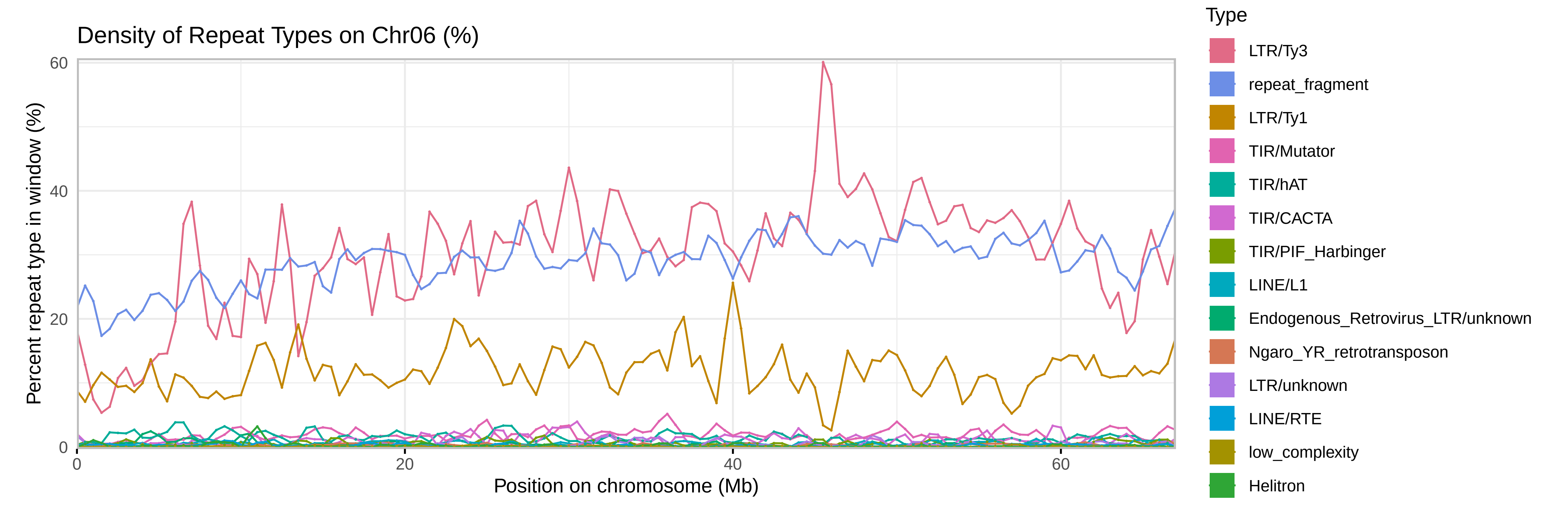

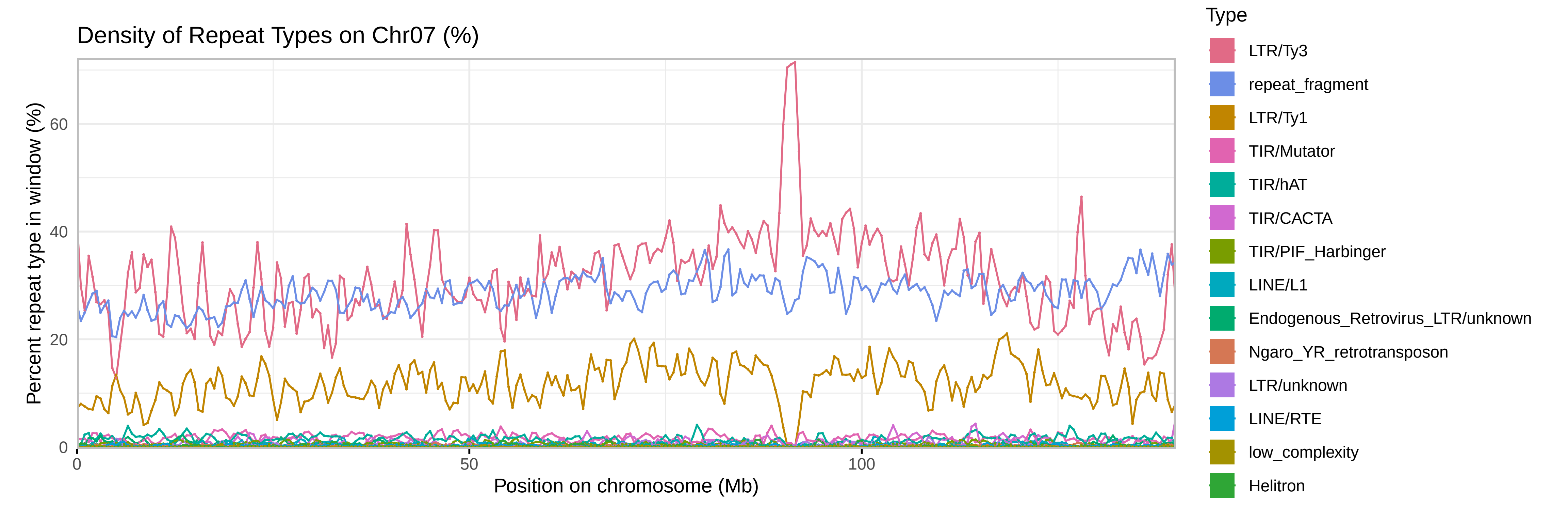

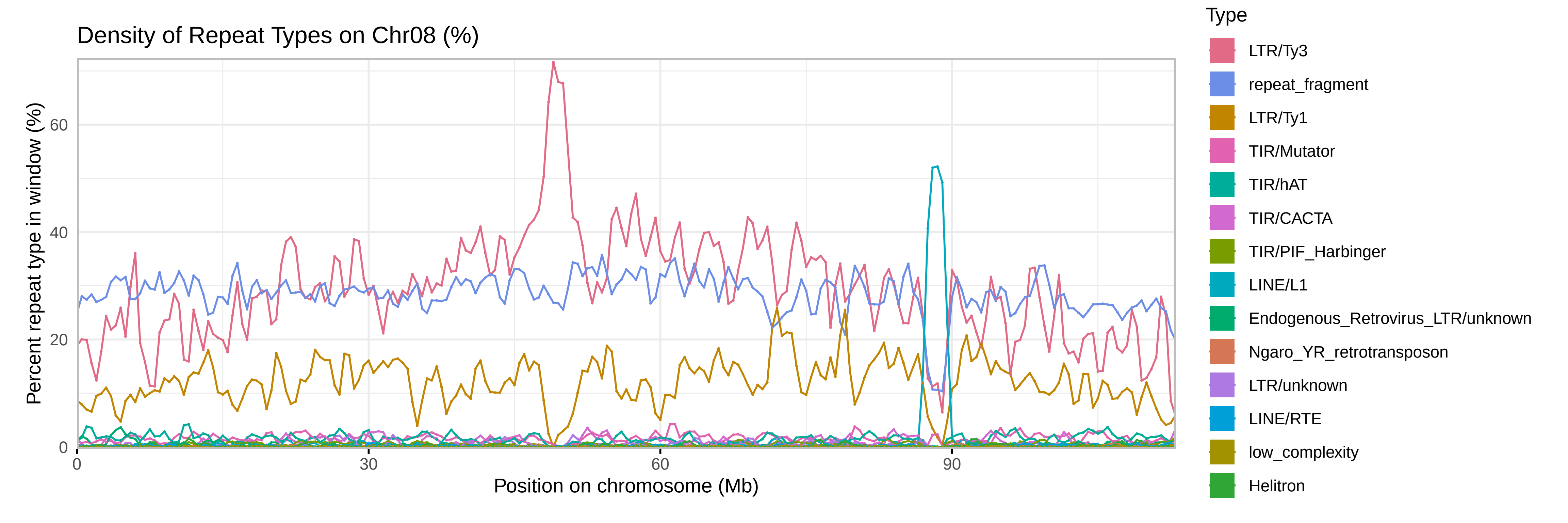

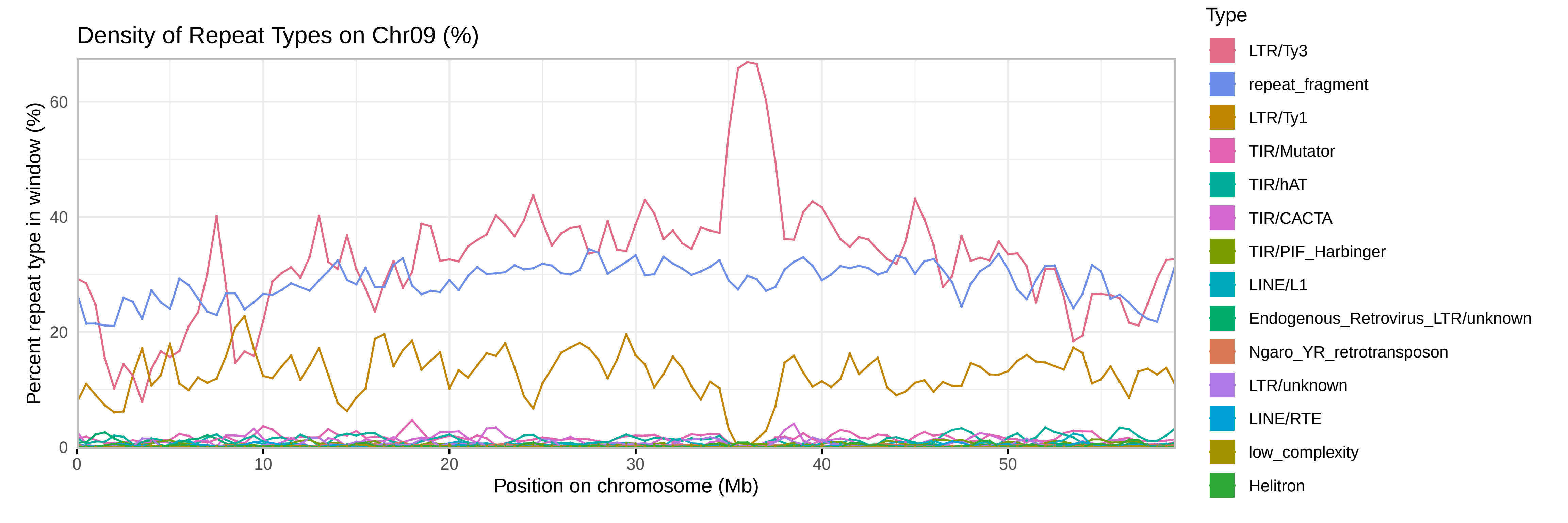

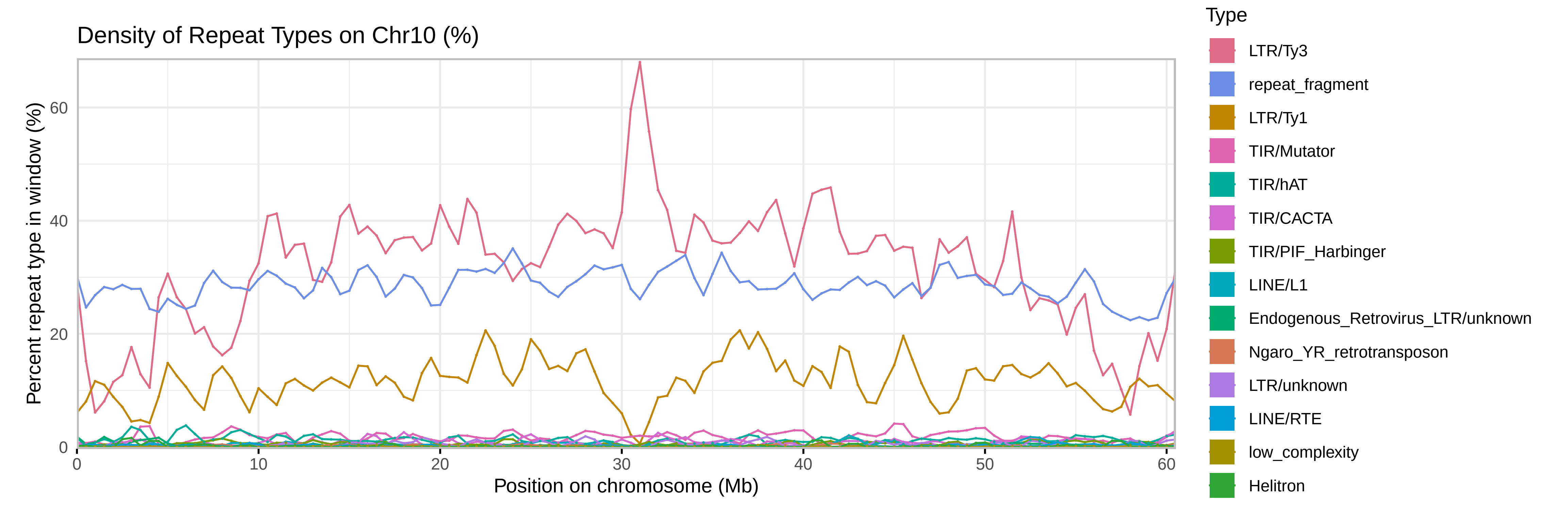

### SupplementalFile9

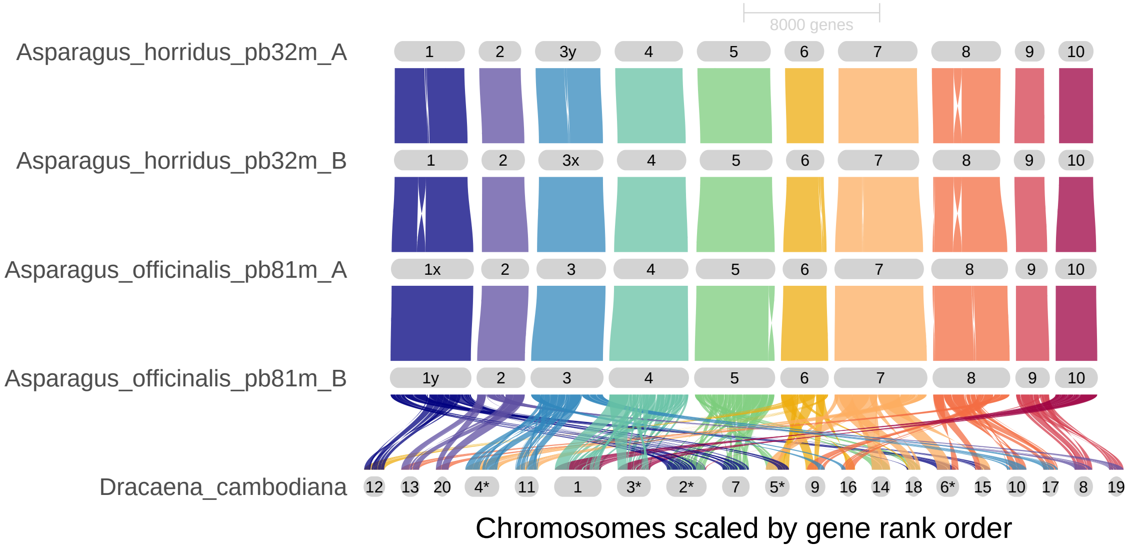

### SupplementalFile10

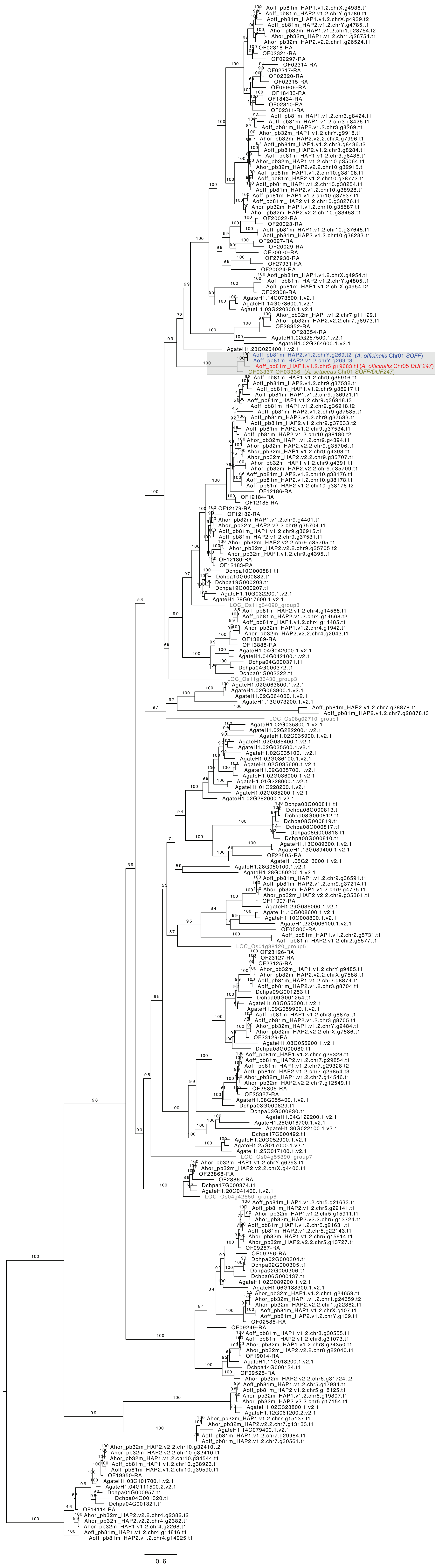
